# Brassinosteroid gene regulatory networks at cellular resolution

**DOI:** 10.1101/2022.09.16.508001

**Authors:** Trevor M Nolan, Nemanja Vukašinović, Che-Wei Hsu, Jingyuan Zhang, Isabelle Vanhoutte, Rachel Shahan, Isaiah W Taylor, Laura Greenstreet, Matthieu Heitz, Anton Afanassiev, Ping Wang, Pablo Szekely, Aiden Brosnan, Yanhai Yin, Geoffrey Schiebinger, Uwe Ohler, Eugenia Russinova, Philip N Benfey

## Abstract

Brassinosteroids (BRs) are plant steroid hormones that regulate diverse processes such as cell division and cell elongation. BRs control thousands of genes through gene regulatory networks that vary in space and time. By using time-series single-cell RNA-sequencing to identify BR-responsive gene expression specific to different cell types and developmental stages of the Arabidopsis root, we uncovered the elongating cortex as a site where BRs trigger a shift from proliferation to elongation associated with increased expression of cell wall-related genes. Our analysis revealed HAT7 and GTL1 as BR-responsive transcription factors that regulate cortex cell elongation. These results establish the cortex as an important site for BR-mediated growth and unveil a BR signaling network regulating the transition from proliferation to elongation, illuminating new aspects of spatiotemporal hormone response.

## Introduction

During development, cells pass through different states as they acquire identities and progress towards end-stage differentiation (*1*). Gene regulatory networks (GRNs) control this progression and must be tuned according to developmental stage, cell identity, and environmental conditions (*2*, *3*). Signaling molecules such as hormones are central players in co-ordinating these networks, but it has been challenging to disentangle how cell identities, developmental states, and hormone responses influence one another. Recent technological advances in single-cell RNA-sequencing (scRNA-seq) (*2*, *4*) and tissue-specific gene manipulations (*5*) make it possible to address this challenge using the *Arabidopsis* root as a model system.

Brassinosteroids (BRs) are a group of plant steroid hormones that affect cell division and cell elongation during root growth (*6–8*). BRs are sensed at the plasma membrane by BRI1 family receptors (*9–11*), initiating signal transduction events that activate BES1 and BZR1 family transcription factors (TFs) to control thousands of genes (*12–15*). The BR GRN is typically represented singularly without consideration of cell specificity (*15*, *16*), but BRs lead to different responses depending on the developmental context (*17–19*).

By profiling BR responses across the cell types and developmental stages of the root using scRNA-seq, we discovered that BRs strongly affect gene expression in the elongating cortex. Reconstruction of cortex trajectories over a scRNA-seq time course showed that BRs trigger a shift from proliferation to elongation, which is associated with up-regulation of cell wall-related genes. Loss of BR signaling in the cortex using tissue-specific CRISPR reduced cell elongation. Our time-course data allowed us to infer BR-responsive GRNs, which led to the identification of HAT7 and GTL1 as validated regulators of BR response in the elongating cortex. These datasets represent more than 180,000 single-cell transcriptomes, providing a view of BR-mediated GRNs at unprecedented resolution.

## Results

### scRNA-seq reveals differential brassinosteroid response in the Arabidopsis root

To investigate spatiotemporal BR responses in the root, we used a sensitized system, which involved inhibiting BR biosynthesis using Brassinazole (BRZ) (*20*), then reactivating signaling with Brassinolide (BL), the most active BR (*8*, *16*) (fig. S1). We treated 7-day-old primary roots for 2 hours with BL or a corresponding mock BRZ control and performed scRNA-seq on protoplasts isolated from three biological replicates of 0.5cm root tips (containing meristem, elongation and early differentiation zones) using the 10X Genomics Chromium system (Methods).

To annotate cell types and developmental stages, we performed label transfer based on our single-cell atlas of the Arabidopsis root (*21*). We distinguished between two domains of the meristem: the proliferation domain, where cells have a high probability to divide, and the transition domain, where cells divide less frequently but have not yet begun rapid expansion (fig. S2A-D and Data S2) (*22*).

After data integration, the 11 major cell types and eight developmental stages identified were logically arranged in 2D uniform manifold approximation and projection (UMAP) space as previously described for root datasets (fig. S3A-B) (*21*, *23*). Marker genes characteristic of cell types and developmental stages remained enriched, suggesting that although BRs alter the expression of thousands of genes, cell identities can be successfully aligned through integration (fig. S3C).

### scRNA-seq captures spatiotemporal patterns of BR-responsive gene expression

Previous studies have profiled BR-responsive gene expression in bulk tissue or in a handful of cell types, conflating cell type and developmental stage (*24–26*). To obtain better spatiotemporal resolution, we performed differential expression analysis for each combination of cell type and developmental stage using pseudobulk expression profiles (see methods). We identified over 8,000 differentially expressed genes (DEGs; Fold-change >1.5, False discovery rate <0.05; Fig. 1A and fig. S4A-B), which were enriched in BES1 and BZR1 targets and had significant overlap with previously identified BR-regulated genes (fig. S4C and Data S3).

**Fig. 1.**
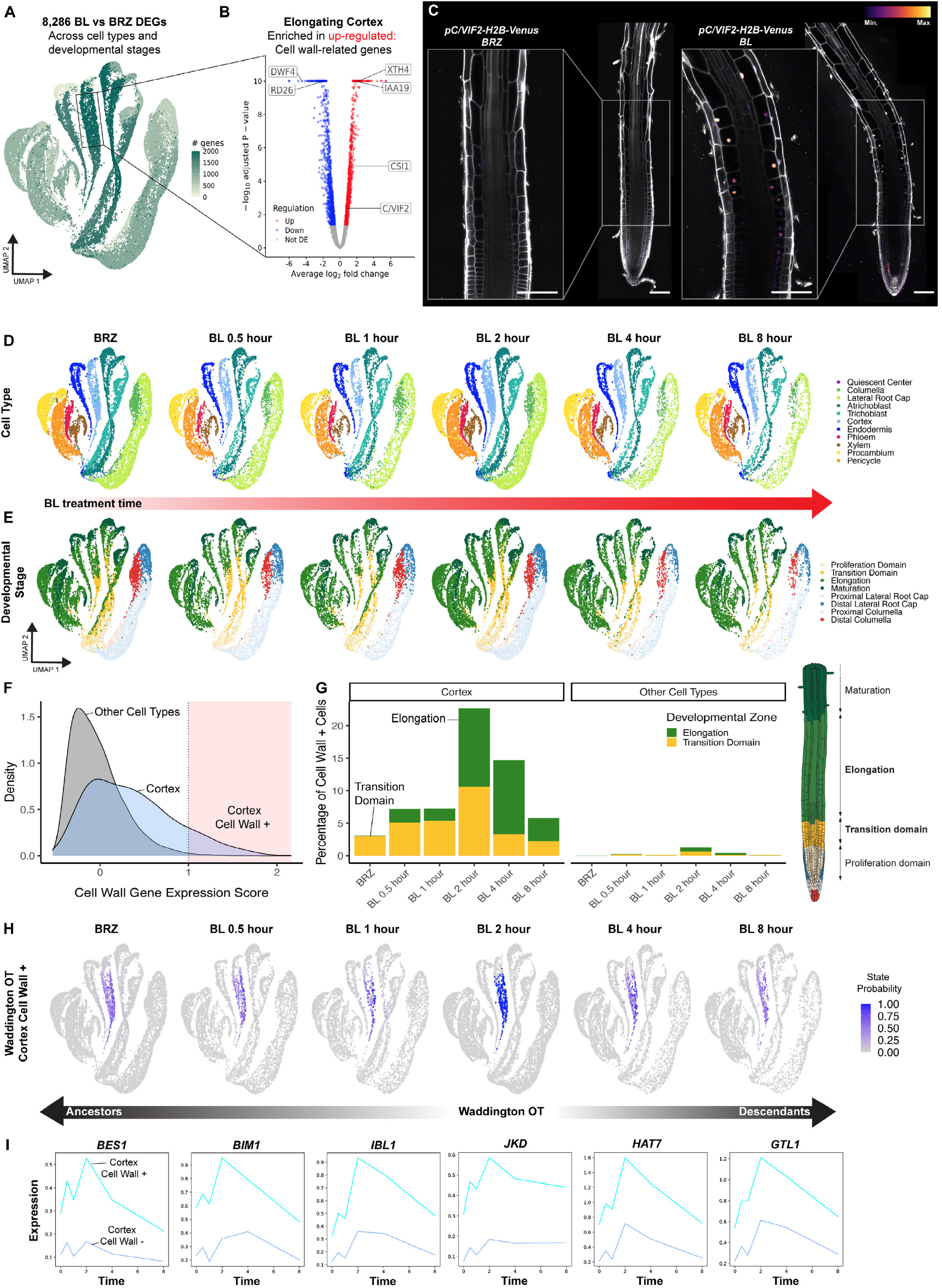
scRNA-seq identifies BR-induction of cell wall-related genes in the cortex associated with the switch to elongation. **(A)** Spatiotemporal response to 2 hour BL treatment vs BRZ control among each combination of cell type and developmental stage of the root. Color on UMAP projection indicates the number of differentially expressed genes (DEGs). **(B)** Volcano plot of BL DEGs in the elongating cortex. Color indicates the direction of regulation. Known markers of BR response including *DWF4, RD26, XTH4* and *IAA19* are indicated. *C/VIF2* and *CSI1* (described in this study) are also indicated. **(C)** *pC/VIF2-H2B-Venus* reporter grown on 1 μM BRZ for 7 days and transferred to 1 μM BRZ or 100 nM BL for 4 hours. Inset shows *C/VIF2* signals in the elongating cortex that increase with BL treatment. Propidium iodide-staining is shown in grey, with the color gradient indicating relative *C/VIF2-H2B-Venus* levels. Scale bars, 100 μm. **(D)** UMAP of 52,921 cells across 6 time points of a scRNA-seq BL treatment time course. Mock BRZ control represents time 0. Colors indicate cell type **(E)** or developmental stage annotation **(F)**. Density plot showing cell wall gene expression score. The shaded region with cell wall expression scores >1 indicates “Cortex Cell Wall+” cells. **(G)** Bar plot showing the percentage of Cell Wall+ cells in the cortex versus other cell types over the time course. Color indicates developmental stage annotation, also depicted in root schematic. Illustration adapted from the Plant Illustrations repository. Only transition and elongation zones are plotted as other zones represent less than 2% of cell wall + cells. **(H)** WaddingtonOT (WOT) probabilities for cortex cell wall+ state along the BL time course. The BL 2 hour time point was used as a reference, therefore all cells have a probability of either 1 or 0 at this time point. **(I)** Expression trends for select TFs differentially expressed along WOT cortex cell wall+ trajectories.

Strikingly, we found that 37% of DEGs were significantly altered in a single cell type/developmental stage and more than 82% were differentially expressed in 5 or fewer cell type/developmental stage combinations (fig. S4B). This indicates that although BRs broadly influence gene expression, they modulate distinct sets of genes in different spatiotemporal contexts.

Among the tissues with many DEGs was the epidermis, as previously described (*7*, *17*, *24*, *25*, *27*). Atrichoblasts, or non-hair cells in the epidermis were particularly affected, showing marked changes across both the meristem and elongation zone. Unexpectedly, our data also indicated that BRs strongly influence gene expression in the cortex, especially in the elongation zone (Fig. 1A and fig. S4A). The cortex has been linked to plant environmental interactions, including response to water limitation (*28*) and hydrotropism (*29*), but it is not known how BRs modulate gene expression in this tissue nor what processes are affected. To address these questions, we focused on BR-mediated gene expression in the elongating cortex.

### Cell wall-related genes are up-regulated by BRs in the elongating cortex

We found that BR treatment led to approximately 1,000 up-regulated genes and about the same number of down-regulated genes in the elongating cortex (Fig. 1B and fig. S4A). Gene ontology (GO) analysis indicated the up-regulated genes were strongly enriched for genes related to “cell wall organization or biogenesis”, which is intriguing given the role of BRs in promoting cell elongation (fig. S4D). The cell wall-related DEGs included *CELLULOSE SYNTHASES (CESAs)*, *CELLULOSE SYNTHASE INTER-ACTIVE1 (CSI1)*, and cell-wall loosening enzymes such as *EXPANSINS* and *XYLOGLUCAN ENDOTRANSGLUCOSYLASES*. Cell-wall-related genes such as *CESAs* have been demonstrated to be direct targets of BES1 and BZR1 (*14*, *15*, *30*), but their spatiotemporal regulation, especially in the cortex, has not been reported.

To monitor their responsiveness to BRs, we generated transcriptional reporters for three of the DEGs with distinct spatiotemporal patterns (Fig. 1C and fig. S5A-E). For example, *CELL WALL / VACUOLAR INHIBITOR OF FRUCTOSIDASE 2 (C/VIF2)* was enriched in the transition domain and elongation zone of the cortex and induced by BL (Fig. 1C). These results confirm that our differential expression analysis captures spatiotemporal BR responses and raise the possibility that BR induction of cell-wall-related genes is associated with cortex cell elongation.

### Waddington optimal transport identifies BR induction of cell wall genes in the cortex associated with the switch to elongation

A recently developed analytical approach for analyzing expression trends over a scRNA-seq time course is Waddington-OT (WOT), which connects snapshots of gene expression to facilitate trajectory reconstruction (*31*). WOT identifies putative ancestors for a given set of cells at earlier time points and descendants at later time points (*31*).

To better understand how BRs influence cell wall-related gene expression we performed scRNA-seq at six time points beginning with BRZ treatment (time 0) and BL treatments for 30 minutes, 1 hour, 2 hours, 4 hours or 8 hours (Fig. 1, D and E). These time points capture the rapid root elongation triggered by re-addition of BRs (*24*).

To examine the trajectories leading to activation of cell wall-related genes in the elongating cortex, we applied WOT (*31*) and created a cell wall gene signature using 107 cell wall-related genes that were induced by BL in the elongating cortex (Data S4). We monitored the relative expression of these genes, resulting in a “cell wall score” for each cell in the time course (see methods). Cortex cells had a higher cell wall score compared to other cell types, which increased with BL treatment (Fig. 1F-G and fig. S6A-B), confirming that the cell wall score represents a BR-responsive module in the cortex. At the 2 hour BL time point, more than 20% of cortex cells had a cell wall score greater than 1, whereas only 5% or fewer cells in other cell types exhibited scores this high (Fig. 1F). We therefore designated cells with a cell wall score of at least 1 as “cortex cell wall+” to indicate their exceptional BR response (Fig. 1, F and G).

An advantage of WOT analysis is that it does not rely on pre-specified boundaries between developmental zones. We used this property to examine the relationship between developmental stage annotation and cell wall score. Under BRZ treatment, cortex cell wall+ cells were sparse and predominately annotated as transition domain. Upon BL treatment, the annotation of cortex cell wall+ cells shifted to the elongation zone (Fig. 1G). These results suggest that BRs are involved in initiating elongation of cortex cells via activation of cell wall genes.

### Differential expression along WOT trajectories identifies BR-responsive transcription factors

Using the cells at the 2 hour time point as a reference, we looked at the probability of cells being ancestors or descendants of cortex cell wall+ cells (Fig. 1H). We also constructed a similar trajectory for the remaining cortex cells, which were designated “cortex cell wall-”. To reveal potential regulators of cell wall-related genes in the cortex, we performed probabilistic differential expression analysis along WOT trajectories, contrasting cells assigned to cortex cell wall+ versus cortex cell wall- states at each time point (see methods). Among the DEGs identified were known TFs in the BR pathway including *BES1* (*12*), *BIM1* (*32*), and *IBL1* (*33*); Fig. 1I and Data S4). We also identified additional TFs including *JACKDAW* (*JKD*), which is involved in ground tissue specification (*34*), the class I HD-ZIP TF *HAT7* (*35*, *36*) and *GTL1*. Because *JKD* was uniquely identified in this analysis, we confirmed its BR-responsiveness (Figures S6, C and D). These results indicate that WOT trajectories can identify BR-responsive TFs that may be involved in regulating cell wall-related genes in the cortex.

### Analysis of the triple receptor mutant bri1-T reveals changes in cortex expression

Since our results indicated that exogenous BRs lead to activation of cell wall-related genes in the elongating cortex, we asked if this is also the case for endogenous BRs. A gradient of BRs is present along the longitudinal axis of the root, with low BR levels in the proliferation domain (*37*). BR biosynthesis increases as cells enter the transition domain and peaks in the elongation zone, shootward of which is a BR signaling maximum (*24*, *37*). Interpretation of this endogenous BR gradient requires receptor BRI1 and its close homologs BRL1 and BRL3 (*9*, *38*, *39*).

To identify differentially expressed genes, we performed two replicates of scRNA-seq on the BR-blind *bri1brl1brl3* triple mutant (*bri1-T)* along with paired wild-type controls (Fig. 2A-C). A previous study profiled single cells from *bri1-T*, suggesting potential BR-responsiveness of the cortex (*40*), but these data were from a single replicate, were compared to a wild-type sample from a different study, and could not resolve developmental stage-specificity. In contrast, our analysis across both cell types and developmental stages identified the elongating cortex as exhibiting substantial differential gene expression (Fig. 2, D and E, fig. S7A and Data S3). The genes down-regulated in the elongating cortex of *bri1-T* were enriched for the GO term “cell wall organization or biogenesis” (Figure S7B). These data indicate that endogenous BR signaling promotes the expression of cell wall-related genes in the elongating cortex.

**Fig. 2.**
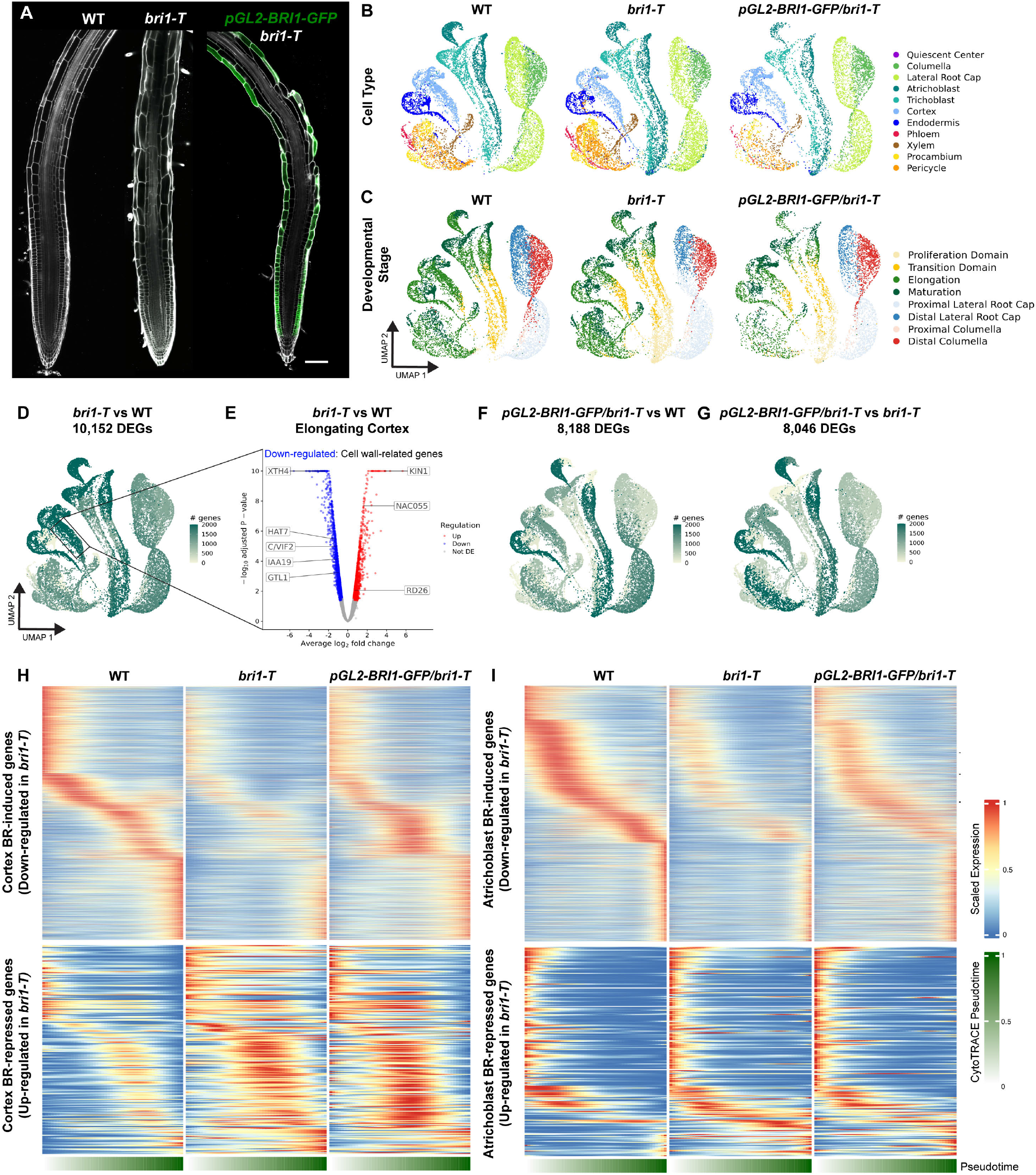
Triple receptor mutant *bri1-T* gene expression changes in cortex and distinct patterns in *pGL2-BRI1-GFP/bri1-T*. **(A)** 7-day old WT, *bri1-T* and *pGL2-BRI1-GFP/bri1-T* roots grown under control conditions. Propidium iodide-staining is shown in grey and GFP in green. Scale bars, 100 μm. **(B)** UMAP projection of scRNA-seq from 14,334 wild-type cells, 12,649 *bri1-T* cells and 7,878 *pGL2-BRI1-GFP/bri1-T* cells. Two biological replicates of scRNA-seq were performed for each genotype. Colors indicate cell type annotation. **(C)** UMAP projection colored by developmental stage annotation. **(D)** UMAP colored by DEGs for each cell type/developmental stage combination of *bri1-T* compared to WT. **(E)** Volcano plot of DEGs in the elongating cortex from *bri1-T* compared to WT. Color indicates the direction of regulation. **(F-G)** UMAP colored by DEGs for each cell type/developmental stage combination of *pGL2-BRI1-GFP/bri1-T* compared to WT (F) or *pGL2:BRI1-GFP/bri1-T* compared to *bri1-T* (G). **(H-I)** Gene expression trends for *bri1-T* vs wild-type DEGs along cortex (H) or atrichoblast (I) trajectories. Scaled expression along cortex pseudotime is plotted for each genotype. Lower bar indicates pseudotime progression calculated by CytoTRACE.

The epidermis is widely described as the major site for BR-promoted gene expression in the root (*7*, *8*, *17*, *24*, *25*, *41*). Previous studies showed that epidermal expression of BRI1 was sufficient to rescue morphological phenotypes including meristem size of *bri1-T* (*7*, *17*, *42*). To determine the extent to which BR-regulated gene expression is restored, we performed scRNA-seq on *pGL2-BRI1-GFP/bri1-T* - a line in which BRI1 is expressed in atrichoblast cells of the epidermis of *bri1-T (7, 25, 42)*. We identified over 8,000 DEGs in comparison with wild type (Fig. 2F) and in comparison with *bri1-T* (Fig. 2G-I and fig. S8A-F), indicating that gene expression remains dramatically perturbed and that this is far from a complete rescue of the *bri1-T phenotype*.

### Tissue-specific CRISPR of BRI1 confirms a role for the cortex in BR-mediated cell expansion

Characterization of cell-type-specific BR signaling has relied on tissue-specific complementation lines, which led to conflicting results and overlooked the role of BR signaling in the cortex (*7*, *17*, *24*, *25*, *40*, *42*, *43*). To selectively block BR signaling in cell types of interest we performed tissue-specific CRISPR (*5*) of *BRI1*. We used a *bri1* mutant complemented with pBRI1-BRI1-mCitrine (Fig. 3A) into which we introduced Cas9 driven by tissue-specific promoters to knock out BRI1 either in the epidermis and lateral root cap (pWER-BRI1-CRISPR) or in the cortex (pCO2-BRI1-CRISPR). mCitrine signals were absent in the expected locations of the tissue-specific CRISPR lines, confirming their efficacy and specificity (Fig. 3A-C and fig. S9A-C).

**Fig. 3.**
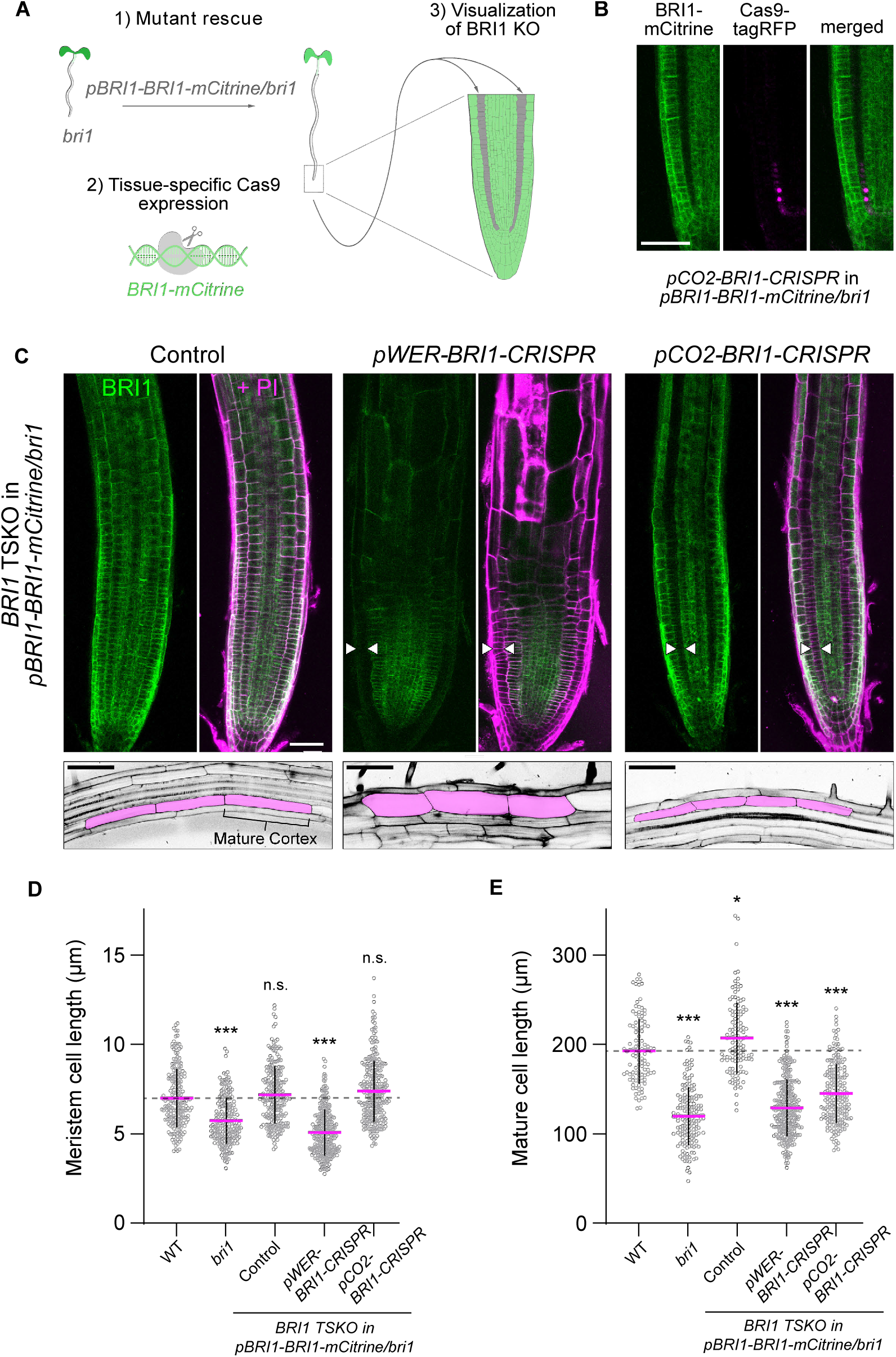
Tissue-specific CRISPR of BRI1 confirms role for cortex in BR-mediated cell expansion. **(A)** Overview of BRI1 tissue-specific CRISPR approach. A *bri1* mutant complemented with pBRI1-BRI1-mCitrine (1) was used as background to introduce tissue-specific Cas9 along with gRNAs targeting BRI1 (2). This allows for visualization of BRI1 knockout in specific cell layers, such as the cortex when pCO2-BRI1-CRISPR is used (3). **(B)** Appearance of Cas9-tagRFP in the cortex is associated with loss of BRI1-mCitrine signal, confirming tissue-specific knockout. **(C)** Confocal images of BRI1 tissue-specific CRISPR lines. Control indicates a broad expression pattern of BRI1-mCitrine in pBRI1-BRI1-mCitrine/*bri1*. BRI1-mCitrine signals are shown in green and propidium iodide staining (PI) in magenta (upper panels). White arrows specify tissues with absence of BRI1-mCitrine signal; epidermis for pWER-BRI1-CRISPR and cortex for pCO2-BRI1-CRISPR. Mature root sections illustrating changes in cell size and length (lower panels). Cortex cells are pseudocolored to indicate their position. **(D)** Quantification of meristematic cortex cell length, defined as the first 20cells of individual roots starting from the quiescent center. Control indicates pBRI1-BRI1-mCitrine/*bri1* complemented line. **(E)** Quantification of mature cortex cell length. For (D) and (E), all individual data points are plotted. Magenta horizontal bars represent the means and error bars represent s.d. Significant differences between each line and wild type were determined by one-way ANOVA and Dunnett’s multiple comparison tests. *** P<0.001, ** P<0.01 and * P<0.05. n.s. not significant. Scale bars, 50 μm. TSKO, tissue-specific knockout.

Since our scRNA-seq data indicated that BRs promote the expression of cell wall-related genes in the elongating cortex, we hypothesized that loss of BR signaling in the cortex would affect final cell size. Indeed, pCO2-BRI1-CRISPR lines displayed significantly shorter mature cortex cells, while meristematic cortex cell length was relatively unaffected (Fig. 3C-E).

In contrast, epidermal knockout of BRI1 in pWER-BRI1-CRISPR lines resulted in both reduced meristem cell size and reduced mature cortex cell length (Fig. 3C-E), which is consistent with the reported role of epidermal BR signaling (*7*, *18*, *24*, *25*, *41*) and BR-responsiveness across developmental zones of the epidermis in our scRNA-seq data. These results indicate that in addition to the epidermis, BR signaling in the cortex is required to promote cell expansion in the elongation zone. The cortex could instruct anisotropic growth through its physical connection with the epidermis, but as the outermost tissue, relaxation of the epidermis appears to be required to allow for cell elongation (*44*, *45*). This may explain the apparent widening of cortex cells in pWER-BRI1-CRISPR lines.

BRI1 was also reported to rescue *bri1-T* morphology when expressed in the developing phloem using the *CVP2* promoter (*40*, *42*, *43*), but gene expression was not fully restored to wild-type levels in either epidermal or phloem rescue lines. Our scRNA-seq of epidermal *pGL2-BRI1-GFP/bri1-T* lines showed patterns of gene expression distinct from either wild type or *bri1-T*. Similarly, scRNA-seq of *pCVP2-BRI1-CITRINE/bri1-T* indicated an intermediate state between wild type and *bri1-T (40)*. Notably, BRI1 driven by its native promoter was still present in the stele of our tissue-specific CRISPR lines when we observed phenotypic defects, suggesting that, unlike *pCVP2-BRI1*, native expression of *BRI1* in the stele is not sufficient for BR-induced cell elongation and root growth. These results confirm the role of the epidermis in BR-regulated root growth and reveal the function of cortex in BR-mediated cell expansion, demonstrating how scRNA-seq can identify a new spatiotemporal context for hormone signaling.

### HAT7 and GTL1 are BR-responsive regulators along cortex trajectories

To define a core set of genes associated with BR response along cortex trajectories we first compared genes induced in the cortex by BL treatment with those down-regulated in the cortex of *bri1-T*. Of the 768 genes in common, we then asked which vary along developmental time in wild-type cortex trajectories (*21*). The intersection of these three lists identified a core set of 163 BR responsive DEGs (Fig. 4A and Data S3). Consistent with regulation by BRs, 69% of the core DEGs are BES1 and BZR1 direct targets from ChIP experiments (*14*, *15*, *46*). Expression along cortex pseudotime illustrates induction by BL treatment and down-regulation in *bri1-T* (Fig. 4B). *HAT7* and *GTL1* were induced along these trajectories, suggesting a potential role for these TFs in controlling BR-regulated gene expression in the cortex (Fig. 4C-E and fig. S10A).

**Fig. 4.**
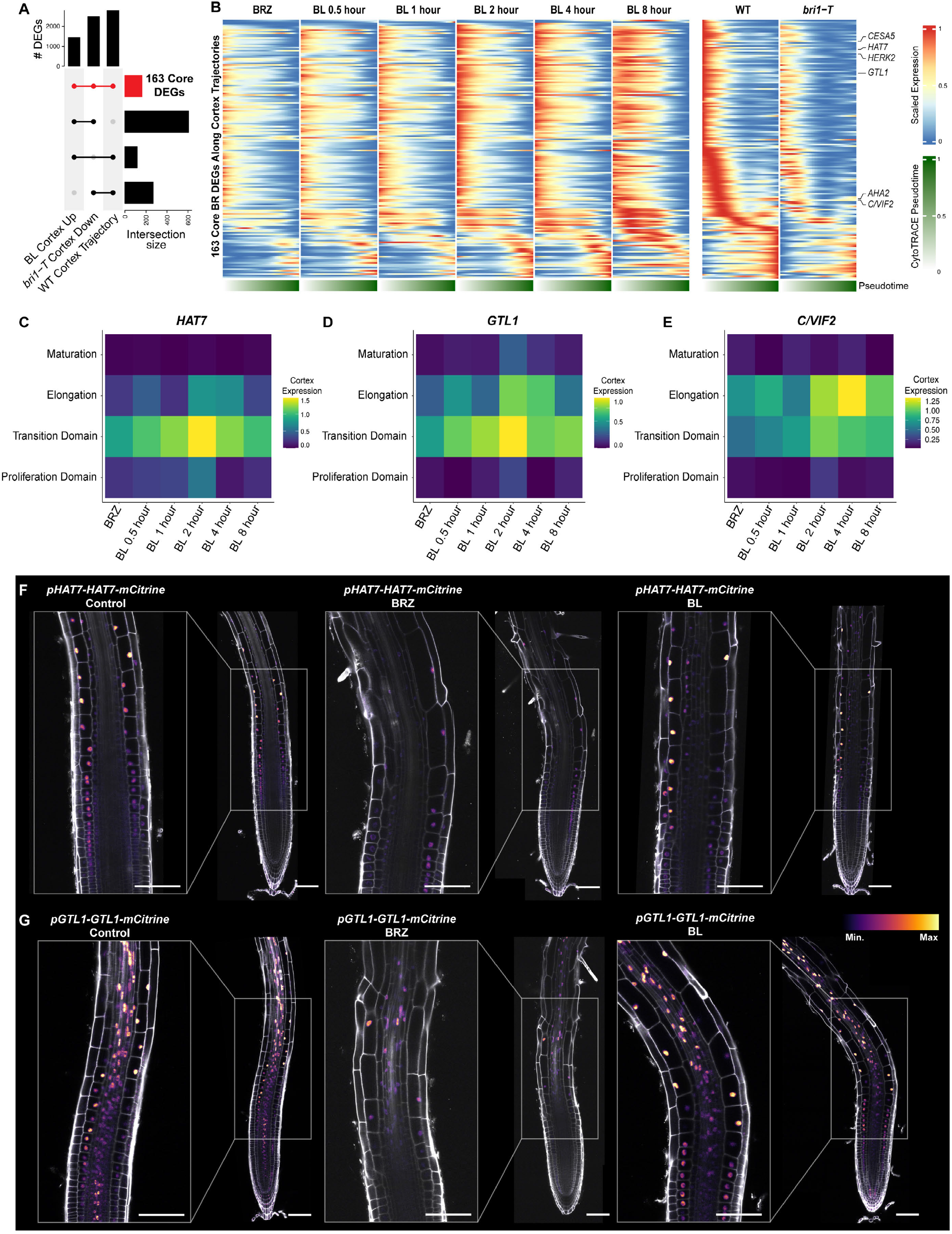
*HAT7* **and** *GTL1* are BR responsive regulators along cortex trajectories. **(A)** Upset plot showing a comparison of genes up-regulated by BL in the cortex, down-regulated in the cortex of *bri1-T*, and differentially expressed along wild-type cortex trajectories. Red color indicates 163 genes common to all three sets. **(B)** Gene expression trends for 163 core BR DEGs along cortex trajectories. Scaled expression along cortex pseudotime is plotted for each time point of the BR time series and for wild type versus *bri1-T*. Lower bar indicates pseudotime progression calculated by CytoTRACE. **(C-E)** Gene expression trends for *HAT7, GTL1* or *C/VIF2* along the developmental zones of the cortex (y-axis) for each time point of the BR time course (x-axis). Color bar indicates the scaled expression level in the cortex. **(F-G)** 7-day old roots expressing *pHAT7-HAT7-mCitrine* (F) or *pGTL1-GTL1-mCitrine* (G) reporters under the indicated treatments. Control represents a mock DMSO solvent. For BRZ and BL treatments, plants were grown on 1 μM BRZ for 7 days and transferred to 1 μM BRZ or 100nM BL for 4 hours. Propidium iodide-staining is shown in grey, with the color gradient indicating relative mCitrine levels. Scale bars, 100 μm.

To gain insight into their roles, we generated translational reporter lines for HAT7 and GTL1. Under control conditions, *pHAT7-HAT7-mCitrine* lines showed expression in the transition domain and elongation zone of the cortex (Fig. 4F). We also observed HAT7 signals in the epidermis and endodermis, in line with expression patterns in our wild-type scRNA-seq atlas (*21*). HAT7 expression was decreased when BR biosynthesis was inhibited with BRZ, and restored upon BL treatment (Fig. 4F and fig. S10A).

*pGTL1-GTL1-mCitrine* was more broadly expressed, with increasing levels in the cortex and epidermis as cells progress from the transition domain to the elongation zone (Fig. 4G and S10A). GTL1-mCitrine expression was reduced by BRZ and increased by BL treatment (Figure 4G). These results confirm that BRs promote the expression of HAT7 and GTL1 coinciding with the onset of cell elongation. Furthermore, HAT7 and GTL1 are direct targets of BES1 and BZR1 (*14*, *15*, *46*), suggesting that they may be part of the BR-directed GRN activated as cells progress from proliferation to elongation.

Previous studies inferred global (*14*, *15*, *26*, *47*) or temporally resolved GRNs (*16*) for BR response, but they lacked cell type and developmental-stage specificity. To infer GRN configurations across our BR time series we used CellOracle (see methods; Data S8) focusing on BL DEGs and associated TFs.

Analysis of network importance scores such as centrality measures is a powerful approach to prioritize candidate regulators among DEGs (*48*). Since the cell wall signature peaked at 2 hours after BL treatment, we prioritized TFs with high network centrality scores in the elongating cortex at this time point. HAT7 was the top-ranked TF in terms of degree centrality and three close homologs: HB13, HB20 and HB23 were also among the top 10 TFs (Fig. 5A-B and Data S8). Together HAT7, HB13, HB20 and HB23 make up the alpha clade HD-ZIP I TFs (*35*, *36*).

**Fig. 5.**
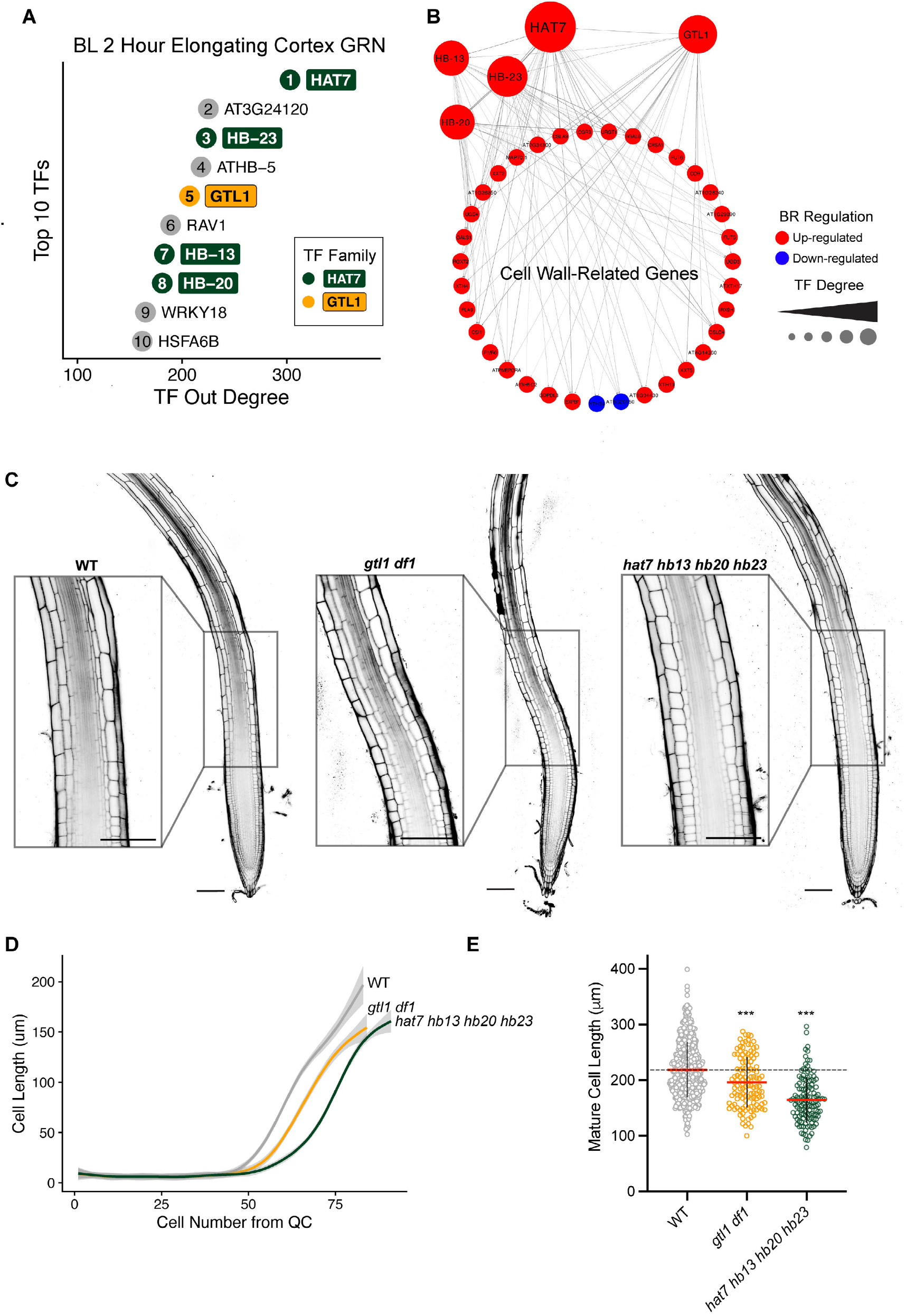
HAT7 and GTL1 are top-ranked regulators in cortex GRNs and affect BR-related phenotypes. **(A)** Top 10 TFs in the CellOracle BL two-hour elongating cortex GRN ranked by out degree. The ranking is indicated by the number inside the circle. Color indicates TF family, with light grey corresponding to any family other than HAT7 or GTL1. **(B)** Subnetwork showing cell wall-related genes that are predicted targets of HAT7 and GTL1 in the CellOracle elongating cortex GRN. HB13, HB20 and HB23 are included in the subnetwork since they are connected to HAT7 and cell-wall-related genes. **(C)** Propidium iodide-staining of 7-day-old WT, *hat7 hb13 hb20 hb23* (line 1-2), and *gtl1 df1* roots. Insets show cortex cells entering the elongation zone. Scale bars, 100 μm. **(D)** Quantification of cortex cell length along the longitudinal axis of the root. The quiescent center was designated as “0” and each cell number consecutively thereafter. The grey area represents the confidence interval of the smoothed mean estimated with a generalized additive model. Number of roots per genotype: WT=51, *gtl1 df1*=26, *hat7 hb13 hb20 hb23*=16. **(E)** Quantification of mature cortex cell length. Red horizontal bars represent the means and error bars represent s.d. Significant differences between each line and wild type were determined by one-way ANOVA and Dunnett’s multiple comparison tests. ***P<0.001.

We used CRISPR to generate *hat7* loss-of-function mutants but did not observe strong phenotypes in terms of cortex cell elongation (Fig. S11A-C). Since *HB13*, *HB20* and *HB23* are induced by BRs and predicted to regulate cell wall-related genes in our GRNs (Fig. 5B, fig. S10A-B and Data S10), we next generated *hat7 hb13 hb20 hb23* quadruple mutants via multiplex CRISPR. Mature cortex cell length was reduced by approximately 25% in two independent quadruple mutants (Fig. 5C-E and fig. S11A-C), providing strong evidence that HAT7 and its homologs are required for cell elongation. Despite the decrease in final cell length, the root length of the quadruple mutant was not dramatically reduced (Fig. S11A), suggesting that the decrease in cell length is at least partially compensated for by increased cell production.

We next investigated GTL1, which was the 5th highest-ranked TF in the BL 2 hour elongating cortex GRN (Fig. 5A-B). Given that GTL1 was shown to function redundantly with its close homolog DF1 in terminating root hair growth (*49*), we examined *gtl1 df1* double mutants, finding significantly shorter mature cortex cell lengths and shorter roots compared to wild-type (Fig. 5C-E and fig. S11A-C). *DF1* was challenging to detect in scRNA-seq (fig. S10A) due to its low expression level (*49*), but we observed increasing trends of *DF1* expression along WOT trajectories in the BL time course, especially in cortex cell wall+ cells (fig. S10B) which was verified using a p*DF1-DF1-GFP* reporter (fig. S10C). Together, our genetic analysis of HAT7 and GTL1 family TFs illustrates the power of GRN-mediated discovery of regulatory factors in spatiotemporal BR response.

### BES1 and GTL1 physically interact and share a common set of target genes

Since BES1 is known to interface with other TFs in controlling BR-regulated gene expression, we compared target genes for BES1 and BZR1 (*14*, *15*, *46*) to ChIP targets of GTL1 and DF1 (*49*). BES1 and BZR1 share 3,020 common targets with GTL1, significantly more than expected by chance (fig. S12A, P < 0.001, Fisher’s exact test). Similarly, BES1 and BZR1 share 2,490 common targets with DF1 (Figure S12A, P < 0.001, Fisher’s exact test). When compared to BR-regulated genes from scRNA-seq, BES1 and GTL1 targets showed the strongest enrichment in genes up-regulated by BRs in the transition domain and elongation zone of the cortex (fig. S12B), with 297 common targets of both BES1 or BZR1 and GTL1 being induced in the elongating cortex by BL treatment.

Given the strong overlap between BES1 and GTL1 targets, we hypothesized that these TFs physically interact to regulate a common set of genes. Co-immunoprecipitation showed that GTL1-FLAG pulled down BES1-GFP (fig. S12C). These results suggest that BRs induce GTL1 and subsequently BES1 and GTL1 interact to control a common set of target genes. This type of feed-forward loop could provide a mechanism to amplify the BR signal and/or to direct BES1 to drive tissue-specific gene expression by interacting with other more specifically expressed TFs.

### scRNA-seq reveals cell-type-specific expression underlying gtl1 df1 phenotypes

Our results indicate that *gtl1 df1* mutants have reduced cortex cell elongation. On the other hand, *gtl1 df1* mutants have longer trichoblasts (*49*). A downstream regulatory network that enables GTL1-mediated growth inhibition has been dissected in trichoblasts (*49*). To identify the cell-type-specific changes in gene expression underlying *gtl1 df1* cortex phenotypes we performed scRNA-seq on *gtl1* and *df1* single mutants, and on the *gtl1 df1* double mutant. Using pseudobulk differential expression analysis, we detected relatively subtle changes in *gtl1* or *df1* single mutants compared to wild type (Fig. 6, A and B). In contrast, over 8,000 genes were differentially expressed in *gtl1 df1* double mutants versus wildtype (Fig. 6C).

**Fig. 6.**
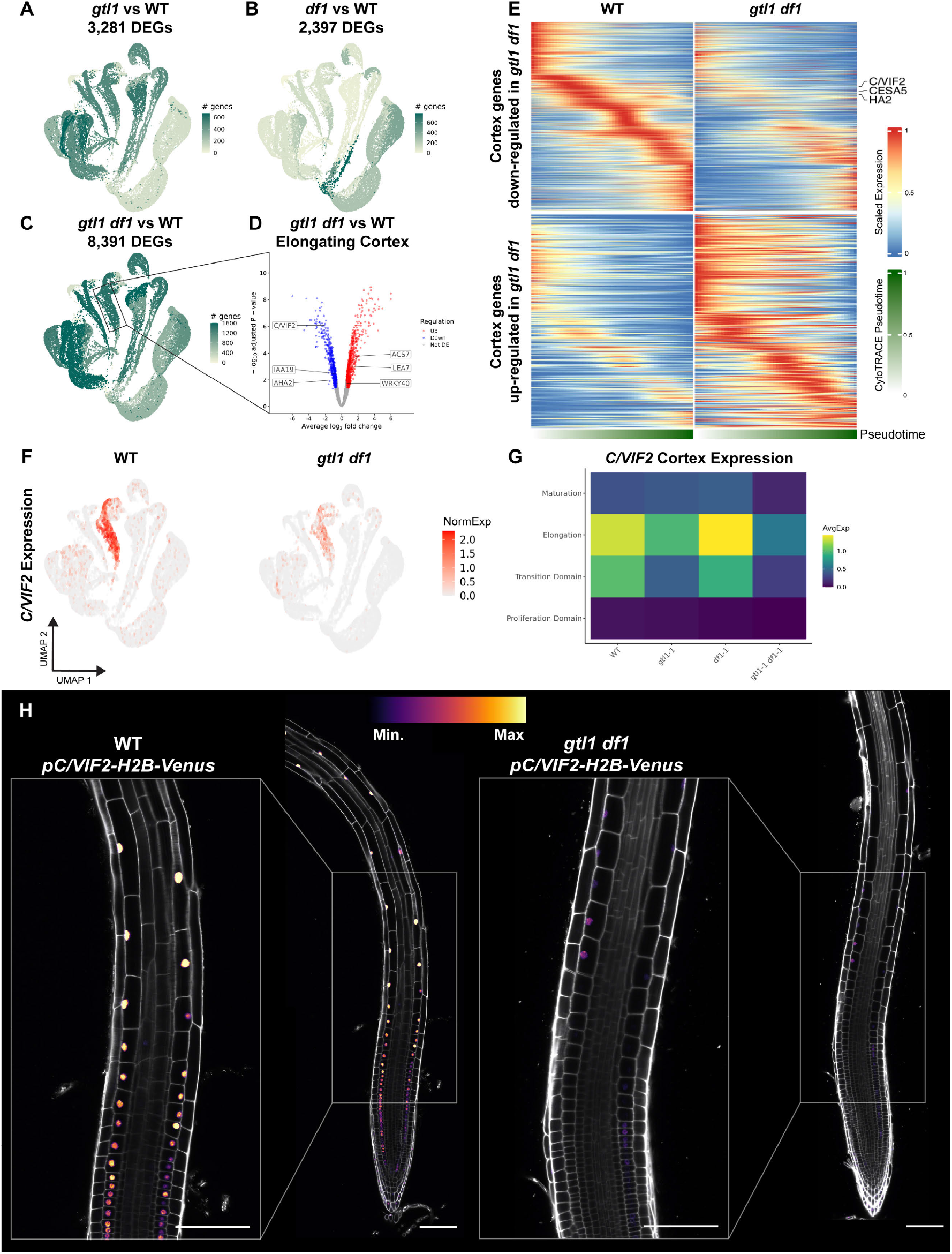
scRNA-seq reveals cell-type-specific expression underlying *gtl1 df1* phenotypes. **(A-C)** UMAP projection of scRNA-seq from 74,810 WT, *gtl1*, *df1*, and *gtl1 df1* cells. Two biological replicates were profiled for each genotype. Color indicates DEGs for each cell type/developmental stage combination of *gtl1* compared to WT (A), *df1* compared to WT (B) or *gtl1 df1* compared to WT (C). **(D)** Volcano plot of DEGs in the elongating cortex from *gtl1 df1* compared to WT. Color indicates the direction of regulation. **(E)** Gene expression trends along cortex trajectories for DEGs in *gtl1 df1* compared to WT. Each row represents the scaled expression of a gene along cortex pseudotime. The lower bar indicates pseudotime progression calculated by CytoTRACE. **(F)** Expression of *C/VIF2* in wild type and *gtl1 df1* scRNA-seq. The color scale represents log normalized, corrected UMI counts. **(G)** Gene expression trends plotted along developmental zones of the cortex (y-axis) for WT, *gtl1*, *df1*, and *gtl1 df1*. The color bar indicates the scaled expression level. **(H)** 7-day old root images of a *pC/VIF2-H2B-Venus* reporter in wild type or *gtl1 df1* under control conditions. Propidium iodide-staining is shown in grey, with the color gradient indicating relative mCitrine levels. Scale bars, 100 μm.

Over 1,000 genes were up-regulated across all developmental stages of the cortex of *gtl1 df1*, and an approximately equal number of genes were down-regulated. The majority of cortex DEGs were affected in the elongation zone (Fig. 6, D and E, fig. S12D and Data S3). Of the down-regulated genes in the cortex of the double mutant, 226 genes were also up-regulated by BL treatment. Furthermore, 31.3% of the core BR DEGs were down-regulated in the cortex of *gtl1 df1*, whereas only 6.8% were up-regulated (Data S3). These results suggest that GTL1 and DF1 promote the expression of a subset of BR-induced genes in the cortex.

Plotting *gtl1 df1* DEGs along cortex pseudotime illustrated the down-regulation of several genes involved in cell elongation including *CESA5* and *AHA2* (Fig. 6E). These genes were significantly enriched for the GO term “cell wall organization or biogenesis” (fig. S12E). We next examined *C/VIF2*, because it is induced by BL in the cortex, but its expression decreased in cortex cells of *gtl1 df1* (Fig. 6, F and G). A *pC/VIF2-H2B-Venus* reporter showed expression of *C/VIF2* in the transition and elongation zone of the wild-type cortex, whereas its expression was reduced in the cortex of *gtl1 df1* mutants (Fig. 6H and Movie S1). The reduced expression of cell wall-related genes in *gtl1 df1* mutants validates our cell-type-specific BR GRNs and identifies a function of GTL1 in promoting cortex cell elongation in response to BRs.

## Discussion

Understanding how hormone-mediated GRNs are controlled in space and time has the potential to enable the engineering of specific downstream responses to optimize plant growth under a changing environment (*8*, *50*). Plant hormones including BRs, auxin, gibberellins, and abscisic acid have been shown to exhibit tissue-specific responses (*51–54*), but how the associated GRNs are modulated in different cell types at particular developmental stages is largely enigmatic. In this study, we profiled BR-responses across cell types, developmental stages and time points of treatment using scRNA-seq, providing a high-resolution map of signaling outputs. These data are publicly available as an interactive browser (https://shiny.mdc-berlin.de/ARVEX/). We identified the elongating cortex as a spatiotemporal context for BR signaling, where BRs activate cell wall-related genes and promote elongation. We further showed that HAT7 and GTL1 are BR-induced regulators along cortex trajectories that control cell elongation. These findings highlight the ability of single-cell genomics to identify context-specific TFs, a capability that could be leveraged to precisely engineer plant growth, development, and/or responses to stress. Our results reveal spatiotemporal BR responses and the underlying GRNs at unprecedented resolution.

## Methods

### Plant materials and growth conditions

Arabidopsis accession Columbia-0 (Col-0) was used as a wild type. The following lines were previously described: *bri1 GABI_134E10 (55); bri1-116brl1brl3* triple mutant (*bri1-T) (56); pGL2-BRI1-GFP/bri1-T (25); gtl1-1* (WiscDsLox413-416C9), *df1-1* (SALK_106258), and *gtl1-1 df1-1 (49)*; JKD-Ypet recombineering line (*34*). Seeds were sterilized using 50% (v/v) bleach with 0.05% Tween-20 for 10-15 minutes, plated on 1/2 Linsmaier and Skoog (LSP03-1LT, Caisson Labs; pH 5.7), 1% sucrose media, and stratified 2-4 days at 4°C in the dark. Plates were kept vertically in a Percival growth chamber set to 22°C, 16 hours light/8 hours dark and grown for 7 days unless otherwise indicated. Chemical treatments were conducted by cooling the growth media to approximately 60°C after autoclaving and adding DMSO (a mock solvent), 1 μM Brassinazole (BRZ, SML1406, Sigma) or 100nM Brassinolide (BL, 21594, Cayman Chemical).

*bri1-T* was maintained as a heterozygote for *bri1-116* and homozygous mutants were confirmed as previously described (*42*). Primers listed in Data S11 were used to amplify genomic DNA and the resulting 552bp amplicon was digested with *PmeI*. The mutant *bri1-116* allele could not be digested, whereas WT was cut into 314bp and 238bp fragments.

### Transgenic reporters

To generate new reporters for BR-responsive genes, we first added the FASTRED seed coat selection cassette (*57*, *58*) and a MoClo (*59*) Level 1 acceptor site to the binary vector pICH86966 (Addgene plasmid #48075). *pHAT7-HAT7-mCitrine* and *pGTL1-GTL1-mCitrine* were assembled into this FASTRED destination vector using Level 1 BsaI golden gate assembly. To facilitate one-step promoter-reporter construction, we assembled an AarI flanked RFP dropout using the overhangs described in the Mobius (*60*) upstream of Venus-H2B followed by the Ubiquitin10 terminator (tUBQ10), a plasma membrane marker (pUBQ10-mScarlet-LTI6-tNos), and a constitutive histone maker (pUBQ10-H2B-CFP-t19s). Promoters containing up to ~3kb of sequence upstream of the ATG start codon for the gene of interest were PCR amilfied with AarI containing primers and used to replace the AarI-RFP module in golden gate reactions to generate Promoter-Venus-H2B constructs. Our Venus-H2B reporter included a Ubiquitin tag to decrease reporter perdurance. Although the plasma membrane marker and histone marker were included as positive controls in the constructs, they were not further analyzed in this study.

Assemblies were confirmed by restriction digestion and sequencing, transformed into *Agrobacterium*, and used to transform Arabidopsis via floral dip (*61*). FASTRED positive T1 seeds were selected under a fluorescent dissecting scope and only lines with 3:1 segregation of seed coat fluorescence in the T2 generation were used. T2 lines with bright seed fluorescence were typically homozygous in our conditions. Therefore, we used bright T2 seeds or homozygous T3 seeds for experiments. We ensured that reporter signals were consistent across at least three independent transgenic lines.

### Generation of mutant lines using multiplex CRISPR

We produced *hat7* single mutants and *hat7 hb13 hb20 hb23* quadruple mutants using FASTRED multiplex CRISPR constructs containing an intronized version of Cas9 (*58*, *62*). Two gRNAs were designed per gene using CHOP-CHOP (*63*). gRNA containing oligos were hybridized and cloned into pDGE sgRNA shuttle vectors using *BpiI* (Data S11). Each of the gRNA containing shuttle vectors were then assembled into pDGE666 (Addgene plasmid # 153231) using BsaI golden gate assembly, sequence verified, and transformed into wild-type Arabidopsis as described above. We selected FASTRED positive T1 seeds and subsequently screened FASTRED negative (putatively Cas9-free) T2 seeds for frameshift mutations using Sanger sequencing coupled with ICE analysis of CRISPR edits (*64*). The edits were similarly confirmed in the T3 generation and at least two homozygous alleles from independent lines were used for experiments.

### BRI1 tissue-specific CRISPR

Two gRNA-BRI1 were simultaneously expressed in a tissue-specific manner (*5*). Primers used for cloning of gRNA BRI1-2 (*65*) and gRNA BRI1-3 can be found in Data S11. The entry module pGG-B-AtU6-26-BRI1-2-C and pGG-A-AtU6-26-BRI1-3-B were generated by annealing oligos for each gRNA and ligating into *BbsI*-digested (New England Biolabs) Golden Gate entry vectors described in (*66*). Next, gRNA modules were combined with pGG-C-linker-G plasmid and cloned into pEN-R2-A-G-L3 by restriction-ligation using BsaI enzyme (New England Biolabs) to obtain pEN-R2-gRNA_BRI1-3-gRNA_BRI1-2-L3. This plasmid was combined with pDONR-L1-Cas9p-tagRFP-L2 (*67*), pDONRL4-L1r carrying either WER or CO2 promoters (*68*) and a destination vector pK8m34GW-FAST (*69*) in a MultiSite Gateway LR reaction (Thermo Fisher Scientific) to obtain expression clones. Expression clones were introduced into Agrobacterium C58 strain and used to transform *pBRI1-BRI1-mCitrine/bri1* plants (*55*) by floral dip. T2 generation seeds were selected based on the presence of GFP signal in the seed coat and 7-day-old seedlings were used for phenotypic analysis. For each root used for quantitative analysis, BRI1-mCitrine signal was acquired in order to confirm efficiency of the tissue-specific knockout system. Statistical analyses were conducted in GraphPad Prism v.9 software.

### Confocal microscopy

Confocal imaging for the majority of experiments was performed using a Zeiss 880 equipped with a 40X objective. Excitation and detection were set as follows: Venus and mCitrine, excitation at 488 nm and detection at 499-571 nm; GFP, excitation at 488 nm and detection at 493–558 nm; PI staining, excitation at 561 nm and detection at 605–695 nm. Confocal images were processed using the Fiji package of ImageJ (*70*). Tile scans were stitched and representative median longitudinal sections for each image are shown. Identical settings were used for images that were directly compared.

For BRI1 TSKO confocal, roots were imaged between a block of agar and cover glass in imaging chambers. Image acquisition was performed with a FluoView1000 inverted confocal microscope (Olympus) equipped with a dry 20X objective (NA 0.75) using 514 nm laser excitation and a spectral detection bandwidth of 500–530 nm for mCitrine and 535 nm laser excitation together with a spectral detection bandwidth of 570– 670 nm for PI.

For the confocal time-lapse video, 7-day-old wild type and *gtl1 df1* seedlings expressing *pC/VIF2-H2B-VENUS* were placed on ½ MS agar blocks containing PI, placed side-by-side in chambered coverglass (Nunc Lab-Tek, ThermoFisher) and imaged under a vertical ZEISS LSM900 microscope equipped with a Plan-Apochromat M27 20x/0.8 n.a. objective. The root tip was imaged every 12 min and automatically tracked with the TipTracker software (*71*). The excitation/emission wavelengths were 514 nm/530-600 nm for Venus and 535 nm/580-650nm for PI.

### BES1 and GTL1 Co-Immunoprecipitation (Co-IP)

Co-IP experiments were conducted as previously described (*72*). p35S-FLAG-GTL1 and a p35S-FLAG-GUS negative control were cloned into pGWB412 (*73*) using gateway LR reactions. The following construct combinations were co-transformed into Arabidopsis mesophyll protoplasts-p35S-BES1-GFP + p35S-FLAG-GUS; p35S-FLAG-GTL1 + p35S-GFP-GUS; p35S-BES1-GFP + p35S-FLAG-GTL1. After overnight incubation, transformed protoplasts were harvested and homogenized in Co-IP buffer (50 mM Tris–HCl, pH 7.5, 150 mM NaCl, 10% (v/v) glycerol, 0.1% (v/v) Nonidet P-40, 1 mM phenylmethysulfonyl fluoride, 20 mM MG132, and proteinase inhibitor cocktail) for 1 h at 4 °C with rotation. 5 μg FLAG M2 antibody (F1804, Sigma) was pre-bound to 40 μL protein G Dynabeads (10003D, Thermo Fisher Scientific) for 30 min in phosphate-buffered saline (PBS) buffer with 0.02% Tween 20 at room temperature. The beads were washed once with the same PBS buffer and resuspended in Co-IP buffer. After protein extraction, 10 μL of anti-FLAG pre-bound Dynabeads was added to each sample for another 1.5 h incubation at 4 °C with rotation. Dynabeads were precipitated using a DynaMagnetic rack (12321D, Thermo Fisher Scientific) and washed twice with Co-IP buffer with Nonidet P-40 and three times with Co-IP buffer without Nonidet P-40. The IP products were eluted in 2XSDS sample buffer and used for immunoblotting with rabbit anti-GFP (A11122, Invitrogen) and rabbit anti-FLAG antibody (F7425, Sigma–Aldrich) at 1:1,000 dilution.

### scRNA-seq profiling of Arabidopsis root protoplasts using the 10X Genomics chromium system

scRNA-seq experiments were performed as previously described (*21*) with minor modifications. Plants were grown for 7 days as described above with the addition of 100 μm nylon mesh (Nitex 03-100/44) on the plates to facilitate root collection. For each sample, ~0.5cm root tips were harvested from 1000-3000 roots and placed into a 35mm petri dish containing a 70 μm cell strainer and 4.5mL enzyme solution (1.5% [w/v] cellulase [ONOZUKA R-10, GoldBio], 0.1% Pectolyase [Sigma P3026], 0.4 M mannitol, 20 mM MES (pH 5.7), 20 mM KCl, 10 mM CaCl2, 0.1% bovine serum albumin, and 0.000194% (v/v) beta-mercaptoethanol). The digestion was incubated on an 85 rpm shaker at 25°C for one hour with additional stirring every 15-20 minutes. The resulting cell solution was filtered twice through 40 μm cell strainers and centrifuged for 5 minutes at 500g in a swinging bucket rotor. The pellet was washed with 2mL washing solution (0.4 M mannitol, 20 mM MES (pH 5.7), 20 mM KCl, 10 mM CaCl2, 0.1% bovine serum albumin, and 0.000194% (v/v) beta-mercaptoethanol), centrifuged again at 500g for 3 minutes, and the pellet resuspended in washing solution at a concentration of ~2000 cells/uL. We loaded 16,000 cells, with the aim to capture 10,000 cells per sample with the 10X Genomics Chromium 3‘ Gene expression v3 or v3.1 kits. Cell barcoding and library construction were performed following the manufacturer’s instructions. cDNA and final library quality were verified using a Bioanalyzer High Sensitivity DNA Chip (Agilent) and sequenced on an Illumina NextSeq 500 or NovaSeq 6000 instrument to produce 100bp paired-end reads.

For BL scRNA-seq, we first grew plants on 1 μM BRZ to deplete endogenous BRs, then transferred plants to either a fresh BRZ plate or 100nM BL. We monitored the efficacy of these treatments using a constitutively expressed 35S-BES1-GFP line. In agreement with previous reports (*12*, *74*, *75*), BES1-GFP was predominantly present in the cytoplasm under low BR conditions resulting from BRZ treatment but accumulated in the nucleus following BL treatment (Fig. S1). We performed two separate BL scRNA-seq treatment experiments. The first consisted of a BRZ and 2 hour BL treatment. The second experiment included two additional replicates of BRZ and BL 2 hours along with the other time points in our time course (BL 0.5, 1, 4, and 8 hour treatments). Each of the BL treatments was staggered so that all samples were collected simultaneously. A total of 70,223 cells were recovered from the BL treatment scRNA-seq experiments.

Wild type Col-0, *bri1-T*, and *pGL2-BRI1-GFP/bri1-T* were similarly profiled in a side-by-side scRNA-seq experiment under control conditions with two replicates per genotype, resulting in 34,861 cells. Lastly, scRNA-seq was performed on Wild type Col-0, *gtl1, df1*, and *gtl1 df1* in duplicate under control conditions, resulting in 74,810 scRNA-seq expression profiles.

### scRNA-seq data pre-processing

Raw sequencing reads were demultiplexed from Illumina BCL files to produce FASTQ files for each sample using CellRanger mkfastq (v3.1.0, 10X Genomics). Reads were then aligned against the Arabidopsis TAIR10 reference genome to generate a gene-by-cell matrix using the scKB script (https://github.com/ohlerlab/scKB), which incorporates kallisto (*76*) and bustools (*77*, *78*). Quality filtering of cells was performed using the R package COPILOT (Cell preprOcessing PIpeline kaLlistO busTools) (*21*, *79*). COPILOT uses a non-arbitrary scheme to remove empty droplets and dying or low-quality cells. We used one iteration of COPILOT filtering, which adequately separated high-quality cells from the background in our samples based on an examination of barcode rank plots. To address issues with doublets and outliers, the resulting high-quality cells were further filtered to remove the top 1% of cells in terms of UMI counts, and putative doublets were removed with DoubletFinder(*80*) using the estimated doublet rate from the 10X Genomics Chromium Single Cell 3’ Reagent Kit user guide.

### Normalization, annotation, and integration of scRNA-seq datasets

Downstream analysis were carried out using Seurat version 3.1.5. Samples were first individually processed and examined. Data were normalized using SCTransform (*81*) and all detected genes were subsequently retained for analysis, except those from mitochondria, chloroplasts or those affected by protoplasting (absolute log2 fold-change >= 2) (*21*, *23*). Principal component analysis (PCA) was performed by calculating 50 principal components using the RunPCA function (with approx=FALSE). UMAP non-linear dimensionality reduction was next calculated via the RunUMAP function using all 50 principal components with parameters n_neighbors = 30, min_dist = 0.3, umap.method = ‘ ‘umap-learn’ ’, metric = ‘ ‘correlation’ ’. These processing steps have been previously described (*21*) and are documented in jupyter notebooks as part of the COPILOT workflow.

To follow the developmental progression from the meristem to the elongation zone more closely, we updated the root atlas (*21*) developmental annotation to subdivide the meristem into the proliferation domain and transition domain as previously defined (*22*). The previous meristem annotation of the root atlas was based on correlation annotation by comparing each cell from scRNA-seq to bulk data from morphologically defined sections (*82*). On the other hand, HIGH PLOIDY2 was used to mark the meristem in a second bulk expression profile (*83*), which corresponds to the proliferation domain defined by Ivanov and Dubrovsky. Therefore, we leveraged correlation-based annotations derived from Li et al., 2016 to re-label the meristem of the atlas. If cells were defined as “meristem” by both Li et al., 2016 and Brady et al., 2007, then they were re-labeled as the proliferation domain. Those that were called elongation in the Li et al., 2016 annotation, but meristem in the Brady et al., 2007 annotation were re-labeled as the transition domain. Finally, cells labeled as elongation in both Brady and Li datasets but annotated as meristem in the root atlas were re-labeled as elongation zone.

Consistent with our annotation, we found that cell cycle-related genes were enriched in the proliferation domain of the atlas (fig. S2B), whereas *SMR1* (AT3G10525), a marker of endoreduplication, increased in the transition domain (*84*). The developmental annotation of cortex markers *CO2* (AT1G62500) and *CORTEX* (AT1G09750) also matched their expression patterns in the root (*85*, *86*).

We used the receiver operating characteristic (ROC) test implemented in Seurat FindMarkers to identify genes enriched in each developmental zone. A largely distinct set of markers was enriched in the transition domain (fig. S2C and Data S2). These included genes involved in vesicle-mediated transport (fig. S21D), in line with the observation that vesicle recycling activity is highest in this region (*87*).

We transferred the cell type and developmental stage labels from the wild-type atlas (*21*) to each sample using the Seurat label transfer workflow (*88*, *89*). To align corresponding cell types and developmental stages, we integrated samples from each experiment using the Seurat reference-based integration pipeline (*88*, *89*). A sample from the atlas with the highest genes detected (sc_12) was used as a reference (*21*) and two previously described samples (dc_1 and dc_2) (*23*) were also included in each integration. PCA and UMAP were subsequently calculated for each integration object using the batch-corrected “integrated” assay as described above. Although sc_12, dc_1 and dc_2 were not used in any subsequent analysis, their inclusion at the integration step helped to generate a comparable UMAP projection among different integration objects that facilitates interpretability.

### Plotting gene expression values on the UMAP projection

We subsequently examined changes in cell state caused by the BL treatments or in the mutants profiled by plotting the log-normalized, ‘corrected’ counts produced by the SCTransform function (Hafemeister and Satija, 2019) rather than the batch-corrected “integrated” values when visualizing changes in expression.

### Pseudotime estimation and heatmaps of gene expression trends

Cortex cells were extracted from the integrated Seurat objects (BR time course, *bri1-T* vs wild type and *gtl1 df1* vs wild type). Pseudotime was then inferred on the SCT assay of the extracted cortex cells using CytoTRACE v0.1.0 (Gulati et al., 2020). Once the pseudotime was calculated, the cortex cells were converted into SingleCellExperiment objects (*90*) before fitting a NB-GAM model (generalized additive model with a negative binomial distribution) using fitGAM function of tradeSeq R package v1.8.0 (*91*). The model-predicted expression trends were plotted with ComplexHeatmap in R (Gu et al., 2016) (v2.10.0).

### Pseudobulk differential expression analysis

Recent benchmarks point towards pseudobulk methods, which aggregate cell level counts for subpopulations of interest on a per-sample basis, as top performers for cross-condition comparisons in scRNA-seq (*92*, *93*). Therefore, we employed a pseudobulk approach implemented in muscat (Multi-sample multi-group scRNA-seq analysis tools) (*92*) to examine changes in each combination of cell type and developmental stage. Pseudobulk expression profiles were aggregated for each of these subpopulations by summing the raw counts using the aggregateData function. We then performed differential expression testing using the edgeR method (*94*) incorporated in the pbDS function. A term for the experimental batch and/or replicate was included in the contrast to adjust for potential batch effects. A gene was considered differentially expressed in a given subpopulation if the false discovery-rate adjusted p-value was <=0.05, absolute fold change was >=1.5 and detection frequency was >=10% in one of the conditions. Gene ontology enrichment analysis was conducted on the differentially expressed genes using the R package “gprofiler2” (*95*). Comparisons between DEG lists were performed using the GeneOverlap package (version 1.12.0; http://shenlab-sinai.github.io/shenlab-sinai/). p-values for intersections between gene lists were computed using Fisher’s exact test. Visualizations were generated using Seurat (*88*), ComplexHeatmap (*96*), and ggplot2 (*97*).

### WOT differential expression along cortex cell wall + trajectories

WOT constructs trajectories of cells from a reference time point by minimizing the difference over all genes (*31*). The algorithm requires as input the expression profiles of cells as well as an estimation of their proliferation rate. We estimated proliferation rates using imaging data (*98*), as previously described (*99*). As only the bottom 0.5 cm of each root was observed at each time, we expect some cells to exit the observed section due to proliferation. We estimated the number of cells that should exit the observed section based on the growth rate with the assumption that the section of root stays in equilibrium and assigned the calculated number of cells with the highest pseudotime a growth rate of zero so that they have no descendents in the observed root section at the next time point. We constructed trajectories using full gene expression profiles and evaluated the quality of the trajectories by checking the proportion of cells whose highest fate probability matched the annotation. We found that for 90% of cells the largest fate assigned by WOT matched the annotation, rising to 97% in the maturation zone where we have the greatest confidence in the annotation.

The cell wall signature was calculated for each cell by taking the sum of Z-scores for each of the 107 BR-induced cell wall-related genes in the signature (GO:0071554), truncated to [-5,5]. We defined the cortex cell wall+ subset as cortex cells with a cell wall score greater or equal to 1. This threshold was chosen as it selected less than 5% of cells from other lineages while still retaining >20% of cortex cells at the 2 hour time point. Any cortex cell that did not belong to the cortex cell wall+ group was labelled as “cortex cell wall−”. We performed differential expression on the cortex cell wall+ and cell wall-subsets at 2 hours, using WOT lineages to also perform differential expression on their putative ancestors and descendants. Statistical significance was evaluated using Welch’s t-Test with adjusted p-values for multiple tests, requiring t_FDR_ < 0.01. Results were ranked by the adjusted expression ratio

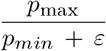

with *ε* = 0.1, where larger *ε* puts more emphasis on genes with non-zero expression in both groups. To generalize our WOT analysis we also constructed trajectories for each combination of cell type and developmental stage and performed differential expression analysis between each time point using the same process (Data S5).

### Gene regulatory networks

In order to construct GRNs, we used CellOracle (v0.7.0) for single-cell GRN inference (*48*). In the first stage of the CellOracle pipeline, a base GRN is defined, representing a global set of biologically plausible Transcription factor-Target interactions. We used publicly available scATAC-seq data from Arabidopsis roots (*100*) GSE155304:GSM4698760; (*100*) to determine regions of open chromatin. Cell Ranger ATAC (v1.2.0) was used to process raw scATAC seq data to call a peak-by-cell matrix. Cicero (v1.11.1) (*101*) was implemented to infer a co-accessibility map of chromatin regions. Transcription start sites were then annotated based on the Arabidopsis TAIR10 genome assembly. Finally, peaks with weak co-accessibility scores were filtered following instructions of CellOracle manual (https://morris-lab.github.io/CellOracle.documentation/tutorials/base_grn.html). To expand the number of TFs present in the base GRN we also included TF-Target interactions from DNA affinity purification sequencing (DAP-seq) (*102*) and a previously constructed integrative gene regulatory network (iGRN) (*103*). Our resulting base GRN contained 11.7 million interactions between 1,601 transcription factors and 31,019 target genes.

In the second step of the CellOracle pipeline, a regularized machine learning approach is used to define active edges and their regulatory strength in clusters or subpopulations of scRNA-seq data. In this process, the expression of target genes is predicted based on regulatory transcription factor levels from the base GRN. Inactive edges with low predictive ability are pruned from the base GRN, revealing context-specific GRN configurations (*48*).

To test CellOracle on *Arabidopsis* root data, we first inferred GRN configurations for each of the 36 cell type and developmental stage combinations in our WT atlas (*21*) using the SCT normalized counts. We limited the base GRN to genes dynamically expressed along pseudotime for each cell type plus associated transcription factors (*104*). Each cell type GRN was then constructed with default parameters following the CellOracle manual. To filter network edges with the “filter_links” function, we retained the top 20,000 edges (p-value <=0.01) for each subnetwork. This recovered known developmental regulators (Data S6 and Data S7) including MYB36 in the endodermis (*105*) and BRN1/BRN2 in the root cap (*106*), confirming that CellOracle analysis of Arabidopsis root scRNA-seq data can infer GRNs configurations for particular cell identities and states.

We implemented similar procedures to infer context-specific GRN configurations for each cell type, developmental stage and time point of the BR time course samples (sc_43-50). We used transcription factors plus DEGs from BL 2 hour vs.

BRZ pseudobulk analysis of each cell type/developmental zone combination. The resulting set of 201 GRN configurations spanned 767,970 edges between 1,164 transcription factors and 7,135 targets (Data S8). Network centrality measures were calculated using the built-in functions of the CellOracle pipeline (Data S9). The data needed to reproduce our results and jupyter notebooks demonstrating the processes are available on ARVEX (https://shiny.mdc-berlin.de/ARVEX/).

## Data and code availability

Single-cell RNA-seq data have been deposited at GEO: GSE212230. All original code is available at https://github.com/tmnolan/Brassinosteroid-gene-regulatory-networks-at-cellular-resolution.

## ACKNOWLEDGEMENTS

This work was funded by the US National Science Foundation (Postdoctoral Research Fellowships in Biology Program grant no. IOS-2010686 and MCB 1818160) to T.M.N. and Y.Y., respectively; US National Institutes of Health (NRSA post-doctoral fellowship 1F32GM136030-01 and MIRA 1R35GM131725) to R.S. and P.N.B., respectively; Research Foundation-Flanders (project no. G002121N to E.R. and a postdoctoral fellowships no. 12R7822N and no. 12R7819N to N.V.); Deutsche Forschungsgemeinschaft (International Research Training Group 2403) to C.-W.H. and U.O.; USDA-NIFA 2021-67034-35139 to I.W.T.; a Burroughs Wellcome Fund Career Award, NFRF Exploration Grant, NSERC Discovery Grant, and CIHR Project Grant to L.G., M.H., A.A., and G.S.; and by the Howard Hughes Medical Institute to P.N.B. as an Investigator. The authors thank Heather Belcher and Megan Perkins Jacobs for technical assistance; Nicolas Devos and Duke GCB for sequencing services; Nick Vangheluwe, Tom Beeckman and Thomas B. Jacobs (VIB-UGent) for CRISPR-TSKO advice; and Keiko Sugimoto for *gtl1* and *df1* seeds. This manuscript was formatted using the Henrique’s lab bioRxiv template on Overleaf.

## AUTHOR CONTRIBUTIONS

T.M.N., N.V., C.-W.H., E.R., and P.N.B., conceptualized the experiments. T.M.N., R.S., and I.W.T. generated the scRNA-seq data. C.-W.H., T.M.N., L.G., M.H., A.A., and G.S. analyzed scRNA-seq data. N.V. and I.V. generated and characterized BRI1-TSKO lines. T.M.N. and J.Z., generated mutants using multiplex CRISPR. T.M.N., J.Z., and P.S. constructed reporter lines. T.M.N. and A.B. performed confocal imaging. P.W. performed Co-IP experiments.

C.-W.H. and T.M.N. developed the ARVEX scRNA-seq browser. T.M.N. wrote the manuscript with input from all authors. Y.Y., G.S., U.O., E.R., and P.N.B. supervised the experiments and analyses.

## COMPETING FINANCIAL INTERESTS

P.N.B. is the co-founder and Chair of the Scientific Advisory Board of Hi Fidelity Technologies, a company that works on crop root growth.

---

**Movie S1**.

Time-lapse confocal microscopy showing pC/VIF2-H2B-Venus in wild-type and *gtl1 df1*.

**Data S1**.

Summary of the scRNA-seq samples reported in this study.

**Data S2**.

Marker genes for updated developmental annotation of Shahan et al WT root atlas.

**Data S3**.

DEGs from pseudobulk analysis of scRNA-seq datasets.

**Data S4**.

DEGs from WOT analysis of cortex cell wall + scRNA-seq datasets.

**Data S5**.

DEGs from WOT analysis of each cell type and developmental stage from BR time series scRNA-seq.

**Data S6**.

CellOracle GRN inferred from wild-type atlas.

**Data S7**.

CellOracle GRN centrality metrics from wild-type atlas.

**Data S8**.

CellOracle GRN inferred from BR time series.

**Data S9**.

CellOracle GRN centrality metrics from BR time series.

**Data S10**.

Predicted targets of HAT7 and GTL1 family TFs from elon-gating cortex CellOracle GRNs in BR time series.

**Data S11**.

Oligos used in this study.

Note that Supplemental data that exceeds the file size limit can be found at: *https://shiny.mdc-berlin.de/ARVEX*/

**Fig. S1.**
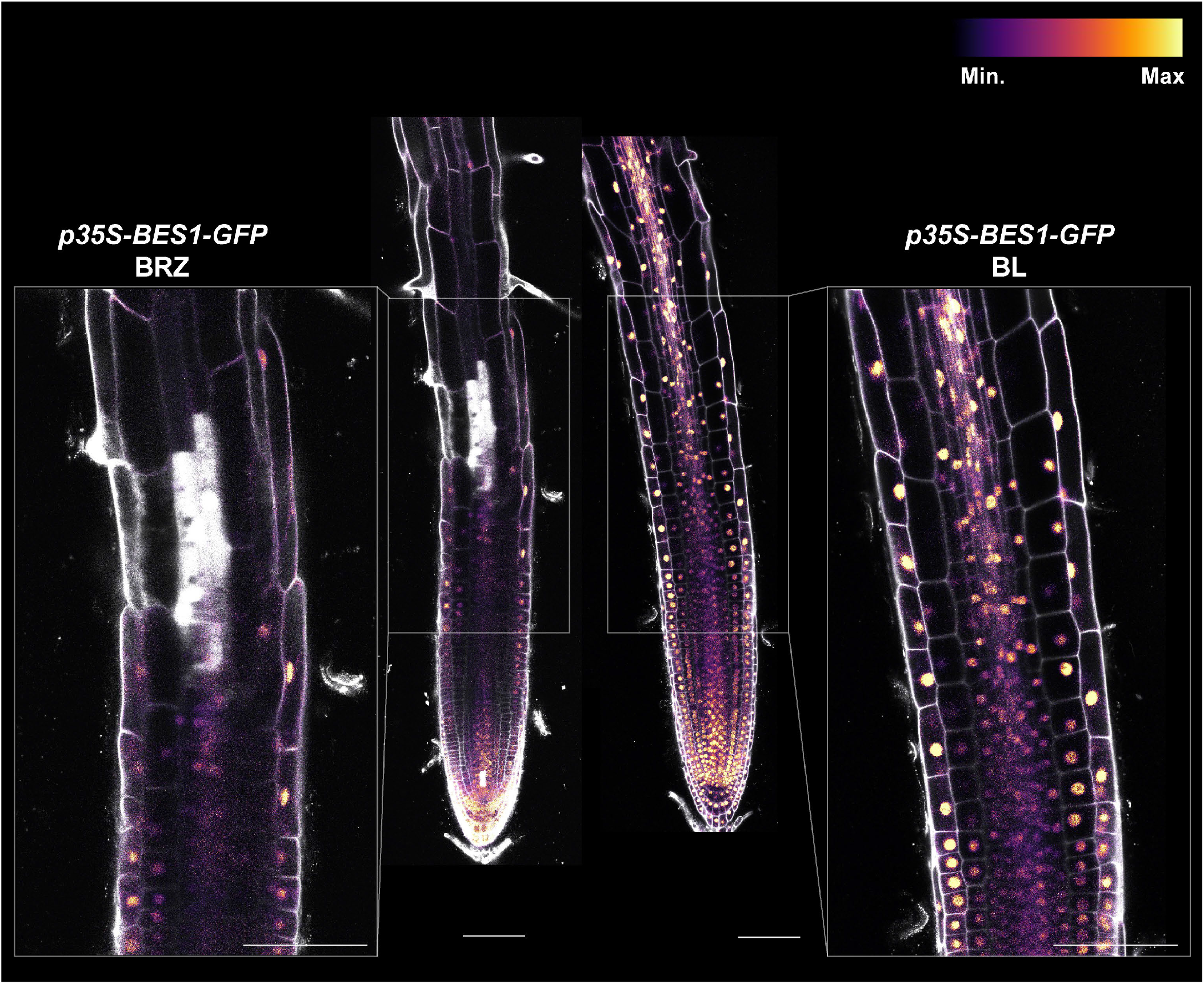
A sensitized system to study spatiotemporal brassinosteroid response. Plants were grown for 7 days on 1 μM Brassinazole (BRZ) to deplete endogenous BRs, then transferred plants to either a fresh BRZ plate or 100 nM Brassinolide (BL) for 2 hours. p35S-BES1-GFP was used to monitor the efficacy of the sensitized system for BL scRNA-seq. BES1-GFP was predominantly present in the cytoplasm under low BR conditions resulting from BRZ treatment but accumulated in the nucleus following BL treatment.

**Fig. S2.**
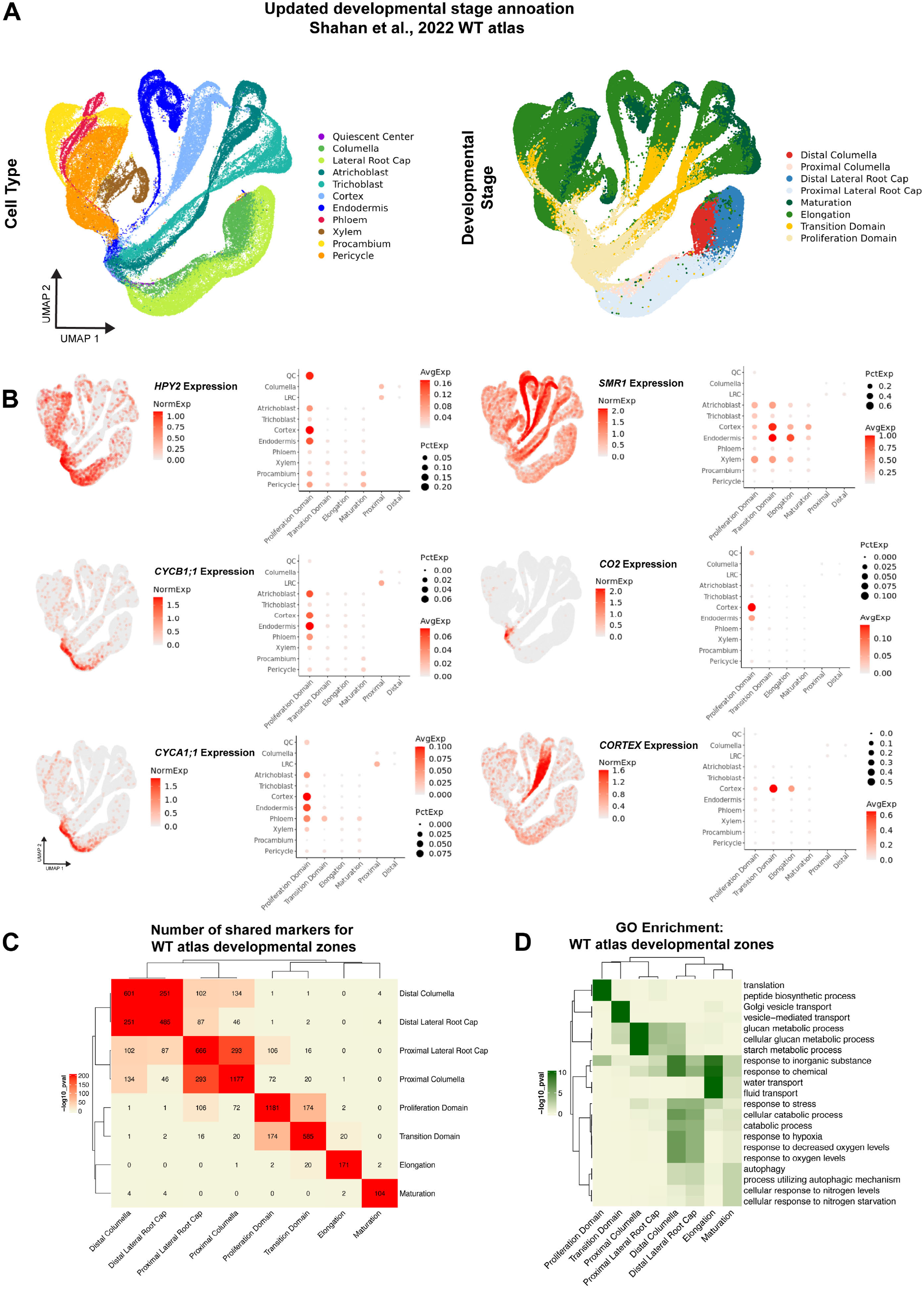
Updated developmental annotation distinguishes between the proliferation domain and transition domain of the meristem. (A) Wild-type atlas of the Arabidopsis root from (*21*) showing updated developmental stage annotation in which the meristem is separated into the proliferation domain and transition domain. (B) Expression of markers in wild-type atlas supporting the developmental stage annotation. The color scale on the UMAP projection represents log normalized, corrected UMI counts. In dotplots, the size of the dot represents the percentage of cells in which the gene is expressed. (C) Comparison of the number of shared markers for each of the developmental zones in the wild-type atlas. Color represents log10 p-values from the indicated overlaps calculated from Fisher’s exact test by GeneOverlap. The number of genes in each intersection is indicated inside each box. (D) Enriched GO terms for markers of each developmental zone in the wild-type atlas.

**Fig. S3.**
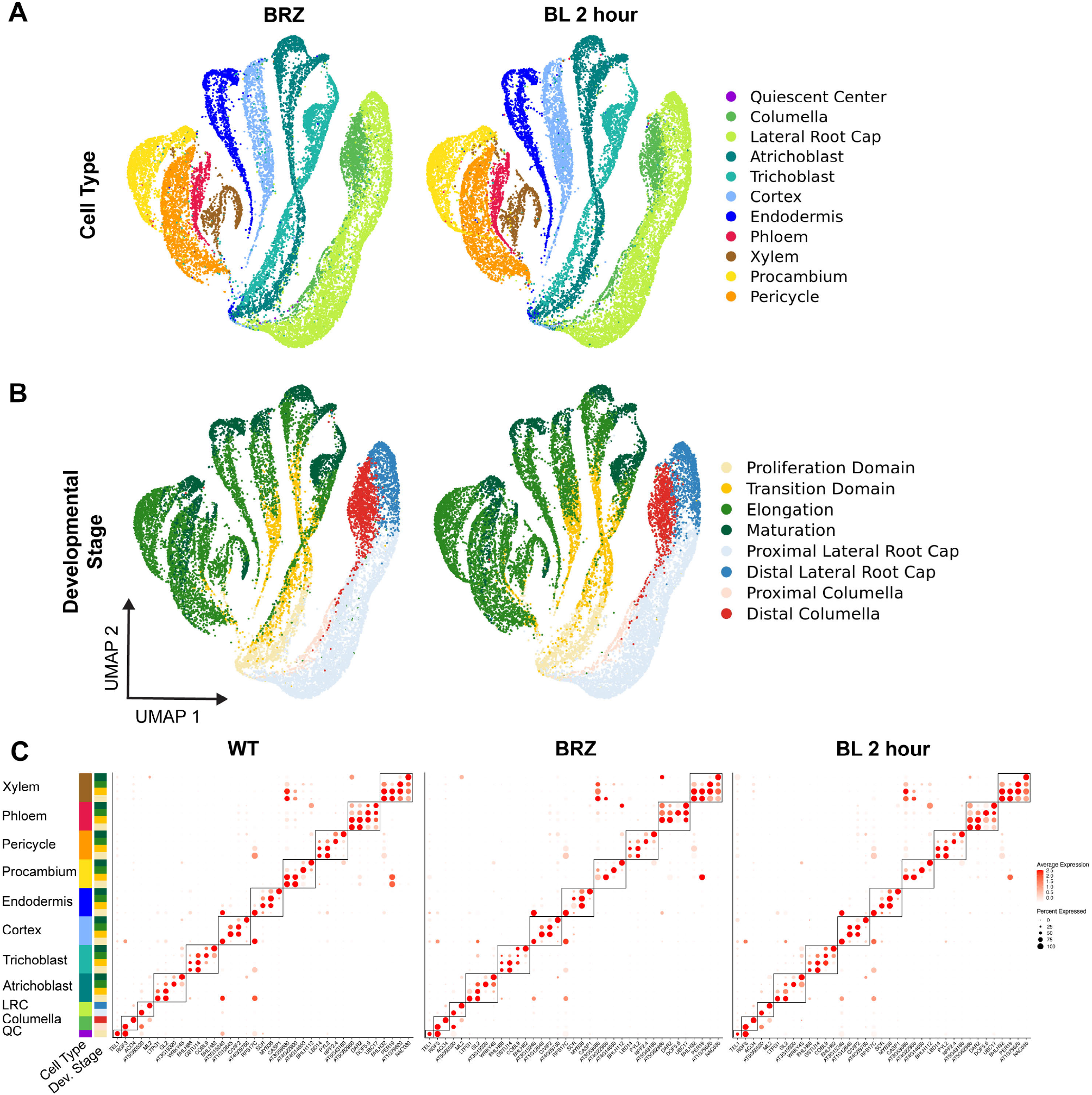
Cell types and developmental stages are identified after label transfer in BL scRNA-seq. Two-dimensional uniform manifold approximation and projection (UMAP) embedding of 21,473 BRZ and 22,275 two-hour BL treated cells across 3 biological replicates of scRNA-seq. Colors indicate **(A)** cell type or **(B)** developmental stage annotation. **(C)** Dotplots from the WT root altas(*21*), BRZ, and BL 2 hours scRNA-seq showing that cell types and developmental stages are identified through label transfer. One marker gene for each cell type and developmental stage combination is shown. Circle size represents the percentage of cells in which a gene is expressed and color represents the average expression level of each gene. Black boxes denote markers from each cell type. Colors of side annotations indicate cell type and developmental stage.

**Fig. S4.**
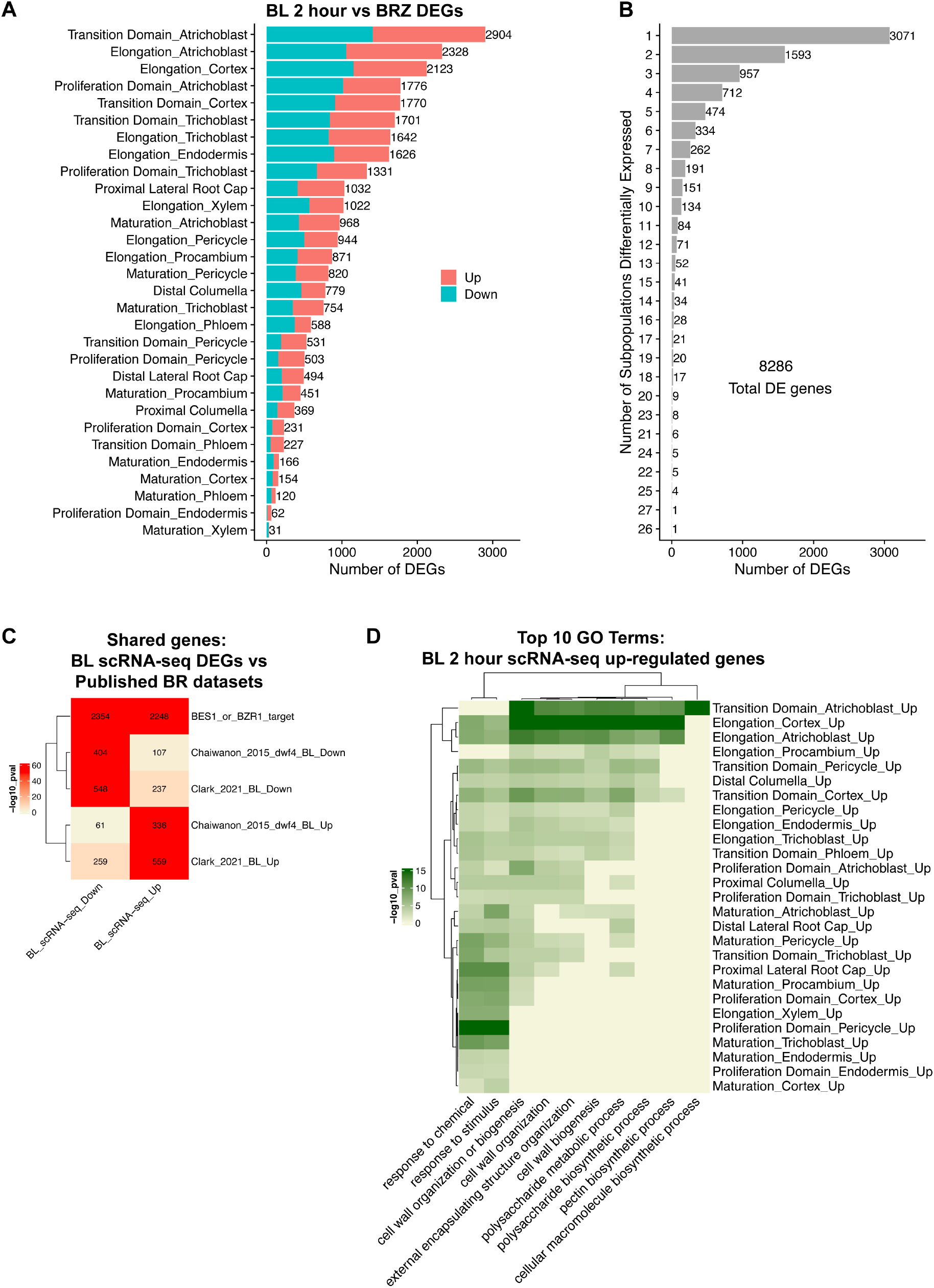
scRNA-seq identifies the elongating cortex as a site of BR-response. **(A)** Number of DEGs for each cell type/developmental stage combination in BL 2 hour scRNA-seq. Color indicates the number of up-regulated vs down-regulated genes. **(B)** The number of cell type/developmental stage combinations (subpopulations) in which each BL DEG is differentially expressed. 3,071/8,286 DEGs were significantly altered in only a single sub-population. **(C)** Comparison of BL 2 hour DEGs from scRNA-seq to BES1 and BZR1 ChIP targets and previous bulk BR RNA-seq datasets. Color represents log10 p-values from the indicated overlaps calculated from Fisher’s exact test by GeneOverlap. The number of genes in each intersection is indicated inside each box. **(D)** Top 10 GO terms in terms of p-value among BL up-regulated DEGs from scRNA-seq. Note the strong enrichment for cell wall-related GO terms in BL up-regulated genes in the elongating cortex.

**Fig. S5.**
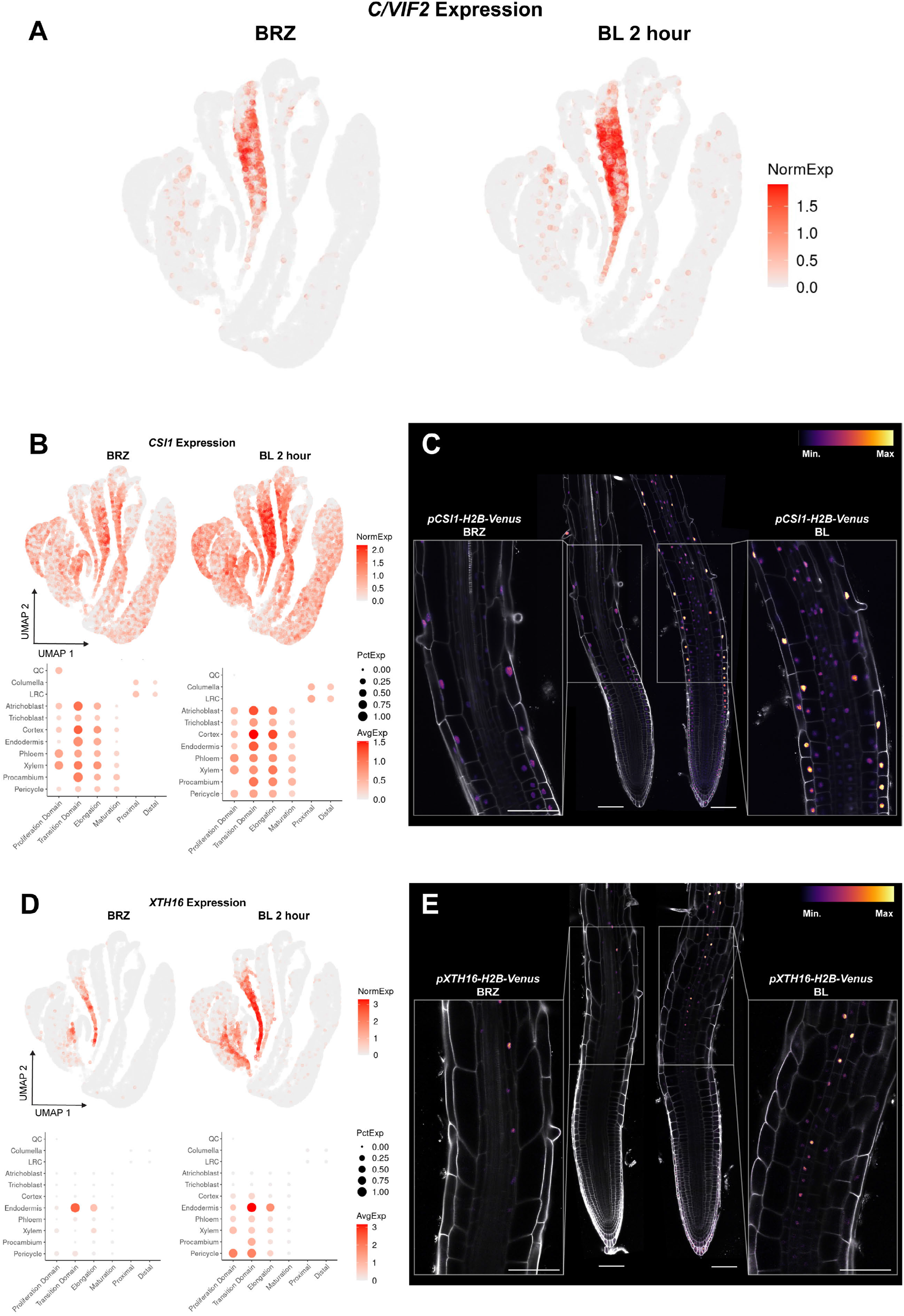
Reporter gene expression is consistent with BL scRNA-seq. **(A)** Expression of *C/VIF2* in BRZ and BL 2 hour scRNA-seq. **(B)** *CSI1* expression in BRZ and BL 2 hour scRNA-seq. The color scale on the UMAP projection represents log normalized, corrected UMI counts. In dotplots, the size of the dot represents the percentage of cells in which the gene is expressed. **(C)** *CSI1-H2B-Venus* reporter grown on 1 μM BRZ for 7 days and transferred to 1 μM BRZ or 100 nM BL for 4 hours. Inset shows *CSI1* signals that are strongest in cortex and epidermis and increase with BL treatment. Propidium iodide-staining is shown in grey, with the color gradient indicating relative *CSI1-H2B-Venus* levels. **(D)** *XTH16* expression in BRZ and BL 2 hour scRNA-seq. **(E)** *XTH16-H2B-Venus* reporter grown on 1 μM BRZ for 7 days and transferred to 1 μM BRZ or 100 nM BL for 4 hours. Inset shows *XTH16* signals in the endodermis that increase with BL treatment. Propidium iodide-staining is shown in grey, with the color gradient indicating relative *XTH16-H2B-Venus* levels. Scale bars, 100μm.

**Fig. S6.**
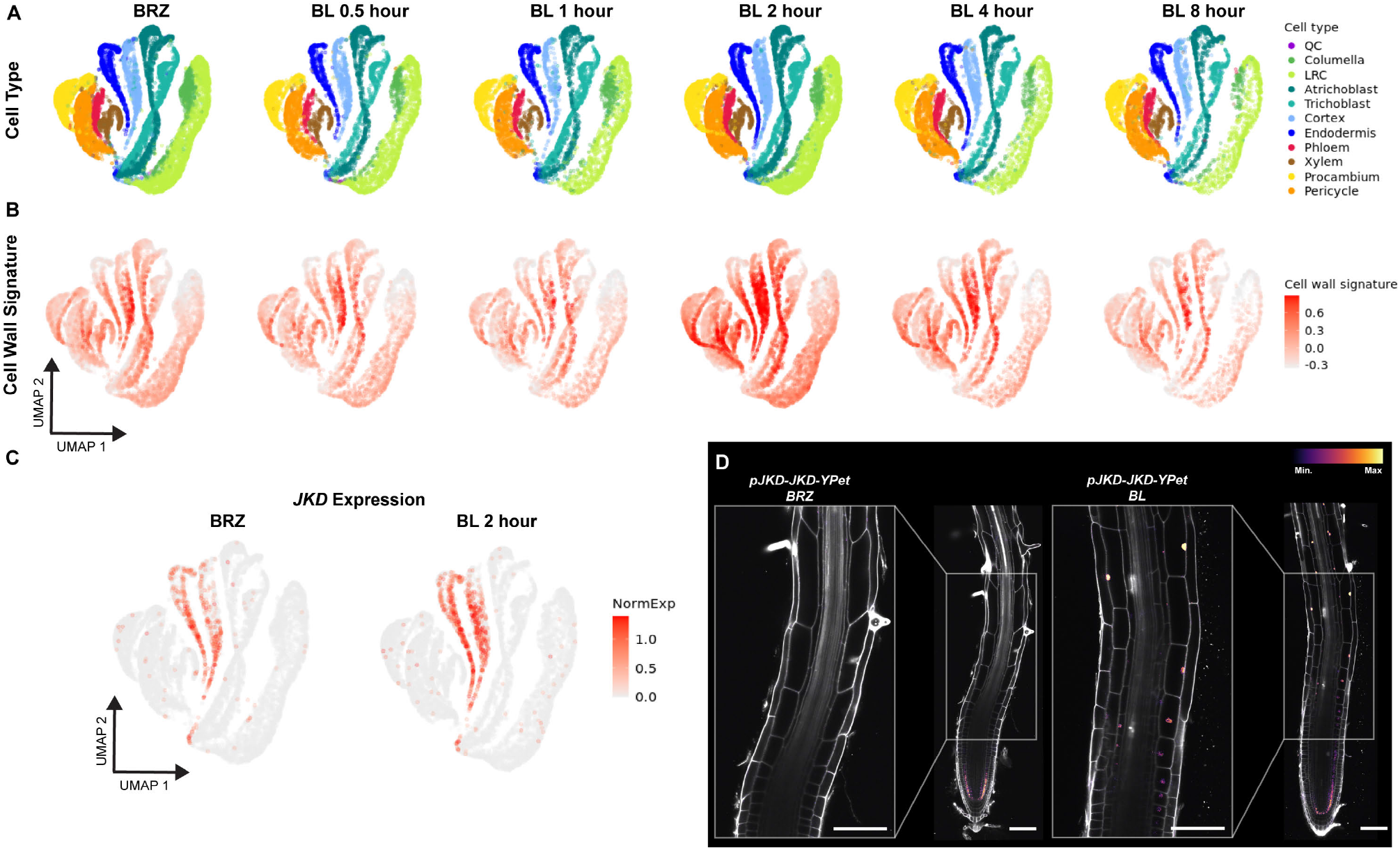
Waddington optimal transport identifies JKD as a BR responsive TF along cortex trajectories. **(A)** UMAP showing cell type annotation across BL scRNA-seq treatment time course. This panel is repeated from the main text figure as a reference for the panel below. **(B)** UMAP projection colored by cell wall signature, calculated as the sum of Z-scores for each of the 107 BR-induced cell wall-related genes in the signature (GO:0071554), truncated to [-5,5]. **(C)** Expression of *JKD* in BRZ and BL 2 hour scRNA-seq. The color scale represents log normalized, corrected UMI counts. **(D)** *pJKD-JKD-YPet* grown on 1 μM BRZ for 7 days and transferred to 1 μM BRZ or 100 nM BL for 4 hours. Inset shows *JKD* signals in the elongating cortex that increase with BL treatment. Propidium iodide-staining is shown in grey, with the color gradient indicating relative JKD-Ypet levels. Scale bars, 100 μm.

**Fig. S7.**
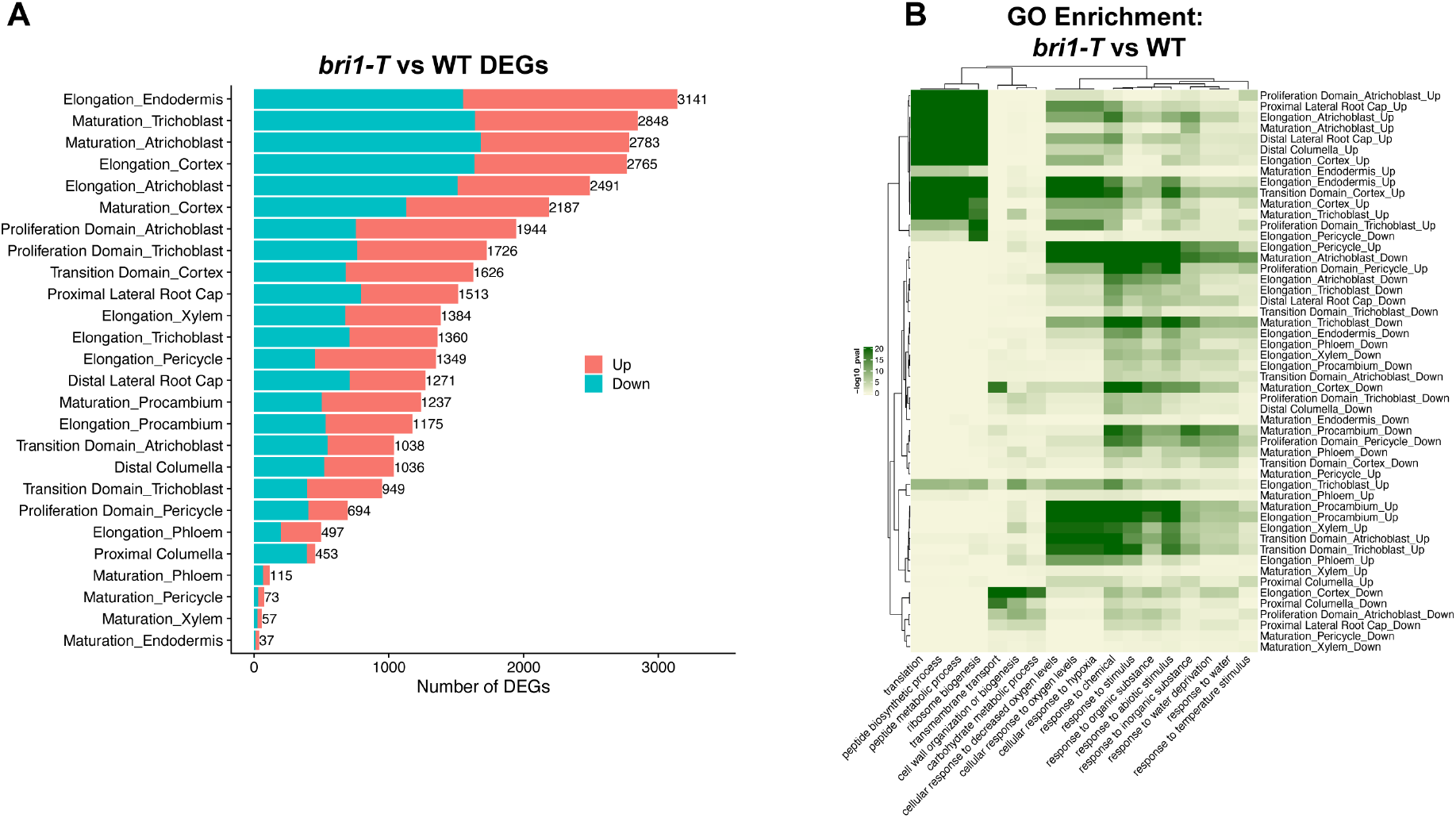
Analysis of the triple receptor mutant *bri1-T* reveals endogenous regulation of cell-wall-related genes by BRs in the elongating cortex. **(A)** Number of DEGs for each cell type/developmental stage combination in *bri1-T* scRNA-seq compared to WT. Color indicates the number of up-regulated vs down-regulated genes. **(B)** GO enrichment of *bri1-T* vs wild-type DEGs. Note the strong enrichment for cell wall-related GO terms in *bri1-T* down-regulated genes in the elongating cortex.

**Fig. S8.**
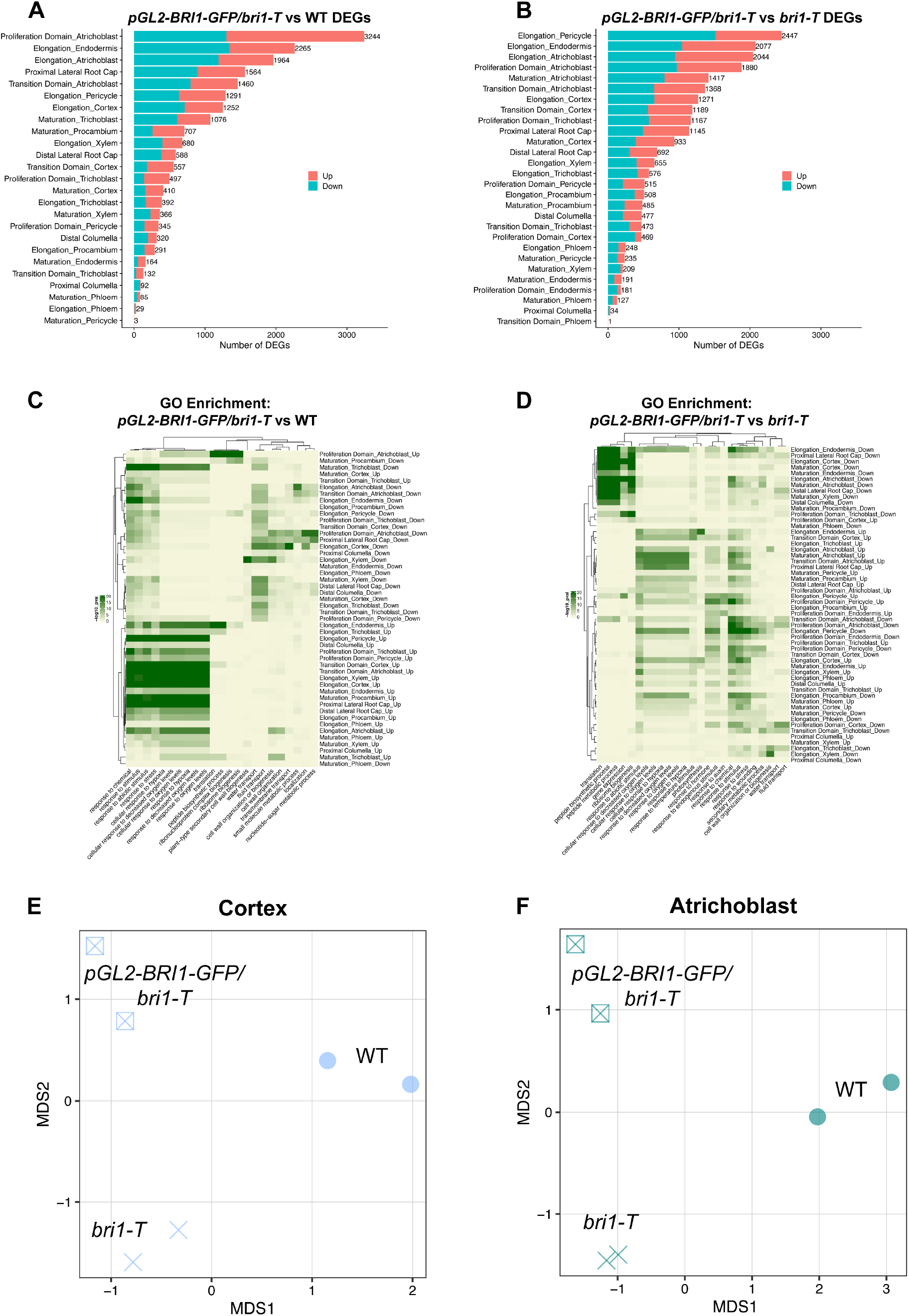
Distinct gene expression patterns of *pGL2-BRI1-GFP/bri1-T*. **(A)** Number of DEGs for each cell type/developmental stage combination in *pGL2-BRI1-GFP/bri1-T* scRNA-seq compared to WT. Color indicates the number of up-regulated vs down-regulated genes. **(B)** GO enrichment of *pGL2-BRI1-GFP/bri1-T* vs wild-type DEGs. **(C)** Number of DEGs for each cell type/developmental stage combination in *pGL2-BRI1-GFP/bri1-T* scRNA-seq compared to *bri1-T*. Color indicates the number of up-regulated vs down-regulated genes. **(D)** GO enrichment of *pGL2-BRI1-GFP/bri1-T* vs *bri1-T* DEGs. **(E)** Multidimensional scaling (MDS) analysis of cortex cells from scRNA-seq. Note that replicates from the same genotype group together, but genotypes are well separated. **(F)** Multidimensional scaling (MDS) analysis of atrichoblast cells from scRNA-seq.

**Fig. S9.**
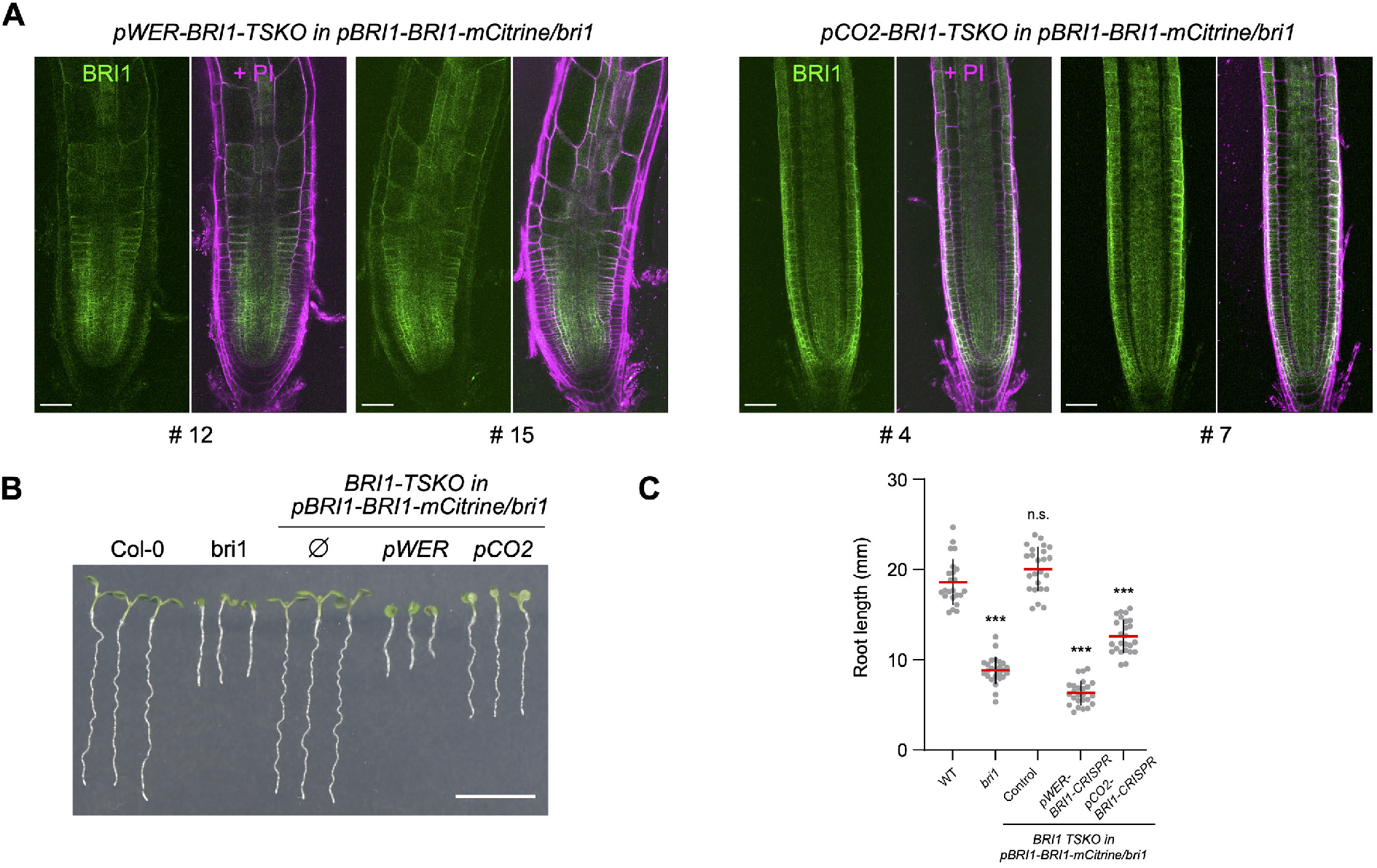
BRI1 CRISPR tissue-specific CRISPR. **(A)** Two individual transgenic lines for *pWER-BRI1-CRISPR* and *pCO2-BRI1-CRISPR* exhibiting similar BRI1-mCitrine expression patterns. Scale bars, 50 μm. **(B)** Seven-day-old BRI1-CRISPR transgenic seedlings with roots shorter than those of the wild-type (Col-0) control and complemented *pBRI1-BRI1-mCitrine/bri1*. Scale bar represents 1 cm. **(C)** Quantification of the root length of transgenic lines shown in (B). All individual data points are plotted. Red horizontal bars represent the means and error bars represent s.d. Significant differences between transgenic lines and the WT control were determined by one-way ANOVA and Dunnett’s multiple comparisons tests. ***P<0.001, **P<0.01, *P<0.05. n.s. not significant.

**Fig. S10.**
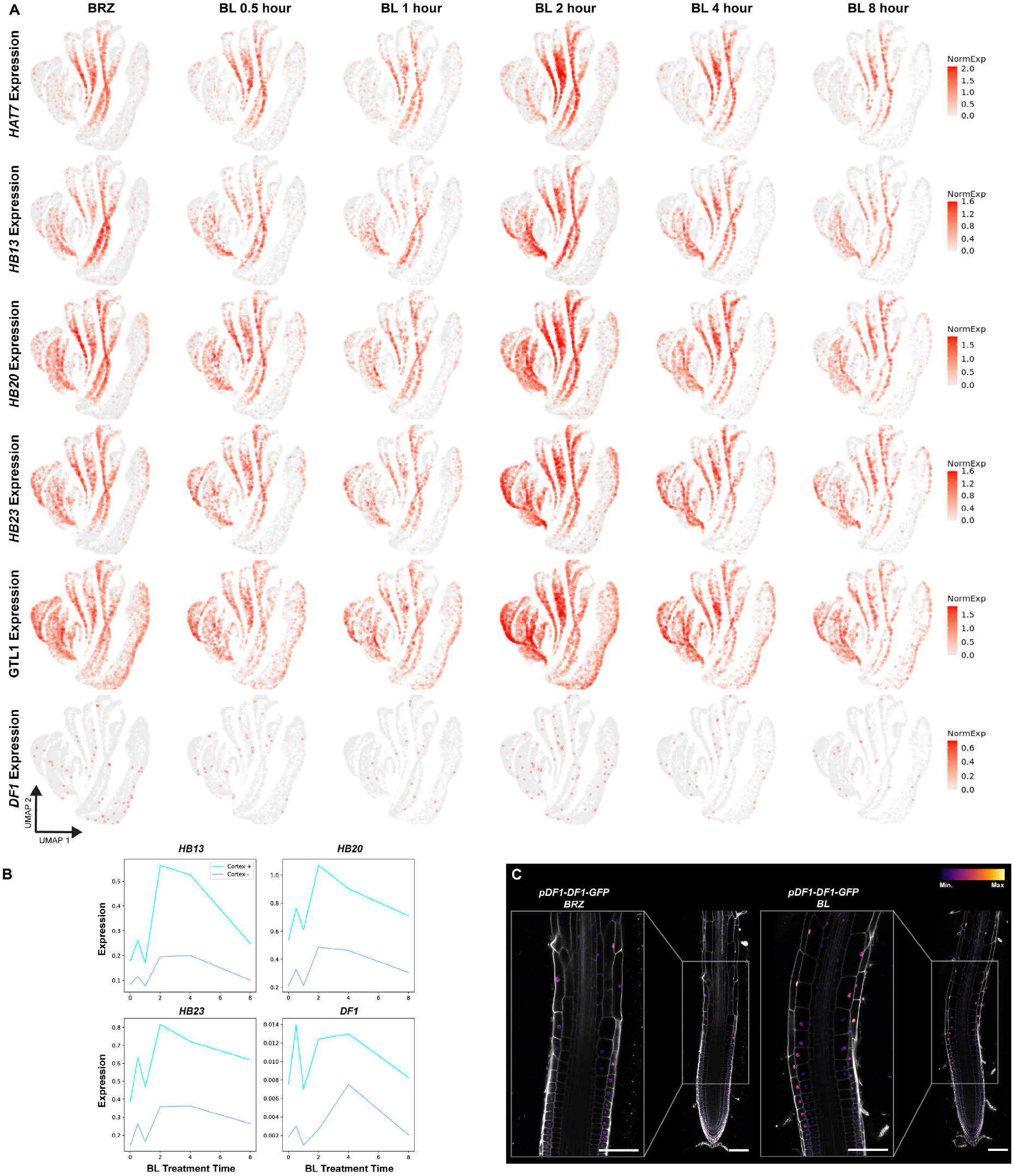
*HAT7* and *GTL1* family TFs are BR responsive regulators along cortex trajectories. **(A)** UMAP projections showing expression levels of HAT7 and GTL1 family transcription factors over the BR time series scRNA-seq experiment. The color scale represents log normalized, corrected UMI counts. **(B)** Expression trends for indicated transcription factors along WOT cortex cell wall + (cortex +) vs cortex cell wall - (cortex -) trajectories. **(C)** *pDF1-DF1-GFP* grown on 1 μM BRZ for 7 days and transferred to 1 μM BRZ or 100nM BL for 4 hours. Inset shows *DF1* signals in the elongating epidermis and cortex that increase with BL treatment. Propidium iodide-staining is shown in grey, with the color gradient indicating relative DF1-GFP levels. Scale bars, 100 μm.

**Fig. S11.**
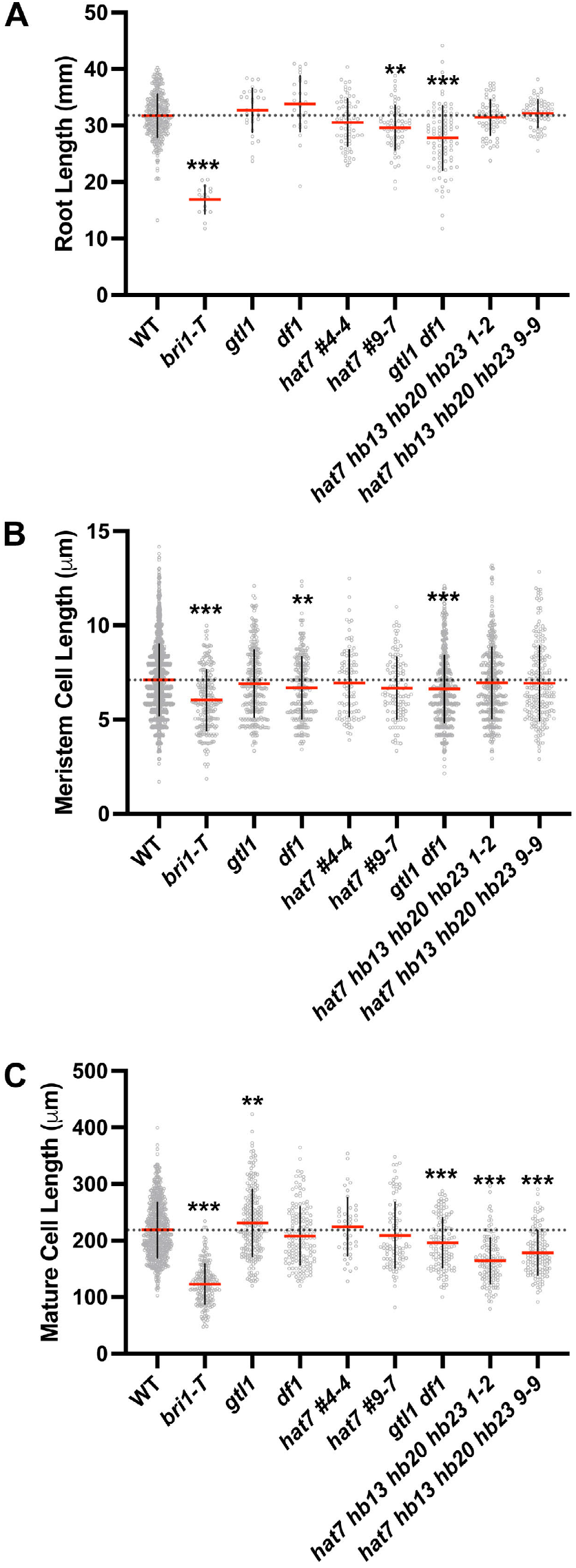
*HAT7* and *GTL1* family TFs affect cortex cell elongation. **(A)** Quantification of the root length in the indicated mutants. *hat7 hb13 hb20 hb23 1-2* and *hat7 hb13 hb20 hb23 9-9* represent two independent CRISPR mutants of. *hat7 hb13 hb20 hb23 1-2* is used as a representative allele throughout this study unless otherwise indicated. **(B)** Quantification of meristematic cortex cell length, defined as the first 20cells of individual roots starting from the quiescent center. **(C)** Quantification of mature cortex cell length. All individual data points are plotted. Red horizontal bars represent the means and error bars represent s.d. Significant differences between mutants and the wild-type control were determined by one-way ANOVA and Dunnett’s multiple comparisons tests. ***P<0.001, **P<0.01, *P<0.05.

**Fig. S12.**
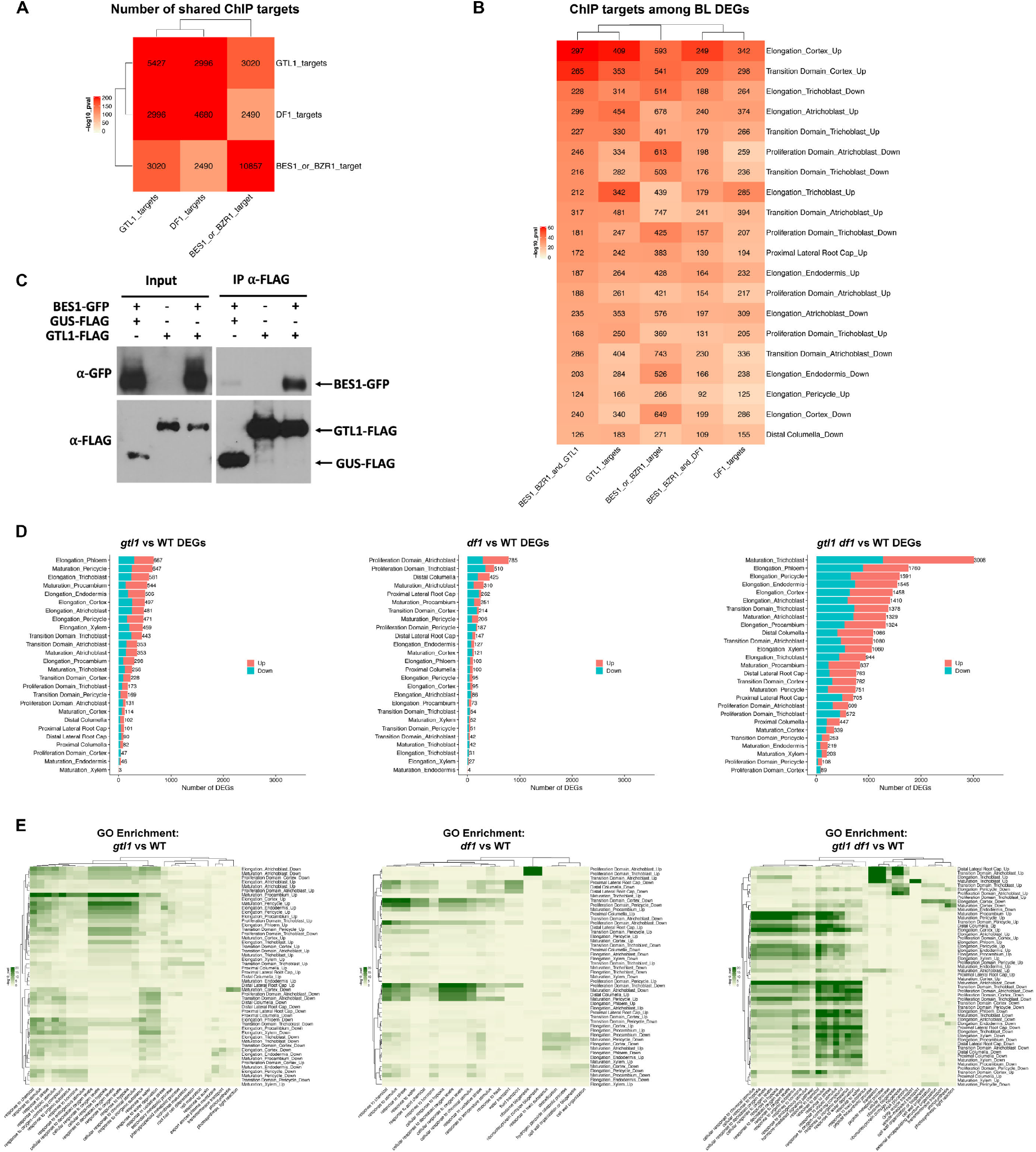
BES1 and GTL1 interact and share a common set of target genes. **(A)** Comparison of BES1 or BZR1, GTL1, and DF1 ChIP targets showing an overrepresentation of shared target genes. **(B)** Comparison of ChIP targets from (A) with BL 2 hour vs BRZ DEGs. The top 20 cell type/developmental stage combinations that are enriched for BES1 or BZR1 and GTL1 shared targets are shown. For (A) and (B) color represents log10 p-values from the indicated overlaps calculated from Fisher’s exact test by GeneOverlap. The number of genes in each intersection is indicated inside each box. **(C)** Co-Immunoprecipitation demonstrating BES1 interaction with GTL1. GTL1-FLAG immunoprecipitaed with anti-FLAG beads pulled down BES1-GFP, whereas a GUS-FLAG negative control did not. **(D)** Number of DEGs for each cell type/developmental stage combination in *gtl1, df1* or *gtl1 df1* scRNA-seq compared to WT. Color indicates the number of up-regulated vs down-regulated genes. **(E)** GO enrichment of DEGs in *gtl1, df1* or *gtl1 df1* scRNA-seq compared to WT.

## Notes

https://shiny.mdc-berlin.de/ARVEX/

### Competing Interest Statement

P.N.B. is the co-founder and Chair of the Scientific Advisory Board of Hi Fidelity Genetics, a company that works on crop root growth.

### Summary of Updates

This version of the manuscript was updated to streamline the narrative and figures.

## REFERENCES

1. E. Pierre-Jerome, C. Drapek, P. N. Benfey, Regulation of Division and Differentiation of Plant Stem Cells. Annu. Rev. Cell Dev. Biol. 34, 289–310 (2018).

2. R. Shahan, T. M. Nolan, P. N. Benfey, Single-cell analysis of cell identity in the Arabidopsis root apical meristem: insights and opportunities. J. Exp. Bot. (2021), doi:10.1093/jxb/erab228.

3. M. Levine, E. H. Davidson, Gene regulatory networks for development. Proc. Natl. Acad. Sci. U. S. A. 102, 4936–4942 (2005).

4. C. Seyfferth, J. Renema, J. R. Wendrich, T. Eekhout, R. Seurinck, N. Vandamme, B. Blob, Y. Saeys, Y. Helariutta, K. D. Birnbaum, B. De Rybel, Advances and Opportunities of Single-Cell Transcriptomics for Plant Research. Annu. Rev. Plant Biol. (2021), doi:10.1146/annurev-arplant-081720-010120.

5. W. Decaestecker, R. A. Buono, M. L. Pfeiffer, N. Vangheluwe, J. Jourquin, M. Karimi, G. Van Isterdael, T. Beeckman, M. K. Nowack, T. B. Jacobs, CRISPR-TSKO: A Technique for Efficient Mutagenesis in Specific Cell Types, Tissues, or Organs in Arabidopsis. Plant Cell. 31, 2868–2887 (2019).

6. S. D. Clouse, M. Langford, T. C. McMorris, A brassinosteroid-insensitive mutant in Arabidopsis thaliana exhibits multiple defects in growth and development. Plant Physiol. 111, 671–678 (1996).

7. Y. Hacham, N. Holland, C. Butterfield, S. Ubeda-Tomas, M. J. Bennett, J. Chory, S. Savaldi-Goldstein, Brassinosteroid perception in the epidermis controls root meristem size. Development. 138, 839–848 (2011).

8. T. M. Nolan, N. Vukašinović, D. Liu, E. Russinova, Y. Yin, Brassinosteroids: Multidimensional Regulators of Plant Growth, Development, and Stress Responses. Plant Cell. 32, 295–318 (2020).

9. A. Caño-Delgado, Y. Yin, C. Yu, D. Vafeados, S. Mora-García, J.-C. Cheng, K. H. Nam, J. Li, J. Chory, BRL1 and BRL3 are novel brassinosteroid receptors that function in vascular differentiation in Arabidopsis. Development. 131, 5341–5351 (2004).

10. T. Kinoshita, A. Caño-Delgado, H. Seto, S. Hiranuma, S. Fujioka, S. Yoshida, J. Chory, Binding of brassinosteroids to the extracellular domain of plant receptor kinase BRI1. Nature. 433, 167–171 (2005).

11. J. Li, J. Chory, A putative leucine-rich repeat receptor kinase involved in brassinosteroid signal transduction. Cell. 90, 929–938 (1997).

12. Y. Yin, Z.-Y. Wang, S. Mora-Garcia, J. Li, S. Yoshida, T. Asami, J. Chory, BES1 Accumulates in the Nucleus in Response to Brassinosteroids to Regulate Gene Expression and Promote Stem Elongation. Cell. 109, 181–191 (2002).

13. Z. Y. Wang, T. Nakano, J. Gendron, J. He, M. Chen, D. Vafeados, Y. Yang, S. Fujioka, S. Yoshida, T. Asami, J. Chory, Nuclear-localized BZR1 mediates brassinosteroid-induced growth and feedback suppression of brassinosteroid biosynthesis. Dev. Cell. 2, 505–513 (2002).

14. Y. Sun, X.-Y. Fan, D.-M. Cao, W. Tang, K. He, J.-Y. Zhu, J.-X. He, M.-Y. Bai, S. Zhu, E. Oh, S. Patil, T.-W. Kim, H. Ji, W. H. Wong, S. Y. Rhee, Z.-Y. Wang, Integration of brassinosteroid signal transduction with the transcription network for plant growth regulation in Arabidopsis. Dev. Cell. 19, 765–777 (2010).

15. X. Yu, L. Li, J. Zola, M. Aluru, H. Ye, A. Foudree, A brassinosteroid transcriptional network revealed by genome-wide identification of BESI target genes in Arabidopsis thaliana. The Plant Journal (2011) (available at https://onlinelibrary.wiley.com/doi/abs/10.1111/j.1365-313X.2010.04449.x).

16. N. M. Clark, T. M. Nolan, P. Wang, G. Song, C. Montes, C. T. Valentine, H. Guo, R. Sozzani, Y. Yin, J. W. Walley, Integrated omics networks reveal the temporal signaling events of brassinosteroid response in Arabidopsis. Nat. Commun. 12, 5858 (2021).

17. Y. Fridman, L. Elkouby, N. Holland, K. Vragović, R. Elbaum, S. Savaldi-Goldstein, Root growth is modulated by differential hormonal sensitivity in neighboring cells. Genes Dev. 28, 912–920 (2014).

18. M. Ackerman-Lavert, Y. Fridman, R. Matosevich, H. Khandal, L. Friedlander-Shani, K. Vragović, R. Ben El, G. Horev, D. Tarkowská, I. Efroni, S. Savaldi-Goldstein, Auxin requirements for a meristematic state in roots depend on a dual brassinosteroid function. Curr. Biol. (2021), doi:10.1016/j.cub.2021.07.075.

19. J. Vilarrasa-Blasi, M.-P. González-García, D. Frigola, N. Fàbregas, K. G. Alexiou, N. López-Bigas, S. Rivas, A. Jauneau, J. U. Lohmann, P. N. Benfey, M. Ibañes, A. I. Caño-Delgado, Regulation of plant stem cell quiescence by a brassinosteroid signaling module. Dev. Cell. 30, 36–47 (2014).

20. T. Asami, Y. K. Min, N. Nagata, K. Yamagishi, S. Takatsuto, S. Fujioka, N. Murofushi, I. Yamaguchi, S. Yoshida, Characterization of brassinazole, a triazole-type brassinosteroid biosynthesis inhibitor. Plant Physiol. 123, 93–100 (2000).

21. R. Shahan, C.-W. Hsu, T. M. Nolan, B. J. Cole, I. W. Taylor, L. Greenstreet, S. Zhang, A. Afanassiev, A. H. C. Vlot, G. Schiebinger, P. N. Benfey, U. Ohler, A single-cell Arabidopsis root atlas reveals developmental trajectories in wild-type and cell identity mutants. Dev. Cell (2022), doi:10.1016/j.devcel.2022.01.008.

22. V. B. Ivanov, J. G. Dubrovsky, Longitudinal zonation pattern in plant roots: conflicts and solutions. Trends Plant Sci. 18, 237–243 (2013).

23. T. Denyer, X. Ma, S. Klesen, E. Scacchi, K. Nieselt, M. C. P. Timmermans, Spatiotemporal Developmental Trajectories in the Arabidopsis Root Revealed Using High-Throughput Single-Cell RNA Sequencing. Dev. Cell. 48, 840–852.e5 (2019).

24. J. Chaiwanon, Z.-Y. Wang, Spatiotemporal brassinosteroid signaling and antagonism with auxin pattern stem cell dynamics in Arabidopsis roots. Curr. Biol. 25, 1031–1042 (2015).

25. K. Vragović, A. Sela, L. Friedlander-Shani, Y. Fridman, Y. Hacham, N. Holland, E. Bartom, T. C. Mockler, S. Savaldi-Goldstein, Translatome analyses capture of opposing tissue-specific brassinosteroid signals orchestrating root meristem differentiation. Proceedings of the National Academy of Sciences. 112, 923–928 (2015).

26. H. Guo, L. Li, M. Aluru, S. Aluru, Y. Yin, Mechanisms and networks for brassinosteroid regulated gene expression. Curr. Opin. Plant Biol. 16, 545–553 (2013).

27. M.-P. González-García, J. Vilarrasa-Blasi, M. Zhiponova, F. Divol, S. Mora-García, E. Russinova, A. I. Caño-Delgado, Brassinosteroids control meristem size by promoting cell cycle progression in Arabidopsis roots. Development. 138, 849–859 (2011).

28. P. E. Verslues, T. Longkumer, Size and activity of the root meristem: a key for drought resistance and a key model of drought-related signaling. Physiol. Plant. 174, e13622 (2022).

29. D. Dietrich, L. Pang, A. Kobayashi, J. A. Fozard, V. Boudolf, R. Bhosale, R. Antoni, T. Nguyen, S. Hiratsuka, N. Fujii, Y. Miyazawa, T.-W. Bae, D. M. Wells, M. R. Owen, L. R. Band, R. J. Dyson, O. E. Jensen, J. R. King, S. R. Tracy, C. J. Sturrock, S. J. Mooney, J. A. Roberts, R. P. Bhalerao, J. R. Dinneny, P. L. Rodriguez, A. Nagatani, Y. Hosokawa, T. I. Baskin, T. P. Pridmore, L. De Veylder, H. Takahashi, M. J. Bennett, Root hydrotropism is controlled via a cortex-specific growth mechanism. Nat Plants. 3, 17057 (2017).

30. L. Xie, C. Yang, X. Wang, Brassinosteroids can regulate cellulose biosynthesis by controlling the expression of CESA genes in Arabidopsis. J. Exp. Bot. 62, 4495–4506 (2011).

31. G. Schiebinger, J. Shu, M. Tabaka, B. Cleary, V. Subramanian, A. Solomon, J. Gould, S. Liu, S. Lin, P. Berube, L. Lee, J. Chen, J. Brumbaugh, P. Rigollet, K. Hochedlinger, R. Jaenisch, A. Regev, E. S. Lander, Optimal-Transport Analysis of Single-Cell Gene Expression Identifies Developmental Trajectories in Reprogramming. Cell. 176 (2019), p. 1517.

32. Y. Yin, D. Vafeados, Y. Tao, S. Yoshida, T. Asami, J. Chory, A new class of transcription factors mediates brassinosteroid-regulated gene expression in Arabidopsis. Cell. 120, 249–259 (2005).

33. M. K. Zhiponova, K. Morohashi, I. Vanhoutte, K. Machemer-Noonan, M. Revalska, M. Van Montagu, E. Grotewold, E. Russinova, Helix-loop-helix/basic helix-loop-helix transcription factor network represses cell elongation in Arabidopsis through an apparent incoherent feed-forward loop. Proc. Natl. Acad. Sci. U. S. A. 111, 2824–2829 (2014).

34. M. A. Moreno-Risueno, R. Sozzani, G. G. Yardımcı, J. J. Petricka, T. Vernoux, I. Blilou, J. Alonso, C. M. Winter, U. Ohler, B. Scheres, P. N. Benfey, Transcriptional control of tissue formation throughout root development. Science. 350, 426–430 (2015).

35. E. Henriksson, A. S. B. Olsson, H. Johannesson, H. Johansson, J. Hanson, P. Engström, E. Söderman, Homeodomain leucine zipper class I genes in Arabidopsis. Expression patterns and phylogenetic relationships. Plant Physiol. 139, 509–518 (2005).

36. J. Mattsson, E. Söderman, M. Svenson, C. Borkird, P. Engström, A new homeobox-leucine zipper gene from Arabidopsis thaliana. Plant Mol. Biol. 18, 1019–1022 (1992).

37. N. Vukašinović, Y. Wang, I. Vanhoutte, M. Fendrych, B. Guo, M. Kvasnica, P. Jiroutová, J. Oklestkova, M. Strnad, E. Russinova, Local brassinosteroid biosynthesis enables optimal root growth. Nat Plants. 7, 619–632 (2021).

38. D. M. Friedrichsen, C. A. Joazeiro, J. Li, T. Hunter, J. Chory, Brassinosteroid-insensitive-1 is a ubiquitously expressed leucine-rich repeat receptor serine/threonine kinase. Plant Physiol. 123, 1247–1256 (2000).

39. Z. He, Z. Y. Wang, J. Li, Q. Zhu, C. Lamb, P. Ronald, J. Chory, Perception of brassinosteroids by the extracellular domain of the receptor kinase BRI1. Science. 288, 2360–2363 (2000).

40. M. Graeff, S. Rana, J. R. Wendrich, J. Dorier, T. Eekhout, A. C. A. Fandino, N. Guex, G. W. Bassel, B. De Rybel, C. S. Hardtke, A single-cell morpho-transcriptomic map of brassinosteroid action in the Arabidopsis root. Mol. Plant. 0 (2021), doi:10.1016/j.molp.2021.07.021.

41. Y. Fridman, S. Strauss, G. Horev, M. Ackerman-Lavert, A. Reiner-Benaim, B. Lane, R. S. Smith, S. Savaldi-Goldstein, The root meristem is shaped by brassinosteroid control of cell geometry. Nature Plants. 7, 1475–1484 (2021).

42. Y. H. Kang, A. Breda, C. S. Hardtke, Brassinosteroid signaling directs formative cell divisions and protophloem differentiation in Arabidopsis root meristems. Development. 144, 272–280 (2017).

43. M. Graeff, S. Rana, P. Marhava, B. Moret, C. S. Hardtke, Local and Systemic Effects of Brassinosteroid Perception in Developing Phloem. Curr. Biol. (2020), doi:10.1016/j.cub.2020.02.029.

44. F. Bou Daher, Y. Chen, B. Bozorg, J. Clough, H. Jönsson, S. A. Braybrook, Anisotropic growth is achieved through the additive mechanical effect of material anisotropy and elastic asymmetry. Elife. 7 (2018), doi:10.7554/eLife.38161.

45. Baskin, Jensen, On the role of stress anisotropy in the growth of stems. J. Exp. Bot. (available at https://academic.oup.com/jxb/article-abstract/64/15/4697/460841).

46. E. Oh, J.-Y. Zhu, M.-Y. Bai, R. A. Arenhart, Y. Sun, Z.-Y. Wang, Cell elongation is regulated through a central circuit of interacting transcription factors in the Arabidopsis hypocotyl. Elife. 3 (2014), doi:10.7554/eLife.03031.

47. R. Seyed Rahmani, T. Shi, D. Zhang, X. Gou, J. Yi, G. Miclotte, K. Marchal, J. Li, Genome-wide expression and network analyses of mutants in key brassinosteroid signaling genes. BMC Genomics. 22, 465 (2021).

48. K. Kamimoto, C. M. Hoffmann, S. A. Morris, CellOracle: Dissecting cell identity via network inference and in silico gene perturbation. bioRxiv (2020), p. 2020.02.17.947416.

49. M. Shibata, C. Breuer, A. Kawamura, N. M. Clark, B. Rymen, L. Braidwood, K. Morohashi, W. Busch, P. N. Benfey, R. Sozzani, K. Sugimoto, GTL1 and DF1 regulate root hair growth through transcriptional repression of ROOT HAIR DEFECTIVE 6-LIKE 4 in Arabidopsis. Development. 145 (2018), doi:10.1242/dev.159707.

50. A. Gupta, A. Rico-Medina, A. I. Caño-Delgado, The physiology of plant responses to drought. Science. 368, 266–269 (2020).

51. B. O. R. Bargmann, S. Vanneste, G. Krouk, T. Nawy, I. Efroni, E. Shani, G. Choe, J. Friml, D. C. Bergmann, M. Estelle, K. D. Birnbaum, A map of cell type-specific auxin responses. Mol. Syst. Biol. 9, 688 (2013).

52. E. Shani, R. Weinstain, Y. Zhang, C. Castillejo, E. Kaiserli, J. Chory, R. Y. Tsien, M. Estelle, Gibberellins accumulate in the elongating endodermal cells of Arabidopsis root. Proc. Natl. Acad. Sci. U. S. A. 110, 4834–4839 (2013).

53. S. Ubeda-Tomás, R. Swarup, J. Coates, K. Swarup, L. Laplaze, G. T. S. Beemster, P. Hedden, R. Bhalerao, M. J. Bennett, Root growth in Arabidopsis requires gibberellin/DELLA signalling in the endodermis. Nat. Cell Biol. 10, 625–628 (2008).

54. Y. Geng, R. Wu, C. W. Wee, F. Xie, X. Wei, P. M. Y. Chan, C. Tham, L. Duan, J. R. Dinneny, A spatio-temporal understanding of growth regulation during the salt stress response in Arabidopsis. Plant Cell. 25, 2132–2154 (2013).

55. Y. Jaillais, Y. Belkhadir, E. Balsemão-Pires, J. L. Dangl, J. Chory, Extracellular leucine-rich repeats as a platform for receptor/coreceptor complex formation. Proc. Natl. Acad. Sci. U. S. A. 108, 8503–8507 (2011).

56. N. G. Irani, S. Di Rubbo, E. Mylle, J. Van den Begin, J. Schneider-Pizoń, J. Hniliková, M. Šíša, D. Buyst, J. Vilarrasa-Blasi, A.-M. Szatmári, D. Van Damme, K. Mishev, M.-C. Codreanu, L. Kohout, M. Strnad, A. I. Caño-Delgado, J. Friml, A. Madder, E. Russinova, Fluorescent castasterone reveals BRI1 signaling from the plasma membrane. Nat. Chem. Biol. 8, 583–589 (2012).

57. T. L. Shimada, T. Shimada, I. Hara-Nishimura, A rapid and non-destructive screenable marker, FAST, for identifying transformed seeds of Arabidopsis thaliana. Plant J. 61, 519–528 (2010).

58. J. Stuttmann, K. Barthel, P. Martin, J. Ordon, J. L. Erickson, R. Herr, F. Ferik, C. Kretschmer, T. Berner, J. Keilwagen, S. Marillonnet, U. Bonas, Highly efficient multiplex editing: one-shot generation of 8× Nicotiana benthamiana and 12× Arabidopsis mutants. Plant J. 106, 8–22 (2021).

59. C. Engler, M. Youles, R. Gruetzner, T.-M. Ehnert, S. Werner, J. D. G. Jones, N. J. Patron, S. Marillonnet, A Golden Gate Modular Cloning Toolbox for Plants. ACS Synth. Biol. 3, 839–843 (2014).

60. A. I. Andreou, N. Nakayama, Mobius Assembly: A versatile Golden-Gate framework towards universal DNA assembly. PLoS One. 13, e0189892 (2018).

61. S. J. Clough, A. F. Bent, Floral dip: a simplified method forAgrobacterium-mediated transformation ofArabidopsis thaliana. Plant J. 16, 735–743 (1998).

62. R. Grützner, P. Martin, C. Horn, S. Mortensen, E. J. Cram, C. W. T. Lee-Parsons, J. Stuttmann, S. Marillonnet, High-efficiency genome editing in plants mediated by a Cas9 gene containing multiple introns. Plant Communications. 2, 100135 (2021).

63. K. Labun, T. G. Montague, M. Krause, Y. N. Torres Cleuren, H. Tjeldnes, E. Valen, CHOPCHOP v3: expanding the CRISPR web toolbox beyond genome editing. Nucleic Acids Res. 47, W171–W174 (2019).

64. D. Conant, T. Hsiau, N. Rossi, J. Oki, T. Maures, K. Waite, J. Yang, S. Joshi, R. Kelso, K. Holden, B. L. Enzmann, R. Stoner, Inference of CRISPR Edits from Sanger Trace Data. CRISPR J. 5, 123–130 (2022).

65. Z. Feng, B. Zhang, W. Ding, X. Liu, D.-L. Yang, P. Wei, F. Cao, S. Zhu, F. Zhang, Y. Mao, J.-K. Zhu, Efficient genome editing in plants using a CRISPR/Cas system. Cell Res. 23, 1229–1232 (2013).

66. A. Houbaert, C. Zhang, M. Tiwari, K. Wang, A. de Marcos Serrano, D. V. Savatin, M. J. Urs, M. K. Zhiponova, G. E. Gudesblat, I. Vanhoutte, D. Eeckhout, S. Boeren, M. Karimi, C. Betti, T. Jacobs, C. Fenoll, M. Mena, S. de Vries, G. De Jaeger, E. Russinova, POLAR-guided signalling complex assembly and localization drive asymmetric cell division. Nature. 563, 574–578 (2018).

67. X. Wang, L. Ye, M. Lyu, R. Ursache, A. Löytynoja, A. P. Mähönen, An inducible genome editing system for plants. Nat Plants. 6, 766–772 (2020).

68. M. D. M. Marquès-Bueno, A. K. Morao, A. Cayrel, M. P. Platre, M. Barberon, E. Caillieux, V. Colot, Y. Jaillais, F. Roudier, G. Vert, A versatile Multisite Gateway-compatible promoter and transgenic line collection for cell type-specific functional genomics in Arabidopsis. Plant J. 85, 320–333 (2016).

69. R. Vanholme, I. Cesarino, K. Rataj, Y. Xiao, L. Sundin, G. Goeminne, H. Kim, J. Cross, K. Morreel, P. Araujo, L. Welsh, J. Haustraete, C. McClellan, B. Vanholme, J. Ralph, G. G. Simpson, C. Halpin, W. Boerjan, Caffeoyl shikimate esterase (CSE) is an enzyme in the lignin biosynthetic pathway in Arabidopsis. Science. 341, 1103–1106 (2013).

70. J. Schindelin, I. Arganda-Carreras, E. Frise, V. Kaynig, M. Longair, T. Pietzsch, S. Preibisch, C. Rueden, S. Saalfeld, B. Schmid, J.-Y. Tinevez, D. J. White, V. Hartenstein, K. Eliceiri, P. Tomancak, A. Cardona, Fiji: an open-source platform for biological-image analysis. Nat. Methods. 9, 676–682 (2012).

71. D. von Wangenheim, R. Hauschild, M. Fendrych, V. Barone, E. Benková, J. Friml, Live tracking of moving samples in confocal microscopy for vertically grown roots. Elife. 6 (2017), doi:10.7554/eLife.26792.

72. Z. Xie, T. Nolan, H. Jiang, B. Tang, M. Zhang, Z. Li, Y. Yin, The AP2/ERF Transcription Factor TINY Modulates Brassinosteroid-Regulated Plant Growth and Drought Responses in Arabidopsis. Plant Cell. 31, 1788–1806 (2019).

73. T. Nakagawa, T. Suzuki, S. Murata, S. Nakamura, T. Hino, K. Maeo, R. Tabata, T. Kawai, K. Tanaka, Y. Niwa, Y. Watanabe, K. Nakamura, T. Kimura, S. Ishiguro, Improved Gateway binary vectors: high-performance vectors for creation of fusion constructs in transgenic analysis of plants. Biosci. Biotechnol. Biochem. 71, 2095–2100 (2007).

74. S. S. Gampala, T.-W. Kim, J.-X. He, W. Tang, Z. Deng, M.-Y. Bai, S. Guan, S. Lalonde, Y. Sun, J. M. Gendron, H. Chen, N. Shibagaki, R. J. Ferl, D. Ehrhardt, K. Chong, A. L. Burlingame, Z.-Y. Wang, An essential role for 14-3-3 proteins in brassinosteroid signal transduction in Arabidopsis. Dev. Cell. 13, 177–189 (2007).

75. H. Ryu, K. Kim, H. Cho, J. Park, S. Choe, I. Hwang, Nucleocytoplasmic shuttling of BZR1 mediated by phosphorylation is essential in Arabidopsis brassinosteroid signaling. Plant Cell. 19, 2749–2762 (2007).

76. N. L. Bray, H. Pimentel, P. Melsted, L. Pachter, Near-optimal probabilistic RNA-seq quantification. Nat. Biotechnol. 34, 525–527 (2016).

77. P. Melsted, V. Ntranos, L. Pachter, The barcode, UMI, set format and BUStools. Bioinformatics. 35, 4472–4473 (2019).

78. P. Melsted, A. S. Booeshaghi, L. Liu, F. Gao, L. Lu, K. H. J. Min, E. da Veiga Beltrame, K. E. Hjörleifsson, J. Gehring, L. Pachter, Modular, efficient and constant-memory single-cell RNA-seq preprocessing. Nat. Biotechnol. 39, 813–818 (2021).

79. C.-W. Hsu, R. Shahan, T. M. Nolan, P. N. Benfey, U. Ohler, Protocol for fast scRNA-seq raw data processing using scKB and non-arbitrary quality control with COPILOT. STAR Protoc. 3, 101729 (2022).

80. C. S. McGinnis, L. M. Murrow, Z. J. Gartner, DoubletFinder: Doublet Detection in Single-Cell RNA Sequencing Data Using Artificial Nearest Neighbors. Cell Syst. 8, 329–337.e4 (2019).

81. C. Hafemeister, R. Satija, Normalization and variance stabilization of single-cell RNA-seq data using regularized negative binomial regression. Genome Biol. 20, 296 (2019).

82. S. M. Brady, D. A. Orlando, J.-Y. Lee, J. Y. Wang, J. Koch, J. R. Dinneny, D. Mace, U. Ohler, P. N. Benfey, A high-resolution root spatiotemporal map reveals dominant expression patterns. Science. 318, 801–806 (2007).

83. S. Li, M. Yamada, X. Han, U. Ohler, P. N. Benfey, High-Resolution Expression Map of the Arabidopsis Root Reveals Alternative Splicing and lincRNA Regulation. Dev. Cell. 39, 508–522 (2016).

84. R. Bhosale, V. Boudolf, F. Cuevas, R. Lu, T. Eekhout, Z. Hu, G. Van Isterdael, G. M. Lambert, F. Xu, M. K. Nowack, R. S. Smith, I. Vercauteren, R. De Rycke, V. Storme, T. Beeckman, J. C. Larkin, A. Kremer, H. Höfte, D. W. Galbraith, R. P. Kumpf, S. Maere, L. De Veylder, A Spatiotemporal DNA Endoploidy Map of the Arabidopsis Root Reveals Roles for the Endocycle in Root Development and Stress Adaptation. Plant Cell. 30, 2330–2351 (2018).

85. R. Heidstra, D. Welch, B. Scheres, Mosaic analyses using marked activation and deletion clones dissect Arabidopsis SCARECROW action in asymmetric cell division. Genes Dev. 18, 1964–1969 (2004).

86. J.-Y. Lee, J. Colinas, J. Y. Wang, D. Mace, U. Ohler, P. N. Benfey, Transcriptional and posttranscriptional regulation of transcription factor expression in Arabidopsis roots. Proc. Natl. Acad. Sci. U. S. A. 103, 6055–6060 (2006).

87. F. Baluska, S. Mancuso, D. Volkmann, P. W. Barlow, Root apex transition zone: a signalling-response nexus in the root. Trends Plant Sci. 15, 402–408 (2010).

88. T. Stuart, A. Butler, P. Hoffman, C. Hafemeister, E. Papalexi, W. M. Mauck 3rd, Y. Hao, M. Stoeckius, P. Smibert, R. Satija, Comprehensive Integration of Single-Cell Data. Cell. 177, 1888–1902.e21 (2019).

89. A. Butler, P. Hoffman, P. Smibert, E. Papalexi, R. Satija, Integrating single-cell transcriptomic data across different conditions, technologies, and species. Nat. Biotechnol. 36, 411–420 (2018).

90. R. A. Amezquita, A. T. L. Lun, E. Becht, V. J. Carey, L. N. Carpp, L. Geistlinger, F. Martini, K. Rue-Albrecht, D. Risso, C. Soneson, L. Waldron, H. Pagès, M. L. Smith, W. Huber, M. Morgan, R. Gottardo, S. C. Hicks, Orchestrating single-cell analysis with Bioconductor. Nat. Methods (2019), doi:10.1038/s41592-019-0654-x.

91. K. Van den Berge, H. Roux de Bézieux, K. Street, W. Saelens, R. Cannoodt, Y. Saeys, S. Dudoit, L. Clement, Trajectory-based differential expression analysis for single-cell sequencing data. Nat. Commun. 11, 1201 (2020).

92. H. L. Crowell, C. Soneson, P.-L. Germain, D. Calini, L. Collin, C. Raposo, D. Malhotra, M. D. Robinson, muscat detects subpopulation-specific state transitions from multi-sample multi-condition single-cell transcriptomics data. Nat. Commun. 11, 6077 (2020).

93. J. W. Squair, M. Gautier, C. Kathe, M. A. Anderson, N. D. James, T. H. Hutson, R. Hudelle, T. Qaiser, K. J. E. Matson, Q. Barraud, A. J. Levine, G. La Manno, M. A. Skinnider, G. Courtine, Confronting false discoveries in single-cell differential expression. Nat. Commun. 12, 5692 (2021).

94. D. J. McCarthy, Y. Chen, G. K. Smyth, Differential expression analysis of multifactor RNA-Seq experiments with respect to biological variation. Nucleic Acids Res. 40, 4288–4297 (2012).

95. L. Kolberg, U. Raudvere, I. Kuzmin, J. Vilo, H. Peterson, gprofiler2 -- an R package for gene list functional enrichment analysis and namespace conversion toolset g:Profiler. F1000Research. 9 (2020), p. 709.

96. Z. Gu, R. Eils, M. Schlesner, Complex heatmaps reveal patterns and correlations in multidimensional genomic data. Bioinformatics. 32, 2847–2849 (2016).

97. H. Wickham, ggplot2: Elegant Graphics for Data Analysis (2016), (available at https://ggplot2.tidyverse.org).

98. R. Rahni, K. D. Birnbaum, Week-long imaging of cell divisions in the Arabidopsis root meristem. Plant Methods. 15, 30 (2019).

99. S. Zhang, A. Afanassiev, L. Greenstreet, T. Matsumoto, G. Schiebinger, Optimal transport analysis reveals trajectories in steady-state systems. PLoS Comput. Biol. 17, e1009466 (2021).

100. A. Farmer, S. Thibivilliers, K. H. Ryu, J. Schiefelbein, M. Libault, Single-nucleus RNA and ATAC sequencing reveals the impact of chromatin accessibility on gene expression in Arabidopsis roots at the single-cell level. Mol. Plant (2021), doi:10.1016/j.molp.2021.01.001.

101. H. A. Pliner, J. S. Packer, J. L. McFaline-Figueroa, D. A. Cusanovich, R. M. Daza, D. Aghamirzaie, S. Srivatsan, X. Qiu, D. Jackson, A. Minkina, A. C. Adey, F. J. Steemers, J. Shendure, C. Trapnell, Cicero Predicts cis-Regulatory DNA Interactions from Single-Cell Chromatin Accessibility Data. Mol. Cell. 71, 858–871.e8 (2018).

102. A. Bartlett, R. C. O’Malley, S.-S. C. Huang, M. Galli, J. R. Nery, A. Gallavotti, J. R. Ecker, Mapping genome-wide transcription-factor binding sites using DAP-seq. Nat. Protoc. 12, 1659–1672 (2017).

103. I. De Clercq, J. Van de Velde, X. Luo, L. Liu, V. Storme, M. Van Bel, R. Pottie, D. Vaneechoutte, F. Van Breusegem, K. Vandepoele, Integrative inference of transcriptional networks in Arabidopsis yields novel ROS signalling regulators. Nature Plants. 7, 500–513 (2021).

104. J. L. Pruneda-Paz, G. Breton, D. H. Nagel, S. E. Kang, K. Bonaldi, C. J. Doherty, S. Ravelo, M. Galli, J. R. Ecker, S. A. Kay, A genome-scale resource for the functional characterization of Arabidopsis transcription factors. Cell Rep. 8, 622–632 (2014).

105. T. Kamiya, M. Borghi, P. Wang, J. M. C. Danku, L. Kalmbach, P. S. Hosmani, S. Naseer, T. Fujiwara, N. Geldner, D. E. Salt, The MYB36 transcription factor orchestrates Casparian strip formation. Proc. Natl. Acad. Sci. U. S. A. 112, 10533–10538 (2015).

106. T. Bennett, A. van den Toorn, G. F. Sanchez-Perez, A. Campilho, V. Willemsen, B. Snel, B. Scheres, SOMBRERO, BEARSKIN1, and BEARSKIN2 Regulate Root Cap Maturation in *Arabidopsis*. The Plant Cell. 22 (2010), pp. 640–654.

