## Supplementary material for "Brassinosteroid gene regulatory networks at cellular resolution": Data S1

Data S1: Summary of the scRNA-seq samples reported in this study

| Sample | Name | Genotype | Treatment | Date | 10X_chemistry | Number of cells after COPILOT filtering | Number of genes detected | Median UMI counts per cell | Median number of genes detected per cell |
|---|---|---|---|---|---|---|---|---|---|
| sc_1 | WT control | WT | Control | 20191214 | v3 | 9759 | 21844 | 3028 | 1528 |
| sc_2 | WT BRZ control | WT | BRZ | 20191214 | v3 | 7512 | 22403 | 6274 | 2497 |
| sc_5 | WT BRZ then 2 hour BL | WT | BL | 20191214 | v3 | 9790 | 22656 | 5415.5 | 2421.5 |
| sc_43 | WT BRZ control | WT | BRZ | 20200212 | v3 | 8874 | 22534 | 4936 | 2071 |
| sc_44 | WT BRZ then 30 mins BL | WT | BL | 20200212 | v3 | 6843 | 22046 | 6935 | 2441 |
| sc_45 | WT BRZ then 1 hour BL | WT | BL | 20200212 | v3 | 6336 | 22434 | 5525.5 | 2067 |
| sc_46 | WT BRZ then 2 hour BL | WT | BL | 20200212 | v3 | 6847 | 22490 | 7506 | 2632 |
| sc_47 | WT BRZ then 4 hour BL | WT | BL | 20200212 | v3 | 6907 | 22893 | 6430 | 2297 |
| sc_48 | WT BRZ then 8 hour BL | WT | BL | 20200212 | v3 | 6389 | 22873 | 5230 | 2014 |
| sc_49 | WT BRZ then 2 hour BL | WT | BL | 20200212 | v3 | 5638 | 22567 | 7764 | 2576 |
| sc_50 | WT BRZ control | WT | BRZ | 20200212 | v3 | 5087 | 22014 | 6800 | 2310 |
| sc_122 | WT | WT | Control | 20210804 | v3.1 | 11614 | 21930 | 6359 | 2377 |
| sc_123 | gtl1-1 | gtl1 | Control | 20210804 | v3.1 | 11141 | 22145 | 7475 | 2537 |
| sc_124 | df1-1 | df1 | Control | 20210804 | v3.1 | 9058 | 21751 | 6572.5 | 2275.5 |
| sc_125 | gtl1-1 df1-1 | gtl1_df1 | Control | 20210804 | v3.1 | 10012 | 22457 | 8517 | 2714 |
| sc_126 | WT | WT | Control | 20210804 | v3.1 | 7290 | 21885 | 9675 | 2822 |
| sc_127 | gtl1-1 | gtl1 | Control | 20210804 | v3.1 | 11453 | 22057 | 6277 | 2299 |
| sc_128 | df1-1 | df1 | Control | 20210804 | v3.1 | 6620 | 22152 | 11138.5 | 2978 |
| sc_129 | gtl1-1 df1-1 | gtl1_df1 | Control | 20210804 | v3.1 | 7622 | 22388 | 10281.5 | 3008 |
| sc_130 | WT | WT | Control | 20211001 | v3.1 | 6589 | 21554 | 10327 | 2983 |
| sc_131 | bri1-T | bri1_T | Control | 20211001 | v3.1 | 7621 | 22130 | 9461 | 3148 |
| sc_133 | pGL2:BRI1-GFP_bri1-T | pGL2_BRI1_GFP_bri1_T | Control | 20211001 | v3.1 | 5975 | 22518 | 11136 | 3370 |
| sc_134 | WT | WT | Control | 20211001 | v3.1 | 7745 | 22305 | 10538 | 3038 |
| sc_135 | bri1-T | bri1_T | Control | 20211001 | v3.1 | 5028 | 21909 | 11838 | 3550.5 |
| sc_137 | pGL2:BRI1-GFP_bri1-T | pGL2_BRI1_GFP_bri1_T | Control | 20211001 | v3.1 | 1903 | 21948 | 18497 | 4085 |

COPILOT summaries for each sample are below. Note that the cell number in each summary is before doublet removal.

### sc_1 Summary

Processed by COPILOT

Summary Analysis

#### Parameters

| | |
|---|---|
| Iteration of Filtering | 1 |
| Mitochondrial Expression Threshold | 5 % |
| Top High Quality Cell Filtered | 1 % |
| Doublet Removed | Yes |

#### UMI Counts Histogram

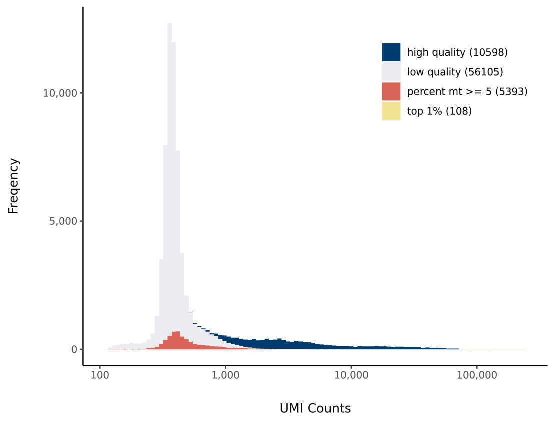


#### Cell Stats

| | |
|---|---|
| Estimated Number of High Quality Cell | 10,598 |
| High Quality Cell | 14.68 % |
| Total UMI Counts in High Quality Cell | 79,958,429 |
| UMI Counts in High Quality Cell | 67.1 % |
| Median UMI Counts per High Quality Cell | 3,079 |
| Median Genes per High Quality Cell | 1,526 |
| Total Genes Detected in High Quality Cell | 24,624 |
| Cell above Mitochondrial Expression Threshold | 7.47 % |
| Estimated Doublet Rate in High Quality Cell | 7.92 % |

#### Number of Genes Histogram

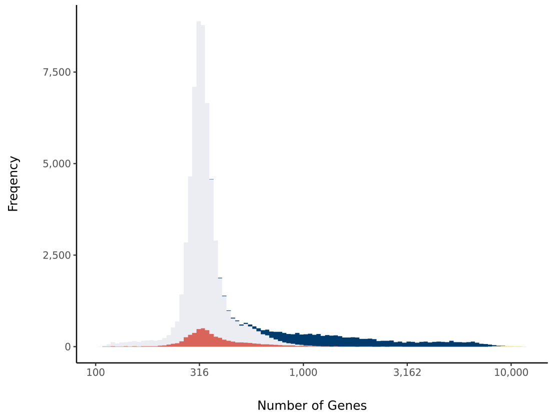


#### Sequencing Stats

| | |
|---|---|
| Number of Reads Processed | 255,953,705 |
| Reads Pseudoaligned | 92.4 % |
| Reads on Whitelist | 95.01 % |
| Total UMI Counts | 119,154,816 |
| Sequencing Technology | 10xv3 |
| Species | Arabidopsis thaliana |
| Transcriptome | TAIR10 |

#### Sample Stats

| | |
|---|---|
| Sample | sc_1 |
| Name | WT control |
| Source | Benfey lab |
| Genotype | WT Col-0 |
| Transgene | NA |
| Treatment | Untreated |
| Age | 7_day |
| Timepoint | 0 |
| Rep | 1 |
| Target Cells | 10,000 |
| Date | 2019-12-14 |
| Seq Run | Nolan_6131 |

#### Barcode Rank Plot

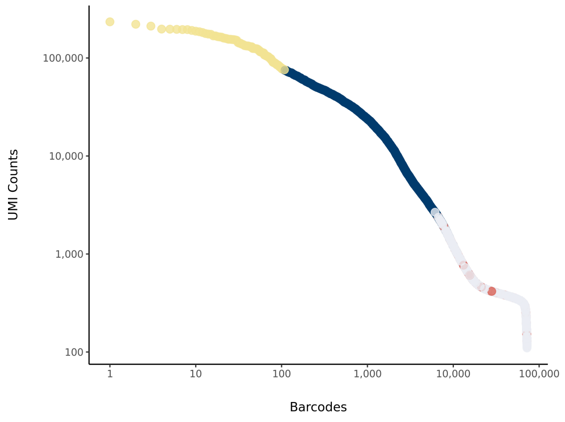

### sc_2 Summary

Processed by COPILOT



#### Parameters

| | |
|---|---|
| Iteration of Filtering | 1 |
| Mitochondrial Expression Threshold | 5 % |
| Top High Quality Cell Filtered | 1 % |
| Doublet Removed | Yes |

#### Cell Stats

| | |
|---|---|
| Estimated Number of High Quality Cell | 7,991 |
| High Quality Cell | 10.97 % |
| Total UMI Counts in High Quality Cell | 104,405,426 |
| UMI Counts in High Quality Cell | 64.57 % |
| Median UMI Counts per High Quality Cell | 6,476 |
| Median Genes per High Quality Cell | 2,552 |
| Total Genes Detected in High Quality Cell | 25,093 |
| Cell above Mitochondrial Expression Threshold | 9.96 % |
| Estimated Doublet Rate in High Quality Cell | 6 % |

#### Sequencing Stats

| | |
|---|---|
| Number of Reads Processed | 319,784,267 |
| Reads Pseudoaligned | 89.2 % |
| Reads on Whitelist | 93.58 % |
| Total UMI Counts | 161,702,644 |
| Sequencing Technology | 10xv3 |
| Species | Arabidopsis thaliana |
| Transcriptome | TAIR10 |

#### Sample Stats

| | |
|---|---|
| Sample | sc_2 |
| Name | WT BRZ |
| Source | Benfey lab |
| Genotype | WT Col-0 |
| Transgene | NA |
| Treatment | BRZ |
| Age | 7_day |
| Timepoint | 0 |
| Rep | 1 |
| Target Cells | 10,000 |
| Date | 2019-12-14 |
| Seq Run | Nolan_6131 |

#### UMI Counts Histogram

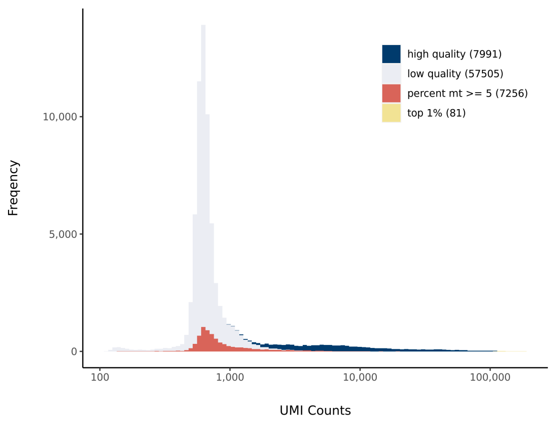


#### Number of Genes Histogram

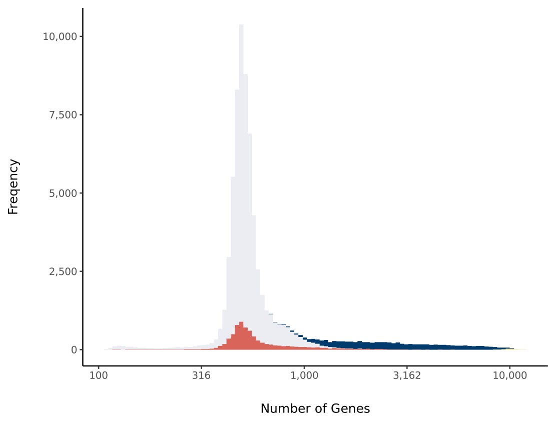


#### Barcode Rank Plot

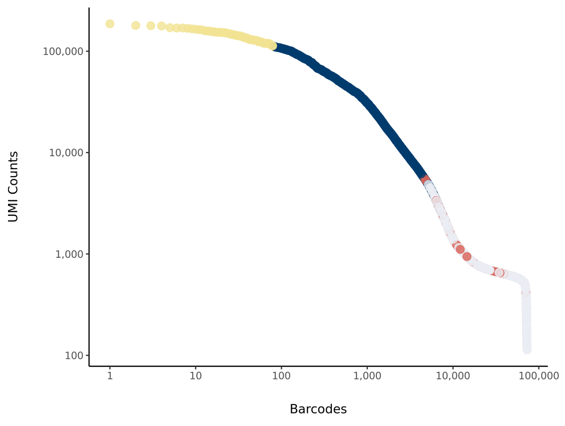

### sc_5 Summary

Processed by COPILOT



#### Parameters

| | |
|---|---|
| Iteration of Filtering | 1 |
| Mitochondrial Expression Threshold | 5 % |
| Top High Quality Cell Filtered | 1 % |
| Doublet Removed | Yes |

#### Cell Stats

| | |
|---|---|
| Estimated Number of High Quality Cell | 10,635 |
| High Quality Cell | 14.92 % |
| Total UMI Counts in High Quality Cell | 108,129,868 |
| UMI Counts in High Quality Cell | 59.16 % |
| Median UMI Counts per High Quality Cell | 5,699 |
| Median Genes per High Quality Cell | 2,498 |
| Total Genes Detected in High Quality Cell | 25,409 |
| Cell above Mitochondrial Expression Threshold | 5.16 % |
| Estimated Doublet Rate in High Quality Cell | 7.95 % |

#### Sequencing Stats

| | |
|---|---|
| Number of Reads Processed | 296,501,944 |
| Reads Pseudoaligned | 90.7 % |
| Reads on Whitelist | 94.02 % |
| Total UMI Counts | 182,772,760 |
| Sequencing Technology | 10xv3 |
| Species | Arabidopsis thaliana |
| Transcriptome | TAIR10 |

#### Sample Stats

| | |
|---|---|
| Sample | sc_5 |
| Name | WT BL |
| Source | Benfey lab |
| Genotype | WT Col-0 |
| Transgene | NA |
| Treatment | BL |
| Age | 7_day |
| Timepoint | 2 |
| Rep | 1 |
| Target Cells | 10,000 |
| Date | 2019-12-14 |
| Seq Run | Nolan_6131 |

#### UMI Counts Histogram

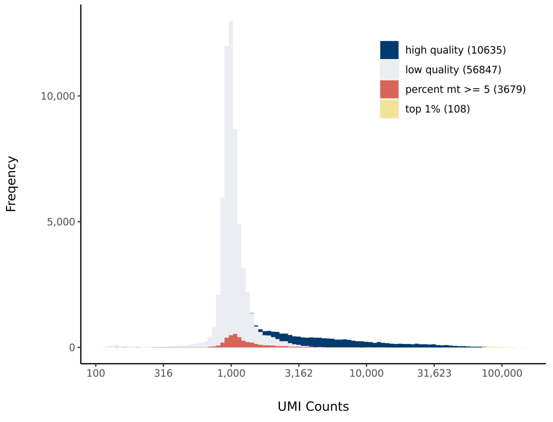


#### Number of Genes Histogram

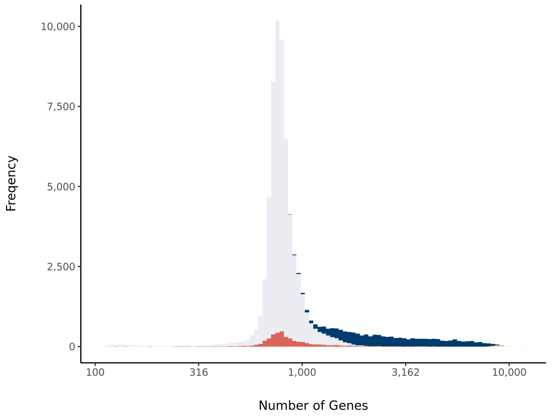


#### Barcode Rank Plot

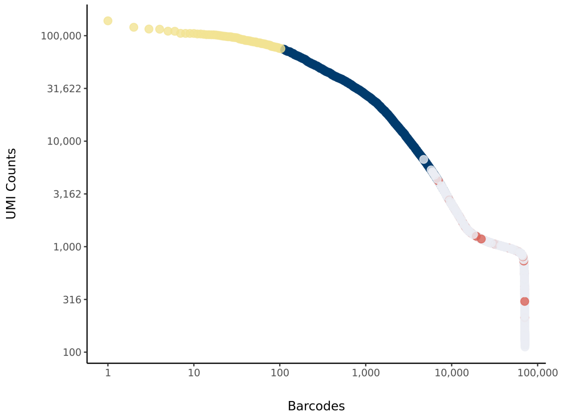

### sc_43 Summary

Processed by COPILOT

Summary Analysis

#### Parameters

| | |
|---|---|
| Iteration of Filtering | 1 |
| Mitochondrial Expression Threshold | 5 % |
| Top High Quality Cell Filtered | 1 % |
| Doublet Removed | Yes |

#### Cell Stats

| | |
|---|---|
| Estimated Number of High Quality Cell | 9,557 |
| High Quality Cell | 13.31 % |
| Total UMI Counts in High Quality Cell | 121,197,900 |
| UMI Counts in High Quality Cell | 79.25 % |
| Median UMI Counts per High Quality Cell | 4,683 |
| Median Genes per High Quality Cell | 1,998 |
| Total Genes Detected in High Quality Cell | 25,096 |
| Cell above Mitochondrial Expression Threshold | 28.11 % |
| Estimated Doublet Rate in High Quality Cell | 7.15 % |

#### Sequencing Stats

| | |
|---|---|
| Number of Reads Processed | 271,120,139 |
| Reads Pseudoaligned | 93.2 % |
| Reads on Whitelist | 96.34 % |
| Total UMI Counts | 152,927,724 |
| Sequencing Technology | 10xv3 |
| Species | Arabidopsis thaliana |
| Transcriptome | TAIR10 |

#### Sample Stats

| | |
|---|---|
| Sample | sc_43 |
| Name | WT BRZ control |
| Source | Benfey lab |
| Genotype | WT Col-0 |
| Transgene | NA |
| Treatment | BRZ |
| Age | 7_day |
| Timepoint | 0 |
| Rep | NA |
| Target Cells | 10,000 |
| Date | 2020-02-12 |
| Seq Run | Nolan_6199 (NextSeq); Nolan_6226 (NovaSeq S4) |

#### UMI Counts Histogram

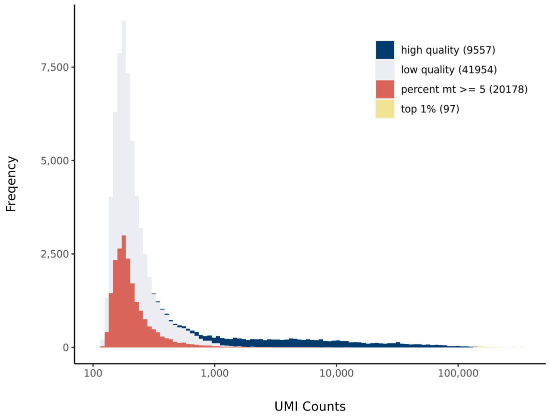


#### Number of Genes Histogram

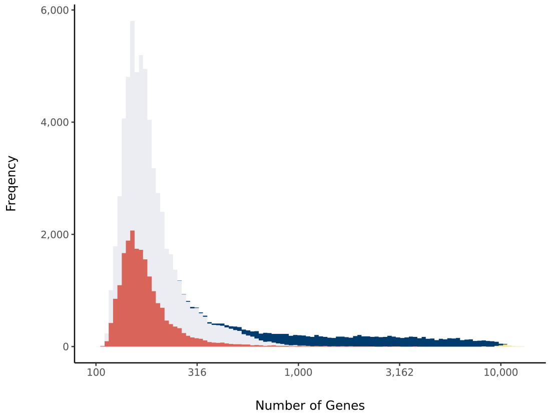


#### Barcode Rank Plot

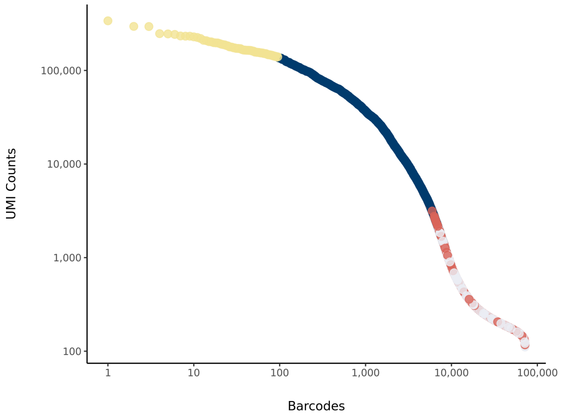

### sc_44 Summary

Processed by COPILOT

Summary Analysis

#### Parameters

| | |
|---|---|
| Iteration of Filtering | 1 |
| Mitochondrial Expression Threshold | 5 % |
| Top High Quality Cell Filtered | 1 % |
| Doublet Removed | Yes |

#### Cell Stats

| | |
|---|---|
| Estimated Number of High Quality Cell | 7,237 |
| High Quality Cell | 9.84 % |
| Total UMI Counts in High Quality Cell | 109,910,011 |
| UMI Counts in High Quality Cell | 77.94 % |
| Median UMI Counts per High Quality Cell | 7,331 |
| Median Genes per High Quality Cell | 2,529 |
| Total Genes Detected in High Quality Cell | 24,730 |
| Cell above Mitochondrial Expression Threshold | 2.92 % |
| Estimated Doublet Rate in High Quality Cell | 5.45 % |

#### Sequencing Stats

| | |
|---|---|
| Number of Reads Processed | 265,218,386 |
| Reads Pseudoaligned | 94.5 % |
| Reads on Whitelist | 96.68 % |
| Total UMI Counts | 141,019,844 |
| Sequencing Technology | 10xv3 |
| Species | Arabidopsis thaliana |
| Transcriptome | TAIR10 |

#### Sample Stats

| | |
|---|---|
| Sample | sc_44 |
| Name | WT BRZ then 30 mins BL |
| Source | Benfey lab |
| Genotype | WT Col-0 |
| Transgene | NA |
| Treatment | BL |
| Age | 7_day |
| Timepoint | 0.5 |
| Rep | NA |
| Target Cells | 10,000 |
| Date | 2020-02-12 |
| Seq Run | Nolan_6199 (NextSeq); Nolan_6226 (NovaSeq S4) |

#### UMI Counts Histogram

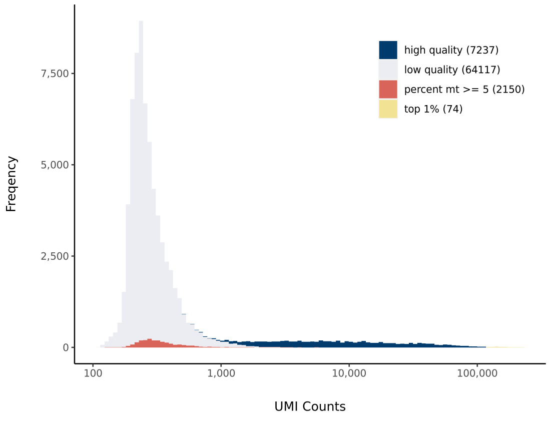


#### Number of Genes Histogram

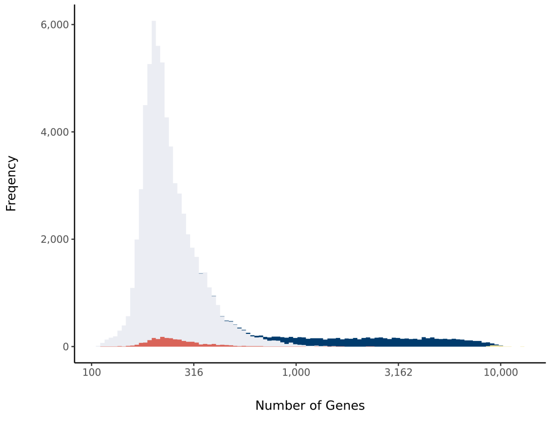


#### Barcode Rank Plot

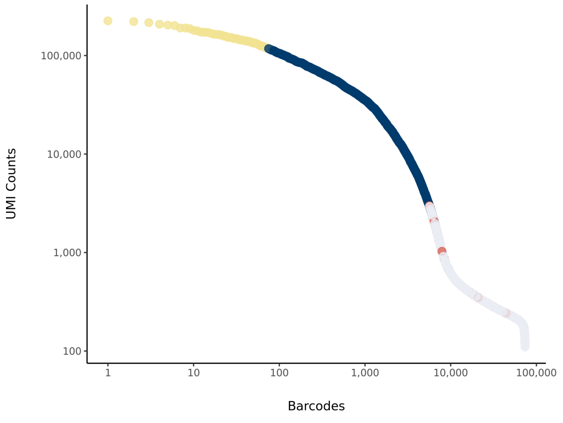

### sc_45 Summary

Processed by COPILOT



#### Parameters

| | |
|---|---|
| Iteration of Filtering | 1 |
| Mitochondrial Expression Threshold | 5 % |
| Top High Quality Cell Filtered | 1 % |
| Doublet Removed | Yes |

#### Cell Stats

| | |
|---|---|
| Estimated Number of High Quality Cell | 6,672 |
| High Quality Cell | 10.2 % |
| Total UMI Counts in High Quality Cell | 97,380,005 |
| UMI Counts in High Quality Cell | 77.07 % |
| Median UMI Counts per High Quality Cell | 5,285 |
| Median Genes per High Quality Cell | 1,993 |
| Total Genes Detected in High Quality Cell | 25,118 |
| Cell above Mitochondrial Expression Threshold | 41.9 % |
| Estimated Doublet Rate in High Quality Cell | 5.03 % |

#### Sequencing Stats

| | |
|---|---|
| Number of Reads Processed | 273,107,979 |
| Reads Pseudoaligned | 92.8 % |
| Reads on Whitelist | 96.2 % |
| Total UMI Counts | 126,354,735 |
| Sequencing Technology | 10xv3 |
| Species | Arabidopsis thaliana |
| Transcriptome | TAIR10 |

#### Sample Stats

| | |
|---|---|
| Sample | sc_45 |
| Name | WT BRZ then 1 hour BL |
| Source | Benfey lab |
| Genotype | WT Col-0 |
| Transgene | NA |
| Treatment | BL |
| Age | 7_day |
| Timepoint | 1 |
| Rep | NA |
| Target Cells | 10,000 |
| Date | 2020-02-12 |
| Seq Run | Nolan_6199 (NextSeq); Nolan_6226 (NovaSeq S4) |

#### UMI Counts Histogram

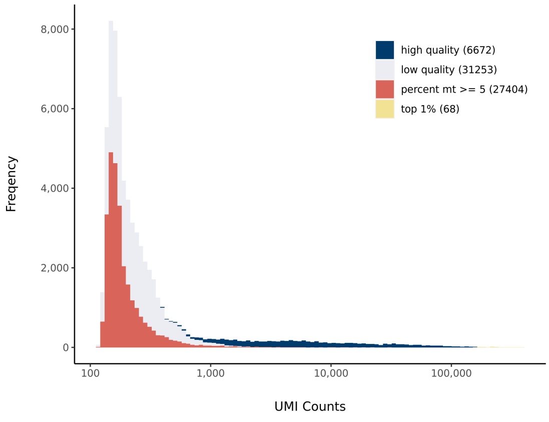


#### Number of Genes Histogram

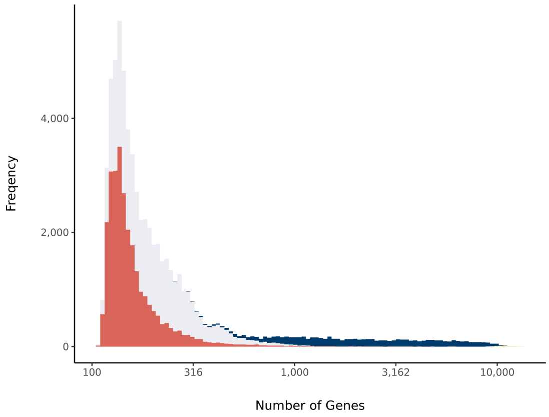


#### Barcode Rank Plot

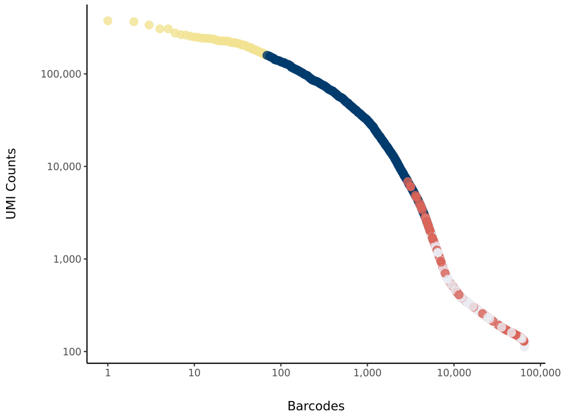

### sc_46 Summary

Processed by COPILOT



#### Parameters

| | |
|---|---|
| Iteration of Filtering | 1 |
| Mitochondrial Expression Threshold | 5 % |
| Top High Quality Cell Filtered | 1 % |
| Doublet Removed | Yes |

#### Cell Stats

| | |
|---|---|
| Estimated Number of High Quality Cell | 7,242 |
| High Quality Cell | 9.74 % |
| Total UMI Counts in High Quality Cell | 120,245,825 |
| UMI Counts in High Quality Cell | 77.74 % |
| Median UMI Counts per High Quality Cell | 7,205.5 |
| Median Genes per High Quality Cell | 2,565 |
| Total Genes Detected in High Quality Cell | 25,124 |
| Cell above Mitochondrial Expression Threshold | 4.73 % |
| Estimated Doublet Rate in High Quality Cell | 5.45 % |

#### Sequencing Stats

| | |
|---|---|
| Number of Reads Processed | 277,240,558 |
| Reads Pseudoaligned | 94.3 % |
| Reads on Whitelist | 96.57 % |
| Total UMI Counts | 154,680,730 |
| Sequencing Technology | 10xv3 |
| Species | Arabidopsis thaliana |
| Transcriptome | TAIR10 |

#### Sample Stats

| | |
|---|---|
| Sample | sc_46 |
| Name | WT BRZ then 2 hour BL |
| Source | Benfey lab |
| Genotype | WT Col-0 |
| Transgene | NA |
| Treatment | BL |
| Age | 7_day |
| Timepoint | 2 |
| Rep | NA |
| Target Cells | 10,000 |
| Date | 2020-02-12 |
| Seq Run | Nolan_6199 (NextSeq); Nolan_6226 (NovaSeq S4) |

#### UMI Counts Histogram

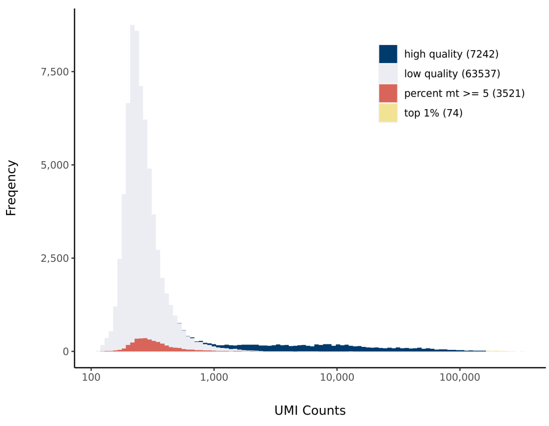


#### Number of Genes Histogram

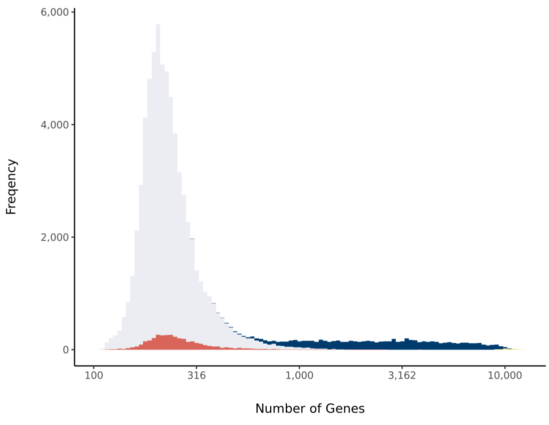


#### Barcode Rank Plot

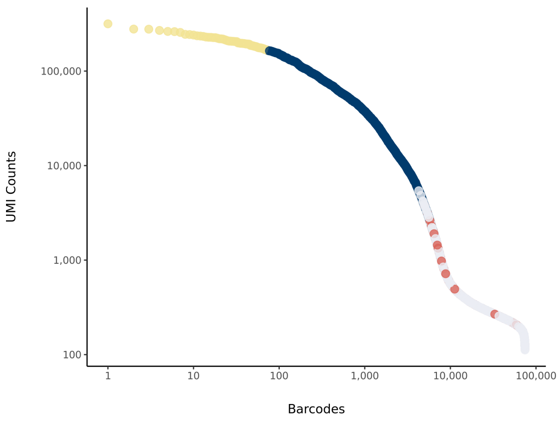

### sc_47 Summary

Processed by COPILOT

Summary | Analysis

#### Parameters

| | |
|---|---|
| Iteration of Filtering | 1 |
| Mitochondrial Expression Threshold | 5 % |
| Top High Quality Cell Filtered | 1 % |
| Doublet Removed | Yes |

#### Cell Stats

| | |
|---|---|
| Estimated Number of High Quality Cell | 7,309 |
| High Quality Cell | 10.8 % |
| Total UMI Counts in High Quality Cell | 103,642,465 |
| UMI Counts in High Quality Cell | 79.31 % |
| Median UMI Counts per High Quality Cell | 6,163 |
| Median Genes per High Quality Cell | 2,175 |
| Total Genes Detected in High Quality Cell | 25,556 |
| Cell above Mitochondrial Expression Threshold | 21.5 % |
| Estimated Doublet Rate in High Quality Cell | 5.5 % |

#### Sequencing Stats

| | |
|---|---|
| Number of Reads Processed | 291,507,817 |
| Reads Pseudoaligned | 93.7 % |
| Reads on Whitelist | 96.36 % |
| Total UMI Counts | 130,685,064 |
| Sequencing Technology | 10xv3 |
| Species | Arabidopsis thaliana |
| Transcriptome | TAIR10 |

#### Sample Stats

| | |
|---|---|
| Sample | sc_47 |
| Name | WT BRZ then 4 hour BL |
| Source | Benfey lab |
| Genotype | WT Col-0 |
| Transgene | NA |
| Treatment | BL |
| Age | 7_day |
| Timepoint | 4 |
| Rep | NA |
| Target Cells | 10,000 |
| Date | 2020-02-12 |
| Seq Run | Nolan_6199 (NextSeq); Nolan_6226 (NovaSeq S4) |

#### UMI Counts Histogram

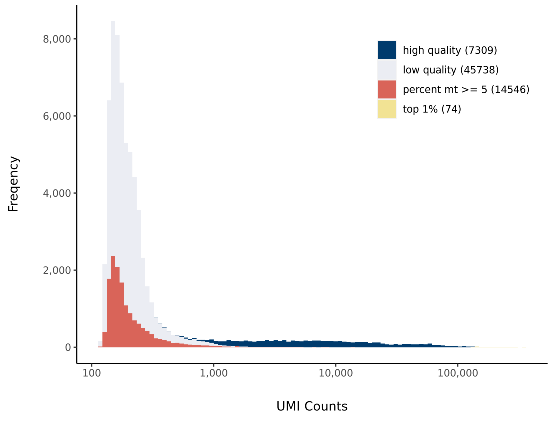


#### Number of Genes Histogram

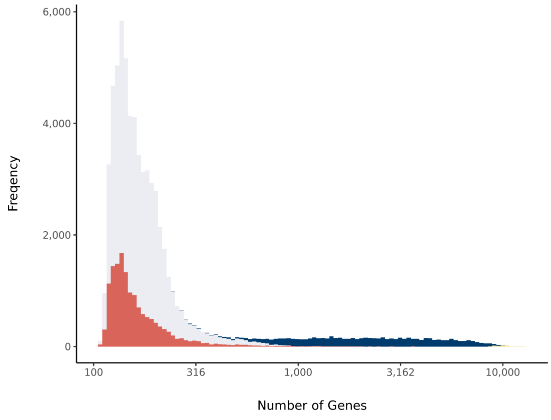


#### Barcode Rank Plot

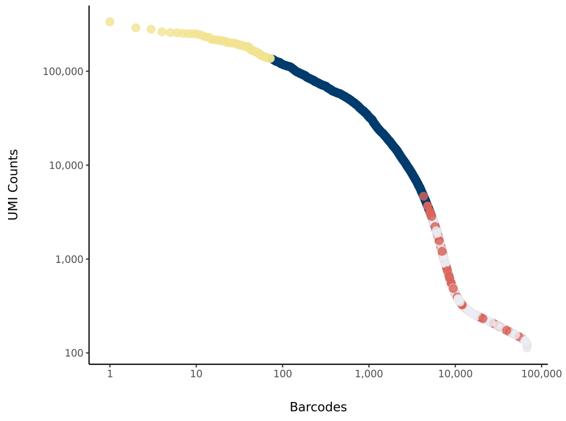

### sc_48 Summary

Processed by COPILOT

Summary Analysis

#### Parameters

| | |
|---|---|
| Iteration of Filtering | 1 |
| Mitochondrial Expression Threshold | 5 % |
| Top High Quality Cell Filtered | 1 % |
| Doublet Removed | Yes |

#### Cell Stats

| | |
|---|---|
| Estimated Number of High Quality Cell | 6,731 |
| High Quality Cell | 16.51 % |
| Total UMI Counts in High Quality Cell | 87,859,537 |
| UMI Counts in High Quality Cell | 81.24 % |
| Median UMI Counts per High Quality Cell | 5,135 |
| Median Genes per High Quality Cell | 1,922 |
| Total Genes Detected in High Quality Cell | 25,534 |
| Cell above Mitochondrial Expression Threshold | 37.03 % |
| Estimated Doublet Rate in High Quality Cell | 5.08 % |

#### Sequencing Stats

| | |
|---|---|
| Number of Reads Processed | 264,836,789 |
| Reads Pseudoaligned | 93.2 % |
| Reads on Whitelist | 96.11 % |
| Total UMI Counts | 108,147,270 |
| Sequencing Technology | 10xv3 |
| Species | Arabidopsis thaliana |
| Transcriptome | TAIR10 |

#### Sample Stats

| | |
|---|---|
| Sample | sc_48 |
| Name | WT BRZ then 8 hour BL |
| Source | Benfey lab |
| Genotype | WT Col-0 |
| Transgene | NA |
| Treatment | BL |
| Age | 7_day |
| Timepoint | 8 |
| Rep | NA |
| Target Cells | 10,000 |
| Date | 2020-02-12 |
| Seq Run | Nolan_6199 (NextSeq); Nolan_6226 (NovaSeq S4) |

#### UMI Counts Histogram

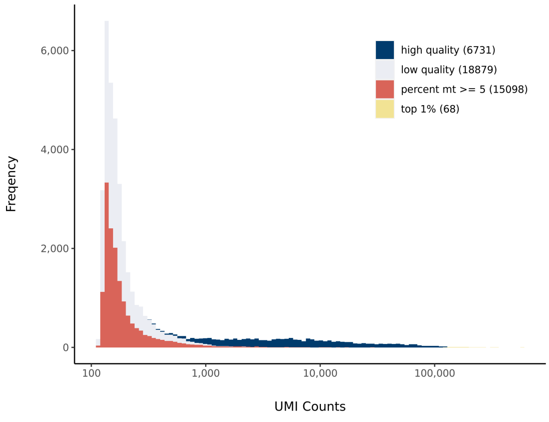


#### Number of Genes Histogram

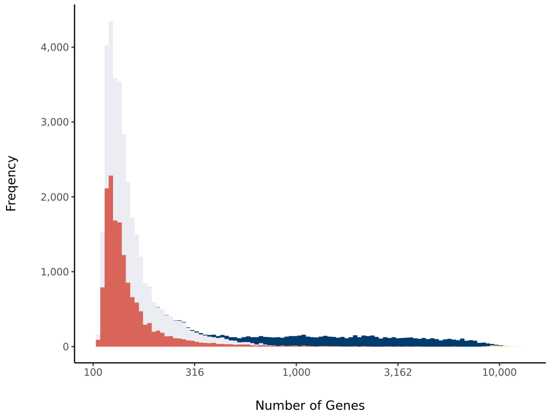


#### Barcode Rank Plot

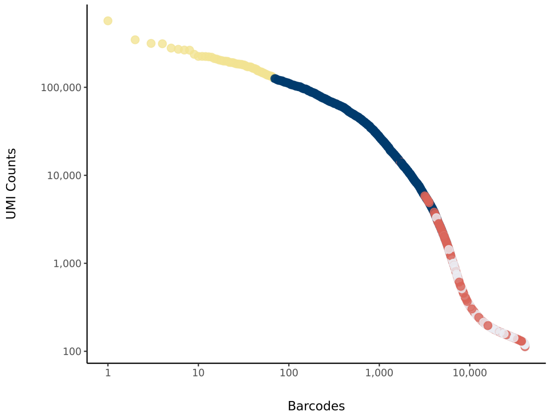

### sc_49 Summary

Processed by COPILOT



#### Parameters

| | |
|---|---|
| Iteration of Filtering | 1 |
| Mitochondrial Expression Threshold | 5 % |
| Top High Quality Cell Filtered | 1 % |
| Doublet Removed | Yes |

#### Cell Stats

| | |
|---|---|
| Estimated Number of High Quality Cell | 5,902 |
| High Quality Cell | 8.42 % |
| Total UMI Counts in High Quality Cell | 94,042,387 |
| UMI Counts in High Quality Cell | 77.97 % |
| Median UMI Counts per High Quality Cell | 7,260.5 |
| Median Genes per High Quality Cell | 2,447 |
| Total Genes Detected in High Quality Cell | 25,254 |
| Cell above Mitochondrial Expression Threshold | 9.31 % |
| Estimated Doublet Rate in High Quality Cell | 4.47 % |

#### Sequencing Stats

| | |
|---|---|
| Number of Reads Processed | 271,798,165 |
| Reads Pseudoaligned | 94 % |
| Reads on Whitelist | 96.31 % |
| Total UMI Counts | 120,612,180 |
| Sequencing Technology | 10xv3 |
| Species | Arabidopsis thaliana |
| Transcriptome | TAIR10 |

#### Sample Stats

| | |
|---|---|
| Sample | sc_49 |
| Name | WT BRZ then 2 hour BL |
| Source | Benfey lab |
| Genotype | WT Col-0 |
| Transgene | NA |
| Treatment | BL |
| Age | 7_day |
| Timepoint | 2 |
| Rep | NA |
| Target Cells | 10,000 |
| Date | 2020-02-12 |
| Seq Run | Nolan_6199 (NextSeq); Nolan_6226 (NovaSeq S4) |

#### UMI Counts Histogram

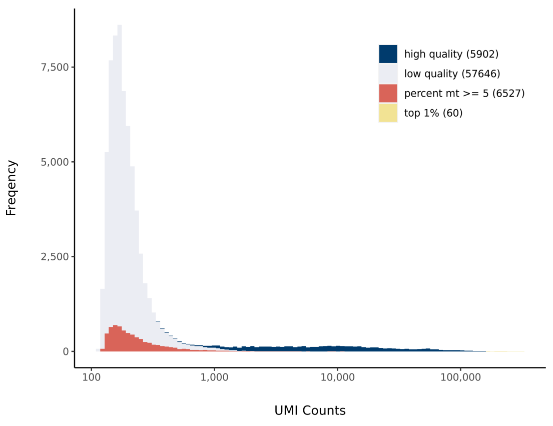


#### Number of Genes Histogram

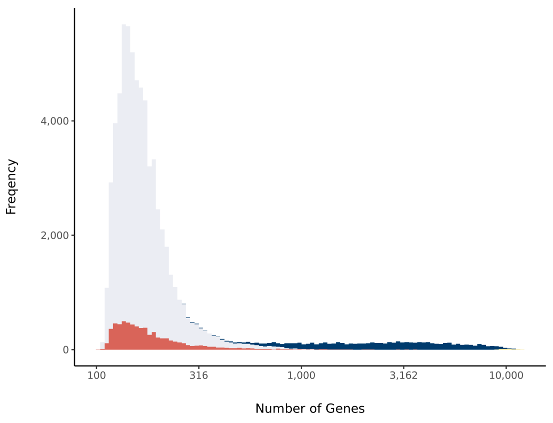


#### Barcode Rank Plot

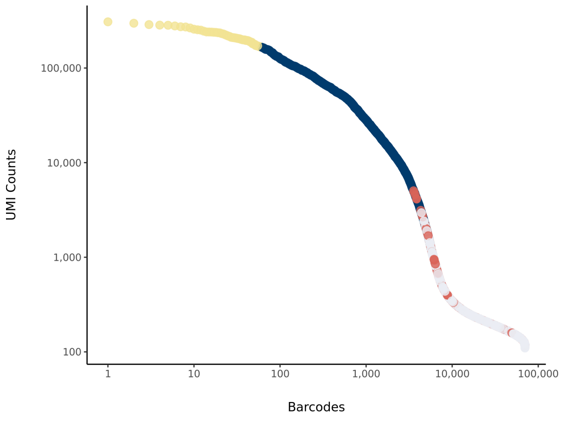

### sc_50 Summary

Processed by COPILOT

Summary | Analysis

#### Parameters

| | |
|---|---|
| Iteration of Filtering | 1 |
| Mitochondrial Expression Threshold | 5 % |
| Top High Quality Cell Filtered | 1 % |
| Doublet Removed | Yes |

#### Cell Stats

| | |
|---|---|
| Estimated Number of High Quality Cell | 5,300 |
| High Quality Cell | 10.46 % |
| Total UMI Counts in High Quality Cell | 92,991,502 |
| UMI Counts in High Quality Cell | 78.59 % |
| Median UMI Counts per High Quality Cell | 6,743.5 |
| Median Genes per High Quality Cell | 2,283.5 |
| Total Genes Detected in High Quality Cell | 24,667 |
| Cell above Mitochondrial Expression Threshold | 7.07 % |
| Estimated Doublet Rate in High Quality Cell | 4.02 % |

#### Sequencing Stats

| | |
|---|---|
| Number of Reads Processed | 261,669,654 |
| Reads Pseudoaligned | 94.3 % |
| Reads on Whitelist | 96.53 % |
| Total UMI Counts | 118,325,019 |
| Sequencing Technology | 10xv3 |
| Species | Arabidopsis thaliana |
| Transcriptome | TAIR10 |

#### Sample Stats

| | |
|---|---|
| Sample | sc_50 |
| Name | WT BRZ control |
| Source | Benfey lab |
| Genotype | WT Col-0 |
| Transgene | NA |
| Treatment | BRZ |
| Age | 7_day |
| Timepoint | 0 |
| Rep | NA |
| Target Cells | 10,000 |
| Date | 2020-02-12 |
| Seq Run | Nolan_6199 (NextSeq); Nolan_6226 (NovaSeq S4) |

#### UMI Counts Histogram

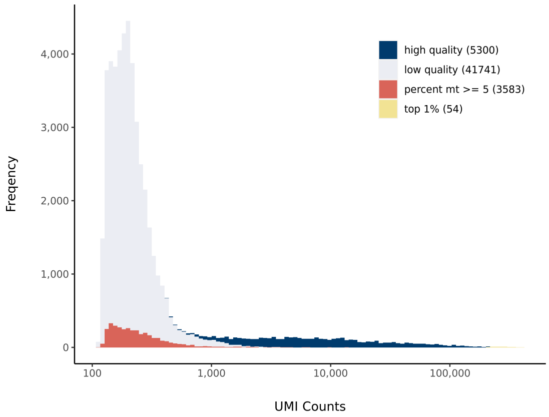


#### Number of Genes Histogram

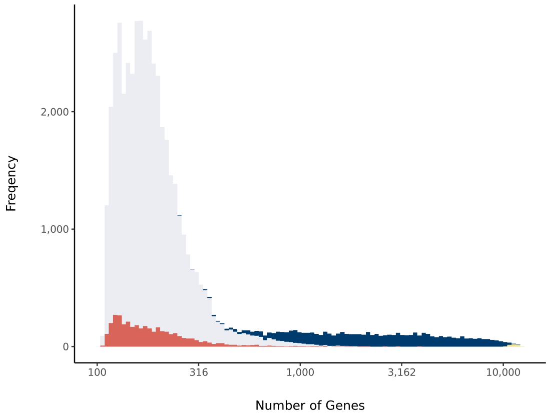


#### Barcode Rank Plot

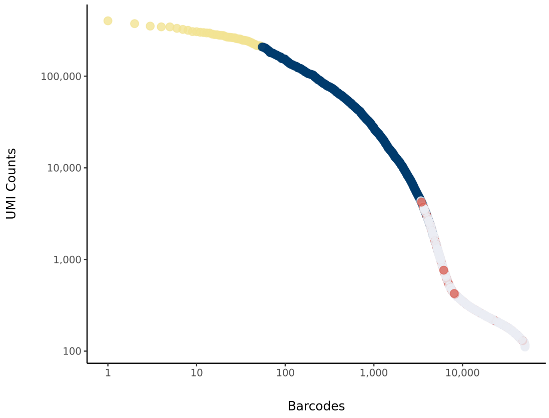

### sc_122 Summary

Processed by COPILOT

Summary Analysis

#### Parameters

| | |
|---|---|
| Iteration of Filtering | 1 |
| Mitochondrial Expression Threshold | 5 % |
| Top High Quality Cell Filtered | 1 % |
| Doublet Removed | Yes |

#### Cell Stats

| | |
|---|---|
| Estimated Number of High Quality Cell | 12,843 |
| High Quality Cell | 14.79 % |
| Total UMI Counts in High Quality Cell | 139,333,645 |
| UMI Counts in High Quality Cell | 58.57 % |
| Median UMI Counts per High Quality Cell | 6,693 |
| Median Genes per High Quality Cell | 2,447 |
| Total Genes Detected in High Quality Cell | 24,728 |
| Cell above Mitochondrial Expression Threshold | 4.85 % |
| Estimated Doublet Rate in High Quality Cell | 9.57 % |

#### Sequencing Stats

| | |
|---|---|
| Number of Reads Processed | 326,482,831 |
| Reads Pseudoaligned | 94.9 % |
| Reads on Whitelist | 96.46 % |
| Total UMI Counts | 237,906,531 |
| Sequencing Technology | 10xv3 |
| Species | Arabidopsis thaliana |
| Transcriptome | TAIR10 |

#### Sample Stats

| | |
|---|---|
| Sample | sc_122 |
| Name | WT |
| Source | Benfey lab |
| Genotype | WT |
| Transgene | NA |
| Treatment | untreated |
| Age | 7_day |
| Timepoint | NA |
| Rep | 1 |
| Target Cells | 10,000 |
| Date | 2021-08-04 |
| Seq Run | Nolan_7191 |

#### UMI Counts Histogram

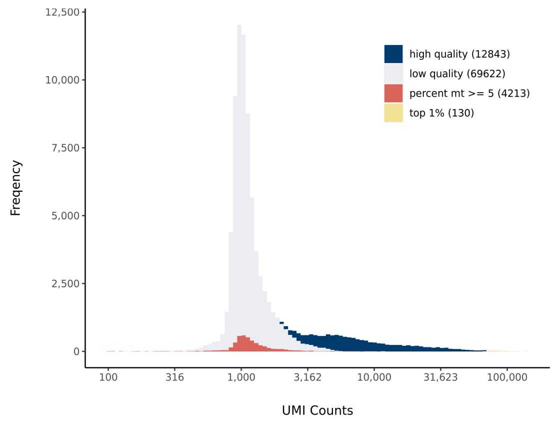


#### Number of Genes Histogram

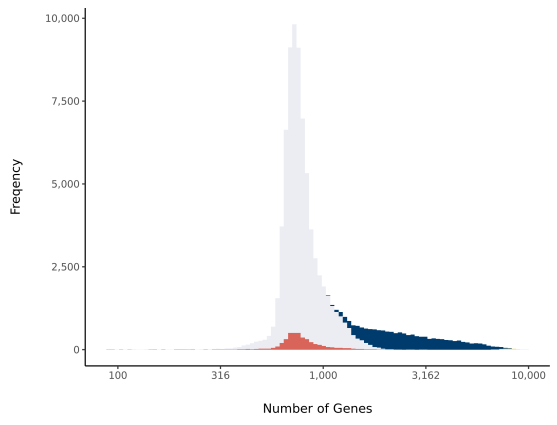


#### Barcode Rank Plot

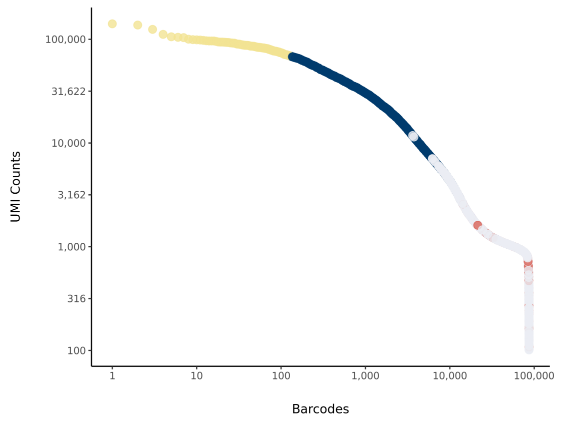

### sc_123 Summary

Processed by COPILOT

Summary Analysis

#### Parameters

| | |
|---|---|
| Iteration of Filtering | 1 |
| Mitochondrial Expression Threshold | 5 % |
| Top High Quality Cell Filtered | 1 % |
| Doublet Removed | Yes |

#### Cell Stats

| | |
|---|---|
| Estimated Number of High Quality Cell | 12,262 |
| High Quality Cell | 14.32 % |
| Total UMI Counts in High Quality Cell | 149,846,646 |
| UMI Counts in High Quality Cell | 56.5 % |
| Median UMI Counts per High Quality Cell | 7,540 |
| Median Genes per High Quality Cell | 2,537 |
| Total Genes Detected in High Quality Cell | 24,959 |
| Cell above Mitochondrial Expression Threshold | 1.96 % |
| Estimated Doublet Rate in High Quality Cell | 9.14 % |

#### Sequencing Stats

| | |
|---|---|
| Number of Reads Processed | 376,362,931 |
| Reads Pseudoaligned | 95.4 % |
| Reads on Whitelist | 96.64 % |
| Total UMI Counts | 265,202,057 |
| Sequencing Technology | 10xv3 |
| Species | Arabidopsis thaliana |
| Transcriptome | TAIR10 |

#### Sample Stats

| | |
|---|---|
| Sample | sc_123 |
| Name | gtl1-1 |
| Source | Benfey lab |
| Genotype | gtl1-1 |
| Transgene | NA |
| Treatment | untreated |
| Age | 7_day |
| Timepoint | NA |
| Rep | 1 |
| Target Cells | 10,000 |
| Date | 2021-08-04 |
| Seq Run | Nolan_7191 |

#### UMI Counts Histogram

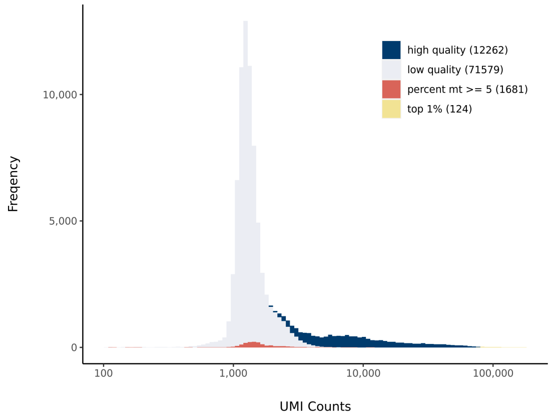


#### Number of Genes Histogram

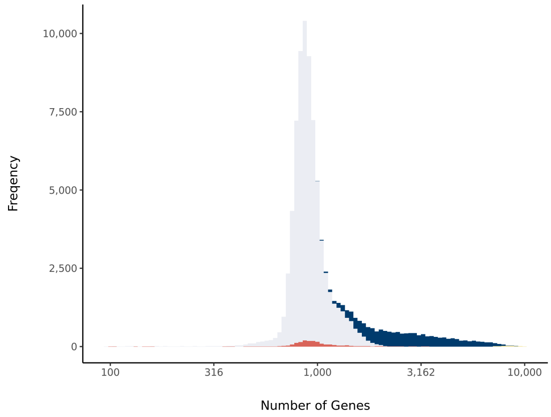


#### Barcode Rank Plot

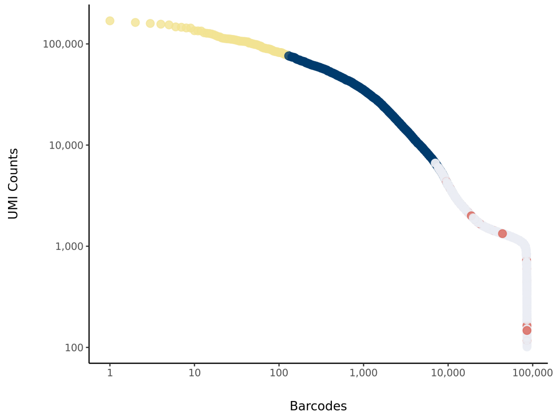

### sc_124 Summary

Processed by COPILOT

Summary Analysis

#### Parameters

| | |
|---|---|
| Iteration of Filtering | 1 |
| Mitochondrial Expression Threshold | 5 % |
| Top High Quality Cell Filtered | 1 % |
| Doublet Removed | Yes |

#### Cell Stats

| | |
|---|---|
| Estimated Number of High Quality Cell | 9,772 |
| High Quality Cell | 11.2 % |
| Total UMI Counts in High Quality Cell | 99,745,666 |
| UMI Counts in High Quality Cell | 52.25 % |
| Median UMI Counts per High Quality Cell | 6,756 |
| Median Genes per High Quality Cell | 2,286.5 |
| Total Genes Detected in High Quality Cell | 24,593 |
| Cell above Mitochondrial Expression Threshold | 14.1 % |
| Estimated Doublet Rate in High Quality Cell | 7.31 % |

#### Sequencing Stats

| | |
|---|---|
| Number of Reads Processed | 283,737,048 |
| Reads Pseudoaligned | 94.4 % |
| Reads on Whitelist | 96.32 % |
| Total UMI Counts | 190,889,781 |
| Sequencing Technology | 10xv3 |
| Species | Arabidopsis thaliana |
| Transcriptome | TAIR10 |

#### Sample Stats

| | |
|---|---|
| Sample | sc_124 |
| Name | df1-1 |
| Source | Benfey lab |
| Genotype | df1-1 |
| Transgene | NA |
| Treatment | untreated |
| Age | 7_day |
| Timepoint | NA |
| Rep | 1 |
| Target Cells | 10,000 |
| Date | 2021-08-04 |
| Seq Run | Nolan_7191 |

#### UMI Counts Histogram

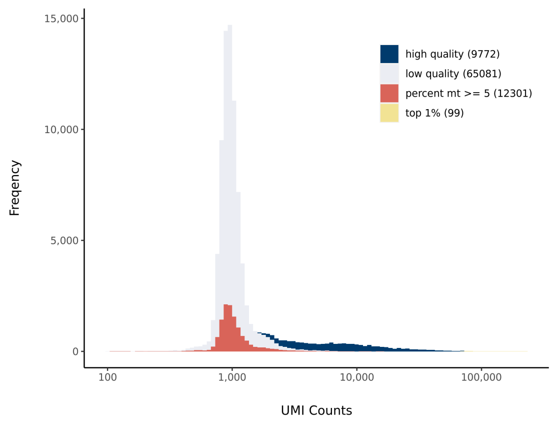


#### Number of Genes Histogram

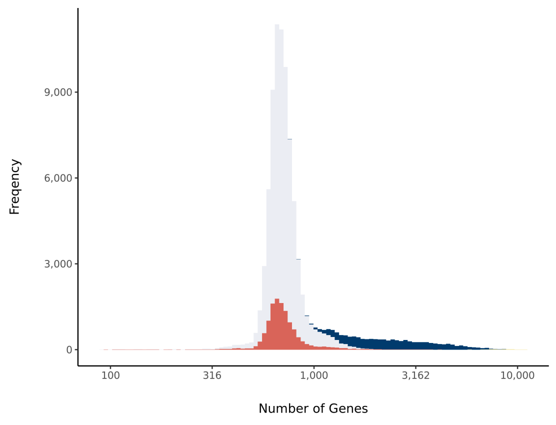


#### Barcode Rank Plot

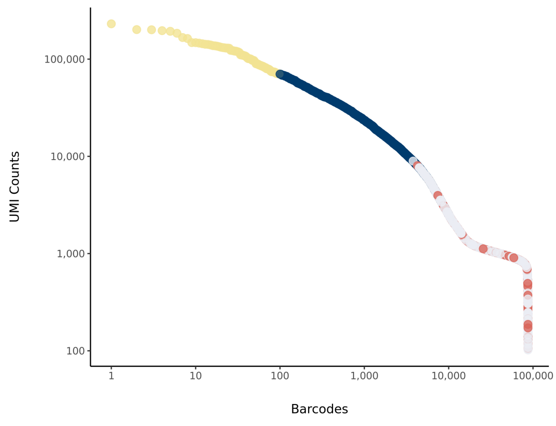

### sc_125 Summary

Processed by COPILOT

Summary | Analysis

#### Parameters

| | |
|---|---|
| Iteration of Filtering | 1 |
| Mitochondrial Expression Threshold | 5 % |
| Top High Quality Cell Filtered | 1 % |
| Doublet Removed | Yes |

#### Cell Stats

| | |
|---|---|
| Estimated Number of High Quality Cell | 10,899 |
| High Quality Cell | 12.53 % |
| Total UMI Counts in High Quality Cell | 146,785,190 |
| UMI Counts in High Quality Cell | 58.97 % |
| Median UMI Counts per High Quality Cell | 9,000 |
| Median Genes per High Quality Cell | 2,804 |
| Total Genes Detected in High Quality Cell | 25,295 |
| Cell above Mitochondrial Expression Threshold | 5.96 % |
| Estimated Doublet Rate in High Quality Cell | 8.14 % |

#### Sequencing Stats

| | |
|---|---|
| Number of Reads Processed | 366,207,321 |
| Reads Pseudoaligned | 95.4 % |
| Reads on Whitelist | 96.6 % |
| Total UMI Counts | 248,908,711 |
| Sequencing Technology | 10xv3 |
| Species | Arabidopsis thaliana |
| Transcriptome | TAIR10 |

#### Sample Stats

| | |
|---|---|
| Sample | sc_125 |
| Name | gtl1-1 df1-1 |
| Source | Benfey lab |
| Genotype | gtl1-1 df1-1 |
| Transgene | NA |
| Treatment | untreated |
| Age | 7_day |
| Timepoint | NA |
| Rep | 1 |
| Target Cells | 10,000 |
| Date | 2021-08-04 |
| Seq Run | Nolan_7191 |

#### UMI Counts Histogram

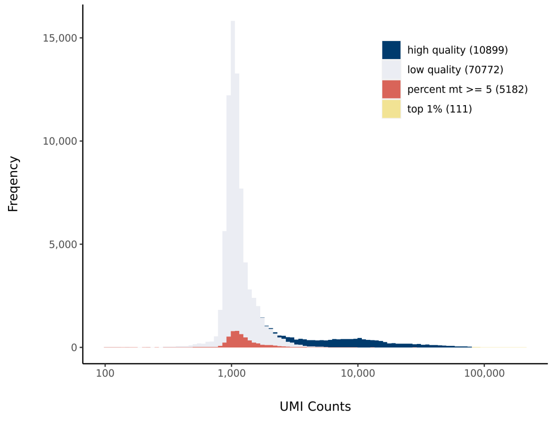


#### Number of Genes Histogram

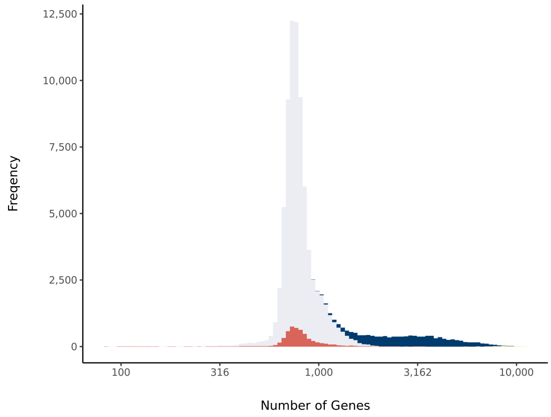


#### Barcode Rank Plot

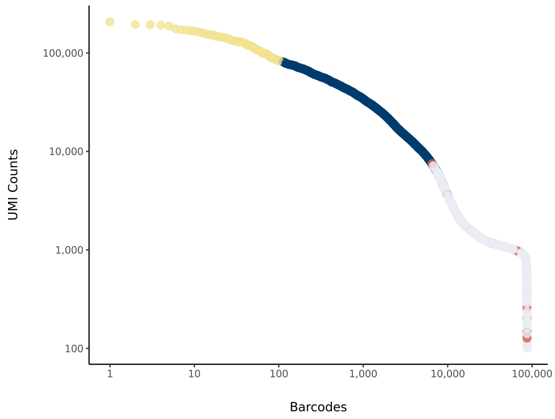

### sc_126 Summary

Processed by COPILOT

Summary Analysis

#### Parameters

| | |
|---|---|
| Iteration of Filtering | 1 |
| Mitochondrial Expression Threshold | 5 % |
| Top High Quality Cell Filtered | 1 % |
| Doublet Removed | Yes |

#### Cell Stats

| | |
|---|---|
| Estimated Number of High Quality Cell | 7,741 |
| High Quality Cell | 8.91 % |
| Total UMI Counts in High Quality Cell | 128,314,754 |
| UMI Counts in High Quality Cell | 52.16 % |
| Median UMI Counts per High Quality Cell | 10,111 |
| Median Genes per High Quality Cell | 2,888 |
| Total Genes Detected in High Quality Cell | 24,631 |
| Cell above Mitochondrial Expression Threshold | 2.3 % |
| Estimated Doublet Rate in High Quality Cell | 5.82 % |

#### Sequencing Stats

| | |
|---|---|
| Number of Reads Processed | 375,494,818 |
| Reads Pseudoaligned | 95.2 % |
| Reads on Whitelist | 96.49 % |
| Total UMI Counts | 246,007,796 |
| Sequencing Technology | 10xv3 |
| Species | Arabidopsis thaliana |
| Transcriptome | TAIR10 |

#### Sample Stats

| | |
|---|---|
| Sample | sc_126 |
| Name | WT |
| Source | Benfey lab |
| Genotype | WT |
| Transgene | NA |
| Treatment | untreated |
| Age | 7_day |
| Timepoint | NA |
| Rep | 2 |
| Target Cells | 10,000 |
| Date | 2021-08-04 |
| Seq Run | Nolan_7191 |

#### UMI Counts Histogram

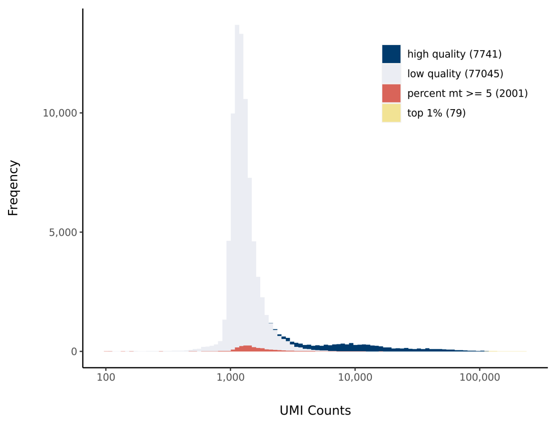


#### Number of Genes Histogram

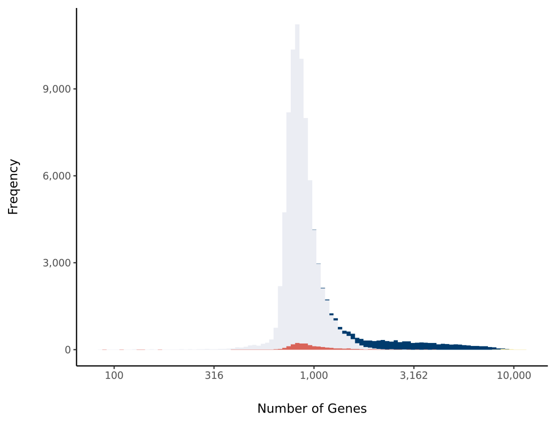


#### Barcode Rank Plot

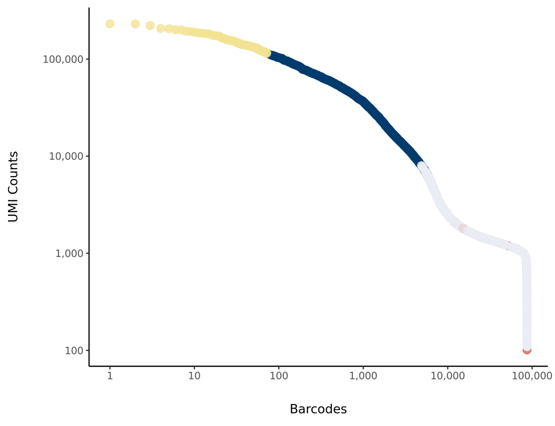

### sc_127 Summary

Processed by COPILOT

Summary Analysis

#### Parameters

| | |
|---|---|
| Iteration of Filtering | 1 |
| Mitochondrial Expression Threshold | 5 % |
| Top High Quality Cell Filtered | 1 % |
| Doublet Removed | Yes |

#### Cell Stats

| | |
|---|---|
| Estimated Number of High Quality Cell | 12,644 |
| High Quality Cell | 14.26 % |
| Total UMI Counts in High Quality Cell | 122,023,275 |
| UMI Counts in High Quality Cell | 52.99 % |
| Median UMI Counts per High Quality Cell | 6,501 |
| Median Genes per High Quality Cell | 2,354 |
| Total Genes Detected in High Quality Cell | 24,929 |
| Cell above Mitochondrial Expression Threshold | 2.89 % |
| Estimated Doublet Rate in High Quality Cell | 9.42 % |

#### Sequencing Stats

| | |
|---|---|
| Number of Reads Processed | 322,509,318 |
| Reads Pseudoaligned | 95.1 % |
| Reads on Whitelist | 96.52 % |
| Total UMI Counts | 230,263,769 |
| Sequencing Technology | 10xv3 |
| Species | Arabidopsis thaliana |
| Transcriptome | TAIR10 |

#### Sample Stats

| | |
|---|---|
| Sample | sc_127 |
| Name | gtl1-1 |
| Source | Benfey lab |
| Genotype | gtl1-1 |
| Transgene | NA |
| Treatment | untreated |
| Age | 7_day |
| Timepoint | NA |
| Rep | 2 |
| Target Cells | 10,000 |
| Date | 2021-08-04 |
| Seq Run | Nolan_7191 |

#### UMI Counts Histogram

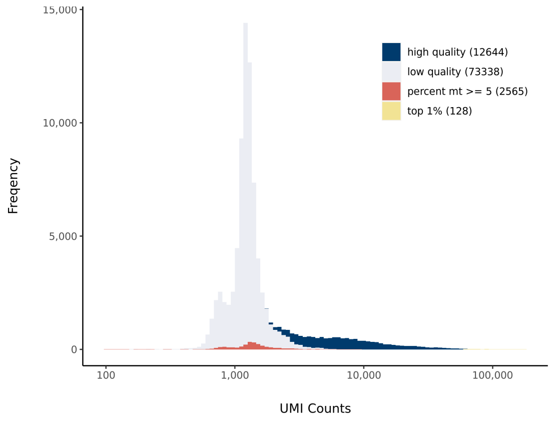


#### Number of Genes Histogram

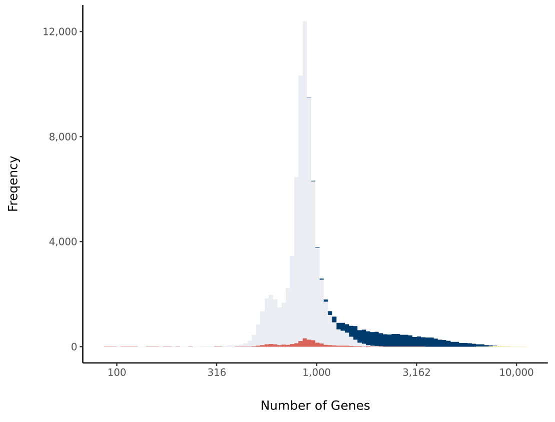


#### Barcode Rank Plot

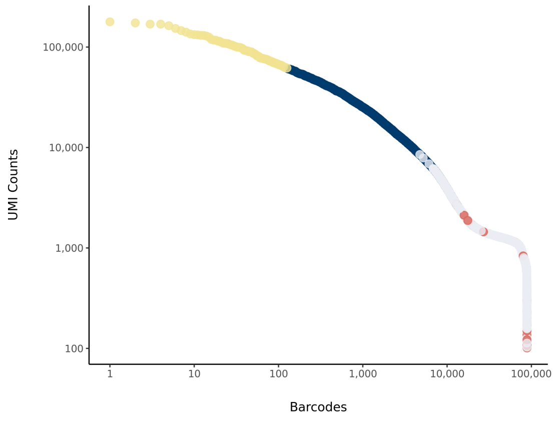

### sc_128 Summary

Processed by COPILOT

Summary Analysis

#### Parameters

| | |
|---|---|
| Iteration of Filtering | 1 |
| Mitochondrial Expression Threshold | 5 % |
| Top High Quality Cell Filtered | 1 % |
| Doublet Removed | Yes |

#### Cell Stats

| | |
|---|---|
| Estimated Number of High Quality Cell | 6,988 |
| High Quality Cell | 8.03 % |
| Total UMI Counts in High Quality Cell | 134,268,224 |
| UMI Counts in High Quality Cell | 59.94 % |
| Median UMI Counts per High Quality Cell | 11,606 |
| Median Genes per High Quality Cell | 3,062.5 |
| Total Genes Detected in High Quality Cell | 24,917 |
| Cell above Mitochondrial Expression Threshold | 6.51 % |
| Estimated Doublet Rate in High Quality Cell | 5.26 % |

#### Sequencing Stats

| | |
|---|---|
| Number of Reads Processed | 365,390,040 |
| Reads Pseudoaligned | 93.9 % |
| Reads on Whitelist | 96.55 % |
| Total UMI Counts | 223,985,991 |
| Sequencing Technology | 10xv3 |
| Species | Arabidopsis thaliana |
| Transcriptome | TAIR10 |

#### Sample Stats

| | |
|---|---|
| Sample | sc_128 |
| Name | df1-1 |
| Source | Benfey lab |
| Genotype | df1-1 |
| Transgene | NA |
| Treatment | untreated |
| Age | 7_day |
| Timepoint | NA |
| Rep | 2 |
| Target Cells | 10,000 |
| Date | 2021-08-04 |
| Seq Run | Nolan_7191 |

#### UMI Counts Histogram

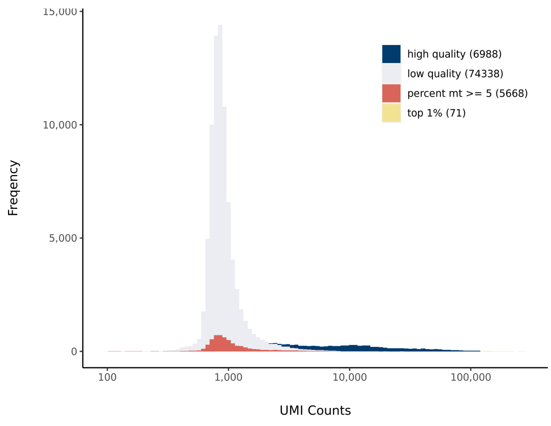


#### Number of Genes Histogram

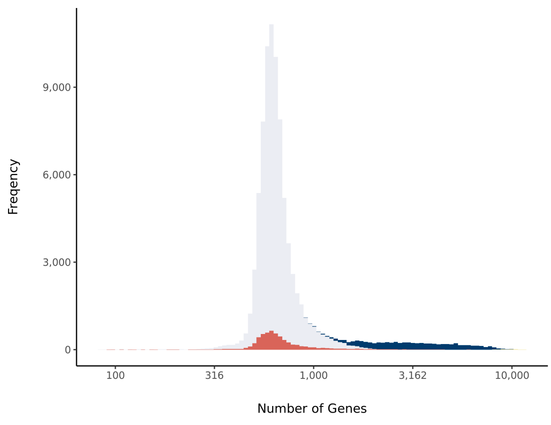


#### Barcode Rank Plot

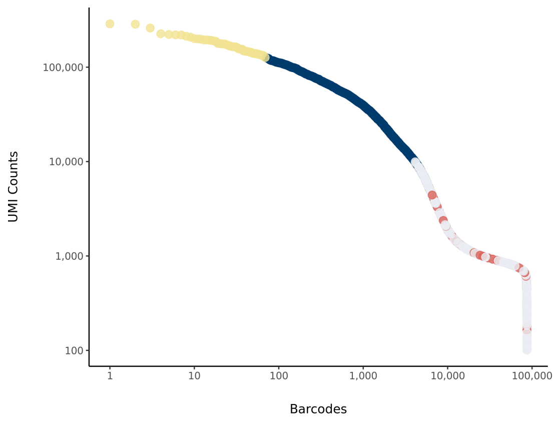

### sc_129 Summary

Processed by COPILOT

Summary Analysis

#### Parameters

| | |
|---|---|
| Iteration of Filtering | 1 |
| Mitochondrial Expression Threshold | 5 % |
| Top High Quality Cell Filtered | 1 % |
| Doublet Removed | Yes |

#### Cell Stats

| | |
|---|---|
| Estimated Number of High Quality Cell | 8,117 |
| High Quality Cell | 9.12 % |
| Total UMI Counts in High Quality Cell | 139,829,922 |
| UMI Counts in High Quality Cell | 57.84 % |
| Median UMI Counts per High Quality Cell | 10,808 |
| Median Genes per High Quality Cell | 3,104 |
| Total Genes Detected in High Quality Cell | 25,180 |
| Cell above Mitochondrial Expression Threshold | 1.54 % |
| Estimated Doublet Rate in High Quality Cell | 6.1 % |

#### Sequencing Stats

| | |
|---|---|
| Number of Reads Processed | 375,752,207 |
| Reads Pseudoaligned | 94.8 % |
| Reads on Whitelist | 96.56 % |
| Total UMI Counts | 241,760,841 |
| Sequencing Technology | 10xv3 |
| Species | Arabidopsis thaliana |
| Transcriptome | TAIR10 |

#### Sample Stats

| | |
|---|---|
| Sample | sc_129 |
| Name | gtl1-1 df1-1 |
| Source | Benfey lab |
| Genotype | gtl1-1 df1-1 |
| Transgene | NA |
| Treatment | untreated |
| Age | 7_day |
| Timepoint | NA |
| Rep | 2 |
| Target Cells | 10,000 |
| Date | 2021-08-04 |
| Seq Run | Nolan_7191 |

#### UMI Counts Histogram

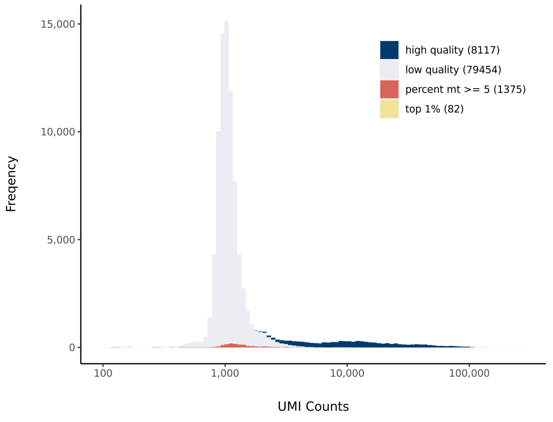


#### Number of Genes Histogram

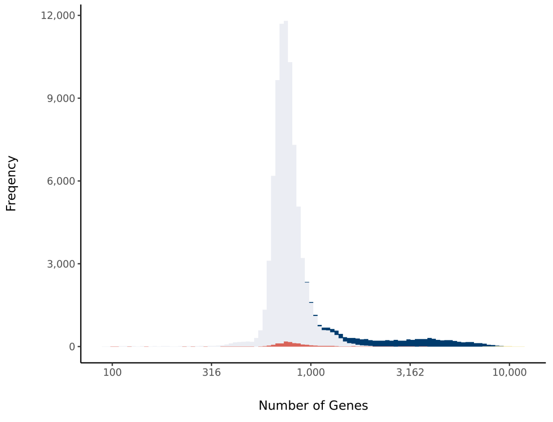


#### Barcode Rank Plot

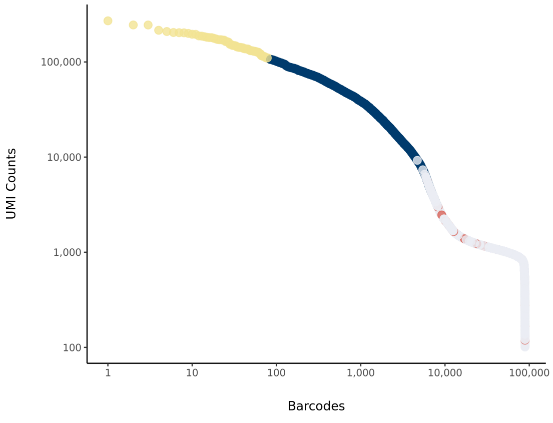

### sc_130 Summary

Processed by COPILOT

Summary | Analysis

#### Parameters

| | |
|---|---|
| Iteration of Filtering | 1 |
| Mitochondrial Expression Threshold | 5 % |
| Top High Quality Cell Filtered | 1 % |
| Doublet Removed | Yes |

#### Cell Stats

| | |
|---|---|
| Estimated Number of High Quality Cell | 6,953 |
| High Quality Cell | 7.66 % |
| Total UMI Counts in High Quality Cell | 136,955,357 |
| UMI Counts in High Quality Cell | 41.53 % |
| Median UMI Counts per High Quality Cell | 10,855 |
| Median Genes per High Quality Cell | 3,052 |
| Total Genes Detected in High Quality Cell | 24,704 |
| Cell above Mitochondrial Expression Threshold | 0.55 % |
| Estimated Doublet Rate in High Quality Cell | 5.24 % |

#### Sequencing Stats

| | |
|---|---|
| Number of Reads Processed | 529,797,376 |
| Reads Pseudoaligned | 95.7 % |
| Reads on Whitelist | 96.84 % |
| Total UMI Counts | 329,785,001 |
| Sequencing Technology | 10xv3 |
| Species | Arabidopsis thaliana |
| Transcriptome | TAIR10 |

#### Sample Stats

| | |
|---|---|
| Sample | sc_130 |
| Name | WT |
| Source | Benfey lab |
| Genotype | WT |
| Transgene | NA |
| Treatment | untreated |
| Age | 7_day |
| Timepoint | NA |
| Rep | 1 |
| Target Cells | 10,000 |
| Date | 2021-10-01 |
| Seq Run | Nolan_7335 |

#### UMI Counts Histogram

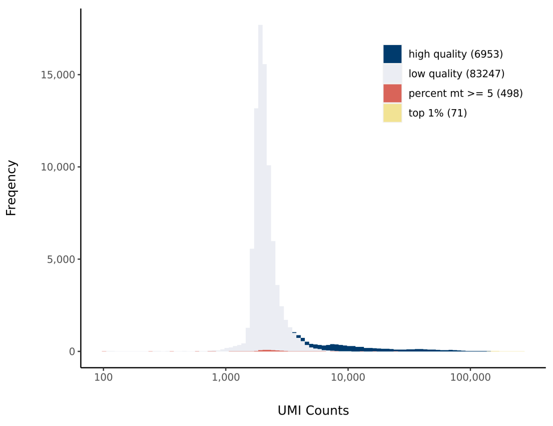


#### Number of Genes Histogram

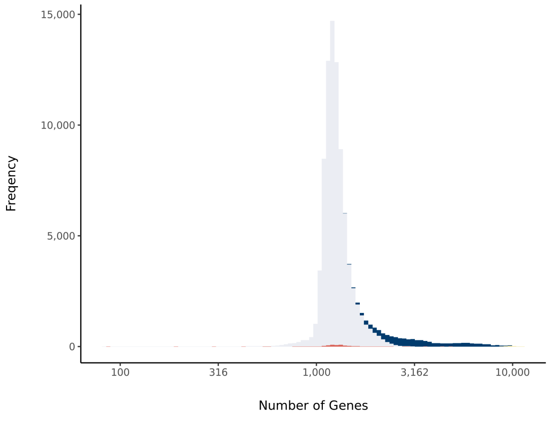


#### Barcode Rank Plot

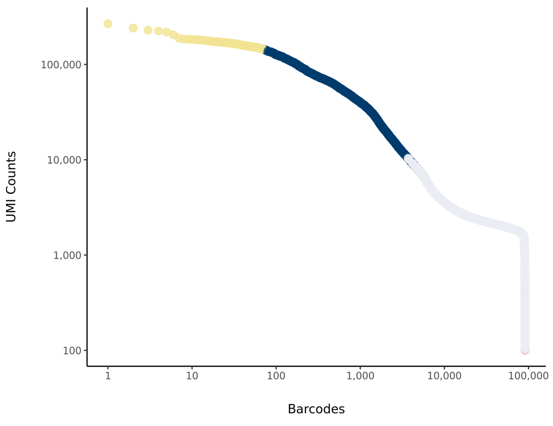

### sc_131 Summary

Processed by COPILOT



#### Parameters

| | |
|---|---|
| Iteration of Filtering | 1 |
| Mitochondrial Expression Threshold | 5 % |
| Top High Quality Cell Filtered | 1 % |
| Doublet Removed | Yes |

#### Cell Stats

| | |
|---|---|
| Estimated Number of High Quality Cell | 8,115 |
| High Quality Cell | 9.33 % |
| Total UMI Counts in High Quality Cell | 130,890,333 |
| UMI Counts in High Quality Cell | 41.96 % |
| Median UMI Counts per High Quality Cell | 9,696 |
| Median Genes per High Quality Cell | 3,184 |
| Total Genes Detected in High Quality Cell | 25,026 |
| Cell above Mitochondrial Expression Threshold | 2.39 % |
| Estimated Doublet Rate in High Quality Cell | 6.09 % |

#### Sequencing Stats

| | |
|---|---|
| Number of Reads Processed | 462,710,489 |
| Reads Pseudoaligned | 95.1 % |
| Reads on Whitelist | 96.74 % |
| Total UMI Counts | 311,907,742 |
| Sequencing Technology | 10xv3 |
| Species | Arabidopsis thaliana |
| Transcriptome | TAIR10 |

#### Sample Stats

| | |
|---|---|
| Sample | sc_131 |
| Name | bri1-T |
| Source | Benfey lab |
| Genotype | bri1-116 brl1 brl3 (bri1-T) |
| Transgene | WOX5-GFP |
| Treatment | untreated |
| Age | 7_day |
| Timepoint | NA |
| Rep | 1 |
| Target Cells | 10,000 |
| Date | 2021-10-01 |
| Seq Run | Nolan_7335 |

#### UMI Counts Histogram

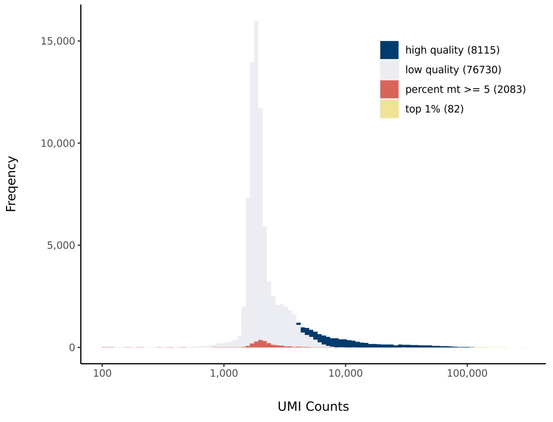


#### Number of Genes Histogram

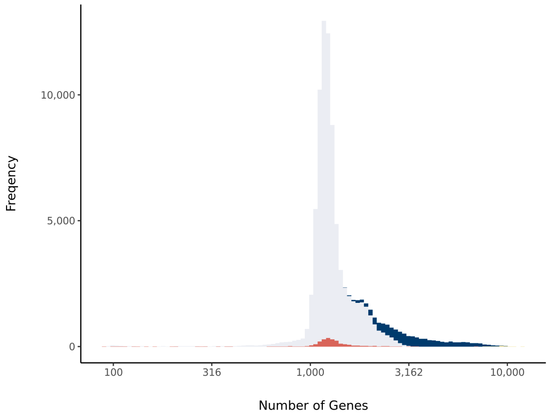


#### Barcode Rank Plot

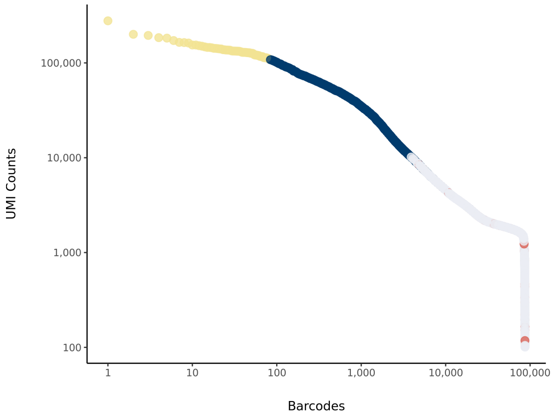

### sc_133 Summary

Processed by COPILOT

Summary | Analysis

#### Parameters

| | |
|---|---|
| Iteration of Filtering | 1 |
| Mitochondrial Expression Threshold | 5 % |
| Top High Quality Cell Filtered | 1 % |
| Doublet Removed | Yes |

#### Cell Stats

| | |
|---|---|
| Estimated Number of High Quality Cell | 6,272 |
| High Quality Cell | 7.21 % |
| Total UMI Counts in High Quality Cell | 131,132,934 |
| UMI Counts in High Quality Cell | 47.03 % |
| Median UMI Counts per High Quality Cell | 11,499 |
| Median Genes per High Quality Cell | 3,427 |
| Total Genes Detected in High Quality Cell | 25,290 |
| Cell above Mitochondrial Expression Threshold | 5.96 % |
| Estimated Doublet Rate in High Quality Cell | 4.74 % |

#### Sequencing Stats

| | |
|---|---|
| Number of Reads Processed | 470,314,179 |
| Reads Pseudoaligned | 94.7 % |
| Reads on Whitelist | 96.75 % |
| Total UMI Counts | 278,806,311 |
| Sequencing Technology | 10xv3 |
| Species | Arabidopsis thaliana |
| Transcriptome | TAIR10 |

#### Sample Stats

| | |
|---|---|
| Sample | sc_133 |
| Name | pGL2:BRI1-GFP/bri1-T |
| Source | Benfey lab |
| Genotype | pGL2:BRI1-GFP/bri1-T |
| Transgene | pGL2:BRI1-GFP |
| Treatment | untreated |
| Age | 7_day |
| Timepoint | NA |
| Rep | 1 |
| Target Cells | 10,000 |
| Date | 2021-10-01 |
| Seq Run | Nolan_7335 |

#### UMI Counts Histogram

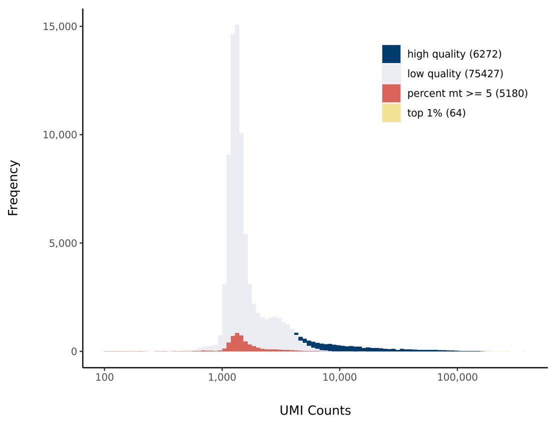


#### Number of Genes Histogram

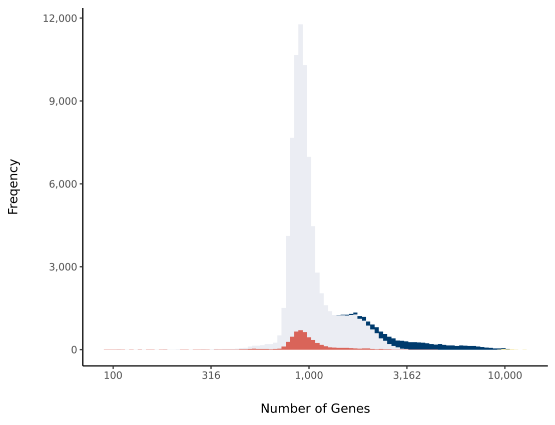


#### Barcode Rank Plot

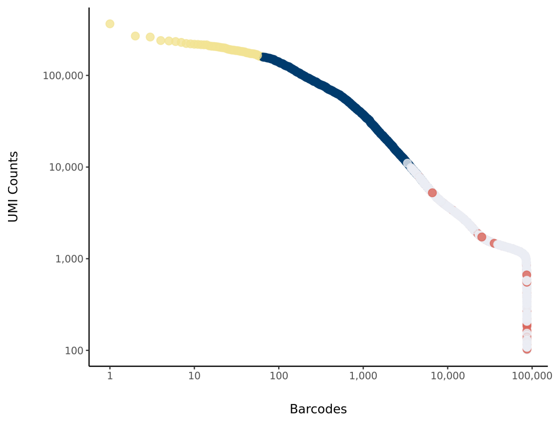

### sc_134 Summary

Processed by COPILOT



#### Parameters

| | |
|---|---|
| Iteration of Filtering | 1 |
| Mitochondrial Expression Threshold | 5 % |
| Top High Quality Cell Filtered | 1 % |
| Doublet Removed | Yes |

#### Cell Stats

| | |
|---|---|
| Estimated Number of High Quality Cell | 8,257 |
| High Quality Cell | 9.33 % |
| Total UMI Counts in High Quality Cell | 159,499,874 |
| UMI Counts in High Quality Cell | 53.76 % |
| Median UMI Counts per High Quality Cell | 10,757 |
| Median Genes per High Quality Cell | 3,062 |
| Total Genes Detected in High Quality Cell | 24,979 |
| Cell above Mitochondrial Expression Threshold | 1.3 % |
| Estimated Doublet Rate in High Quality Cell | 6.2 % |

#### Sequencing Stats

| | |
|---|---|
| Number of Reads Processed | 455,487,382 |
| Reads Pseudoaligned | 95.8 % |
| Reads on Whitelist | 96.86 % |
| Total UMI Counts | 296,703,166 |
| Sequencing Technology | 10xv3 |
| Species | Arabidopsis thaliana |
| Transcriptome | TAIR10 |

#### Sample Stats

| | |
|---|---|
| Sample | sc_134 |
| Name | WT |
| Source | Benfey lab |
| Genotype | WT |
| Transgene | NA |
| Treatment | untreated |
| Age | 7_day |
| Timepoint | NA |
| Rep | 2 |
| Target Cells | 10,000 |
| Date | 2021-10-01 |
| Seq Run | Nolan_7335 |

#### UMI Counts Histogram

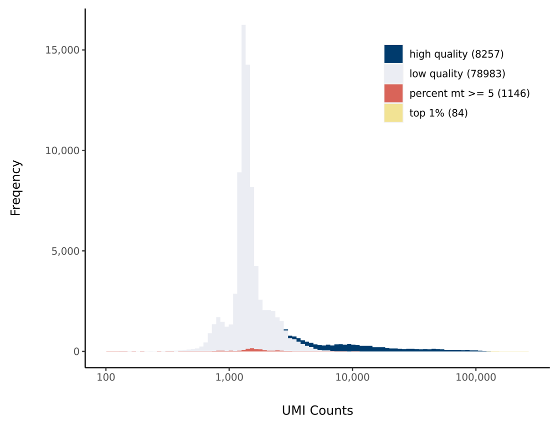


#### Number of Genes Histogram

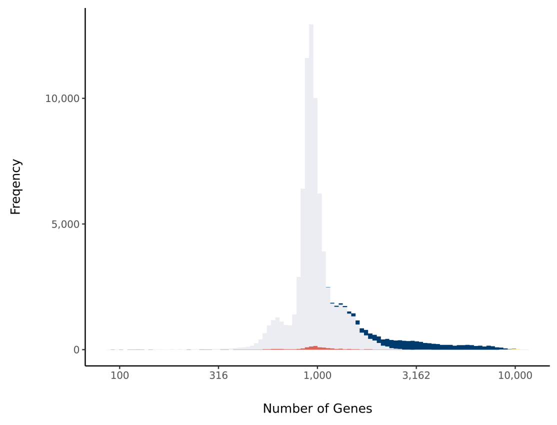


#### Barcode Rank Plot

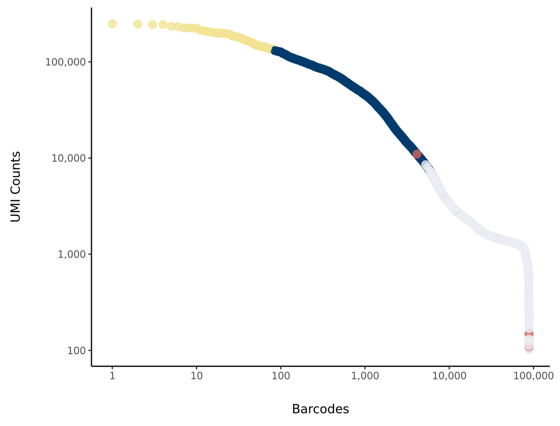

### sc_135 Summary

Processed by COPILOT

Summary Analysis

#### Parameters

| | |
|---|---|
| Iteration of Filtering | 1 |
| Mitochondrial Expression Threshold | 5 % |
| Top High Quality Cell Filtered | 1 % |
| Doublet Removed | Yes |

#### Cell Stats

| | |
|---|---|
| Estimated Number of High Quality Cell | 5,236 |
| High Quality Cell | 6.12 % |
| Total UMI Counts in High Quality Cell | 105,318,949 |
| UMI Counts in High Quality Cell | 37.22 % |
| Median UMI Counts per High Quality Cell | 12,117 |
| Median Genes per High Quality Cell | 3,592 |
| Total Genes Detected in High Quality Cell | 24,793 |
| Cell above Mitochondrial Expression Threshold | 3.17 % |
| Estimated Doublet Rate in High Quality Cell | 3.98 % |

#### Sequencing Stats

| | |
|---|---|
| Number of Reads Processed | 436,804,534 |
| Reads Pseudoaligned | 95 % |
| Reads on Whitelist | 96.81 % |
| Total UMI Counts | 282,987,723 |
| Sequencing Technology | 10xv3 |
| Species | Arabidopsis thaliana |
| Transcriptome | TAIR10 |

#### Sample Stats

| | |
|---|---|
| Sample | sc_135 |
| Name | bri1-T |
| Source | Benfey lab |
| Genotype | bri1-116 brl1 brl3 (bri1-T) |
| Transgene | WOX5-GFP |
| Treatment | untreated |
| Age | 7_day |
| Timepoint | NA |
| Rep | 2 |
| Target Cells | 10,000 |
| Date | 2021-10-01 |
| Seq Run | Nolan_7335 |

#### UMI Counts Histogram

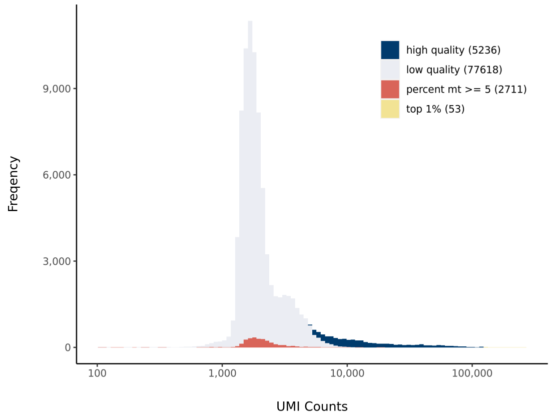


#### Number of Genes Histogram

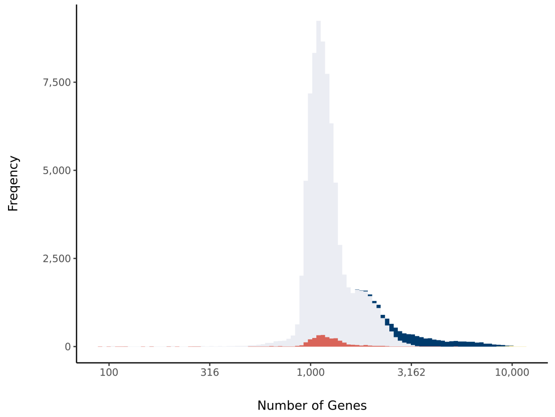


#### Barcode Rank Plot

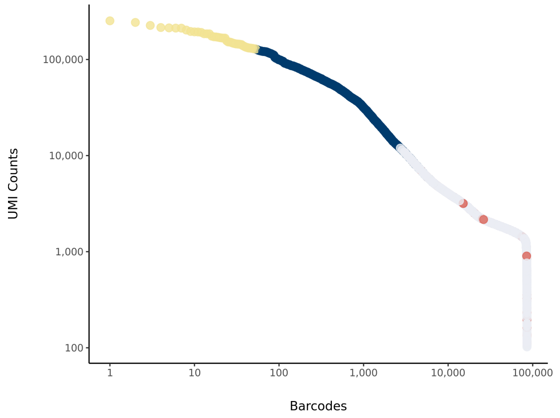

### sc_137 Summary

Processed by COPILOT

Summary | Analysis

#### Parameters

| | |
|---|---|
| Iteration of Filtering | 1 |
| Mitochondrial Expression Threshold | 5 % |
| Top High Quality Cell Filtered | 1 % |
| Doublet Removed | Yes |

#### Cell Stats

| | |
|---|---|
| Estimated Number of High Quality Cell | 1,933 |
| High Quality Cell | 4.29 % |
| Total UMI Counts in High Quality Cell | 71,399,623 |
| UMI Counts in High Quality Cell | 48.41 % |
| Median UMI Counts per High Quality Cell | 18,761 |
| Median Genes per High Quality Cell | 4,147 |
| Total Genes Detected in High Quality Cell | 25,031 |
| Cell above Mitochondrial Expression Threshold | 81.03 % |
| Estimated Doublet Rate in High Quality Cell | 1.55 % |

#### Sequencing Stats

| | |
|---|---|
| Number of Reads Processed | 669,954,882 |
| Reads Pseudoaligned | 90.4 % |
| Reads on Whitelist | 96.3 % |
| Total UMI Counts | 147,484,334 |
| Sequencing Technology | 10xv3 |
| Species | Arabidopsis thaliana |
| Transcriptome | TAIR10 |

#### Sample Stats

| | |
|---|---|
| Sample | sc_137 |
| Name | pGL2:BRI1-GFP/bri1-T |
| Source | Benfey lab |
| Genotype | pGL2:BRI1-GFP/bri1-T |
| Transgene | pGL2:BRI1-GFP |
| Treatment | untreated |
| Age | 7_day |
| Timepoint | NA |
| Rep | 2 |
| Target Cells | 10,000 |
| Date | 2021-10-01 |
| Seq Run | Nolan_7335 |

#### UMI Counts Histogram

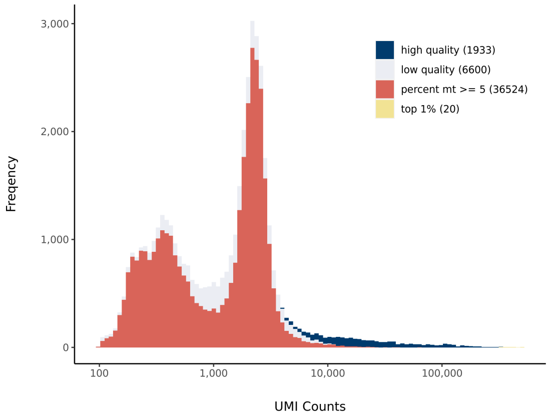


#### Number of Genes Histogram

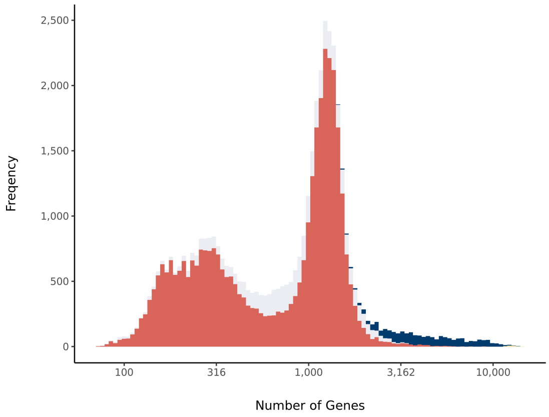


#### Barcode Rank Plot

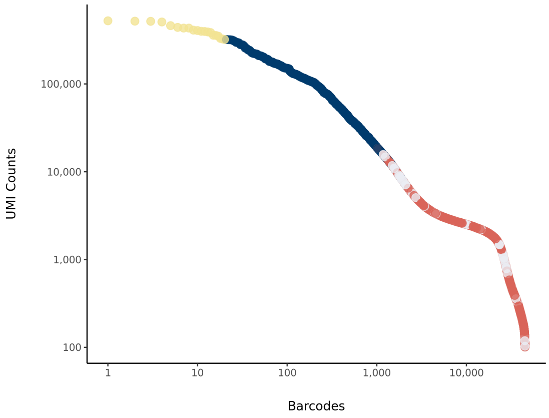
